# Chiral methionine oxidation reagents reveal stereospecific proteome modifications

**DOI:** 10.64898/2026.03.24.713977

**Authors:** Angel Gonzalez-Valero, Annika C. S. Page, Jayde M. Bertoch, Fadi Alsarhan, Jaehee Kim, Amr A. Alazali, Ritwik R. Srinivas, Xiao Xie, Audrey G. Reeves, Kacper Skakuj, Theodore G. Coffey, Scott C. Virgil, Jordan Nafie, Dan He, Nam Dao, Amanda L. Gunawan, Rina Dukor, Andreas Stahl, F. Dean Toste, Christopher J. Chang

## Abstract

Life is predicated on chirality, a molecular asymmetry akin to the left and right versions of human hands. Here we show that privileged protein residues are predisposed for chiral regulation. We developed enantiomeric oxaziridine reagents that systematically identify pro-(*S*) and pro-(*R*) methionine oxidation sites across proteomes that can be erased by stereospecific methionine sulfoxide reductase enzymes A and B, respectively. These probes reveal that chiral regulation of methionine oxidation-reduction processes can allosterically regulate protein function, as shown in cell and murine models of oxidative stress where selective (*R*)-methionine sulfoxide formation on M69 of biphenyl hydrolase-like protein leads to hydrolase inhibition and amplification of proteome *N*-homocysteinylation modifications. This work introduces a platform for characterizing sites of asymmetric methionine oxidation and the functional consequences concomitant with an individual chiral single-atom modification.

## Introduction

Chirality, or the asymmetric orientation of atoms in space, is an intrinsic property of life^1,2^. Nature has evolved preferential handedness as a progenitor for many fundamental aspects of biology. Indeed, the diverse stereochemical ornamentation of natural products and the enantiopurity present in biomacromolecules such as left-handed amino acids or right-handed nucleic acids provide molecular surfaces for selective recognition and regulation events (Fig. 1a)^3,4^. Within proteins, this homochiral structuring is further stratified through the introduction of post-translational modifications (PTMs). In this context, methionine (Met), one of only two sulfur-containing amino acids, is highly susceptible to post-translational modification. Upon reaction with monooxygenases or reactive oxygen species (ROS), Met converts into methionine sulfoxide (MetO)^5^. The oxidation of the thioether moiety introduces a chiral center at sulfur that partitions into two distinct diastereomers: (*S*)*-* and (*R*)*-*MetO^6–8^. These (*S*) and (*R*) single-atom oxygen marks on proteins can be removed through stereospecific reduction of the sulfoxide using methionine sulfoxide reductases (Msr) A and B, respectively^9,10^ (Fig. 1a). Whereas there is direct evidence for the stereospecific reduction of MetO epimers on proteins, the extent of stereospecific oxidation of Met on proteins has not yet been fully realized^11–13^ (Fig. 1a). Critically, Met-based redox events are associated with negative consequences such as the development and progression of neurodegenerative^14^ and cardiovascular diseases^15^, but also positive regulation of life span^16^, protein function^17–19^, and signal transduction^20^.

**Fig 1:**
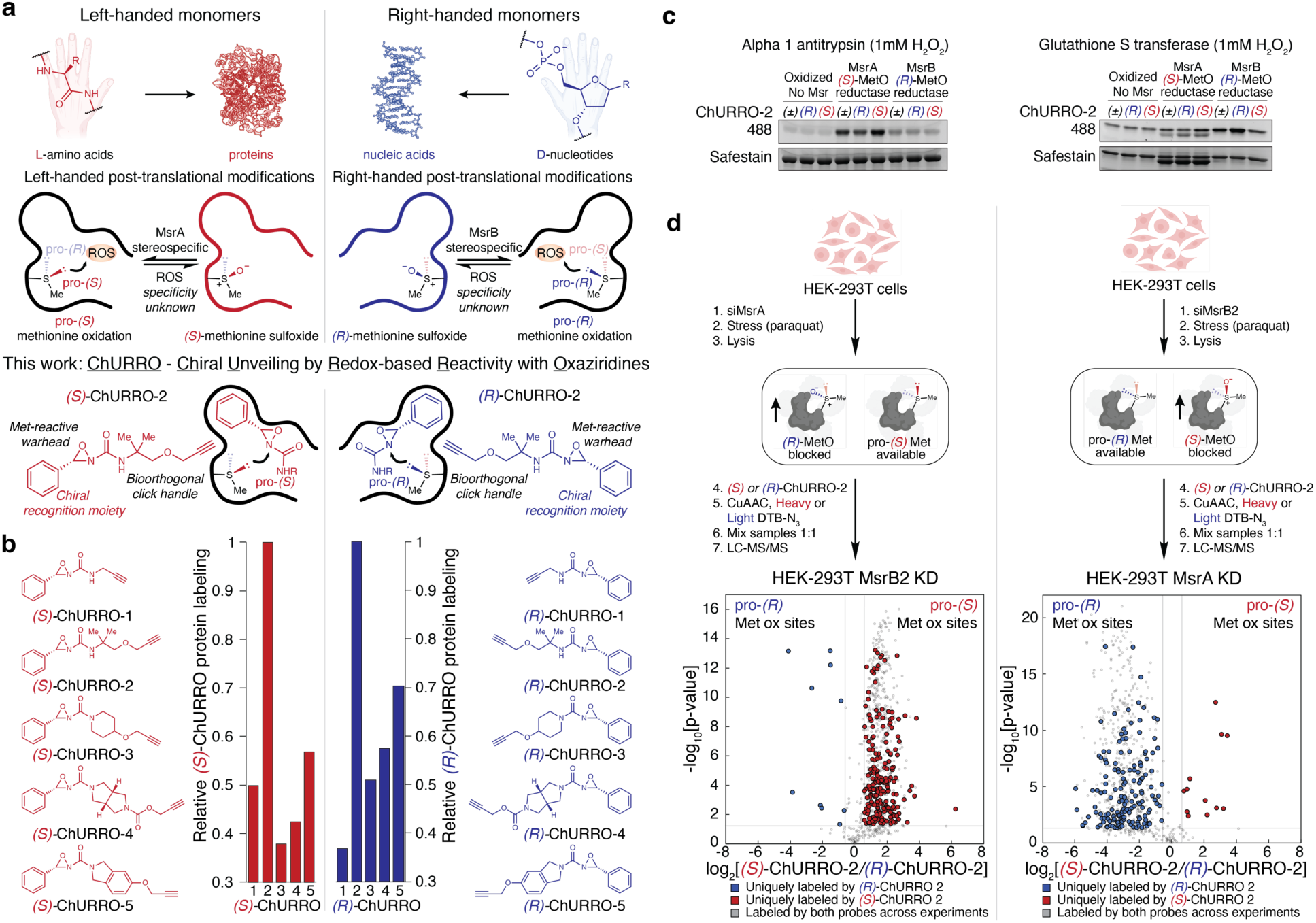
ChURRO for detection of prochiral methionine oxidation sites. **a**, (Top) Homochirality paradigm present within the biomacromolecules of life. (Middle) Chiral post-translational modification of Met and native Met redox regulation. (Bottom) ChURRO strategy for the global detection of prochiral Met oxidation sites. **b**, Development of structurally diverse ChURRO library identifies robust ChURRO probes with high methionine labeling in HEK-293T lysates. **c**, Msr rescue of oxidized proteins and labeling by ChURRO-2 using purified MsrA and MsrB enzymes and target substrates *in vitro*. **d**, ChURRO-ABPP strategy to validate detection of prochiral Met oxidation sites. HEK-293T cells were incubated with siRNA (50 nM) to knock-down endogenous MsrB2 or MsrA followed by addition of paraquat (100 µM) to bias (*R*)- or (*S*)-MetO formation on proteins. Following lysis, lysates were treated with equimolar (200 µM) doses of (*S*)- or (*R*)-ChURRO-2 and labeled with isotopically-encoded desthiobiotin tags through Cu-catalyzed click chemistry. Proteomes were then combined, enriched using streptavidin, tryptically digested, and analyzed by LC-MS/MS. Volcano plots of ChURRO-ABPP identifying prochiral Met oxidation sites in oxidatively-stressed HEK-293T lysates lacking either (*R*)- or (*S*)-MetO rescue mechanisms were then generated (n = 12 technical MS replicates).

Systematically identifying chiral single-atom modification sites on proteins has remained an elusive endeavor, with most examples performed on a protein-by-protein basis using purified Msr enzymes^19,21,22^. With growing evidence of stereospecific MetO redox regulation impacting physiology^18,19,23^, along with incongruencies in the subcellular distribution of both MsrA and MsrB enzymes to fully resolve racemic MetO epimer formation in eukaryotes^24^, we sought to decipher the extent of stereospecific Met oxidation at the proteome level. Inspired by work by Cravatt that identified enantioenriched druggable hotspots in proteomes^25–27^, along with work by our laboratories and others on synthetic methods for selective protein Met bioconjugation^28–31^, we developed an enantiomeric oxaziridine probe platform to systematically identify and interrogate protein chiral microenvironments in which diastereoselective Met oxidation may occur. This method, termed <u>ChURRO</u> (<u>Ch</u>iral <u>U</u>nveiling by <u>R</u>edox-based <u>R</u>eactivity with <u>O</u>xaziridines) (Fig. 1a) provides a chemical framework for the deconvolution of prochiral Met oxidation sites in purified proteins, cell lysates, and tissues, allowing for the rapid identification of putative regulatory Met sites that are privileged for stereospecific post-translational modifications.

### Development of ChURRO for identifying prochiral methionine oxidation sites in proteins and proteomes

Drawing inspiration from reports of enhanced diastereoselectivity for oxidation of sterically constrained thioethers as well as certain Met oxidation sites on proteins with ROS^32,33^, we reasoned that the extent of diastereoselectivity in Met oxidation could be predicated by the protein microenvironment in which a given Met residue resides. The asymmetric oxidation of Met using ROS would then operate by accessibility of either the pro-(*S*) or pro-(*R*) lone pair on the sulfur atom. We hypothesized that a pair of enantiomeric oxaziridine probes could discriminate between asymmetric Met environments by reacting to form sulfilimine adducts as isoelectronic proxies for MetO, while allowing for downstream functionalization and enrichment of ligated proteins^34^. In this design, the chiral *C*–phenyl moiety of a ChURRO probe serves as a hydrophobic directing group for the reactive warhead to access putative prochiral Met oxidation sites by differential protein binding of the enantiomers (Fig. 1a). To meet this goal, we devised a library of ten chemically-diverse (*S*)- and (*R*)-ChURRO probes comprised of alkyl, cyclic, and heterocyclic oxaziridines, which were isolated at preparative scale with >99% ee through chiral SFC separation of the racemic material, and their absolute configuration was determined through experimental and computational vibrational circular dichroism measurements (Supplementary Fig. 1a-e and Supplementary Fig. 2a-e).

We first sought to apply ChURRO in model proteins and proteomes. Screening our ChURRO library in proteome labeling assays revealed that ChURRO-2 excelled in protein labeling compared to other ChURRO reagents and retained high stability to epimerization (Fig. 1b, Supplementary Fig. 3a,b). Further *in vitro* and *in silico* diastereoselectivity studies of (*S*)- and (*R*)-ChURRO-2 suggest differential reactivity of the sulfur lone pairs due to increased steric constraints in the transition state, indicating that these reagents may be used to probe Met oxidation sites in which stereoselectivity is controlled by the protein environment (Supplementary Fig. 3d-f). We further tested this hypothesis on a modest set of proteins with Met oxidation sites whose selective reduction by Msr enzymes was previously reported^35,36^. In-gel fluorescence assays were conducted by oxidizing these proteins then incubating them with either MsrA or MsrB enzyme revealing that the cognate ChURRO-2 enantiomer can correctly ligate a corresponding (*S*) or (*R*)-MetO site (Fig 1c). Indeed, further LC tandem MS (LC-MS/MS) measurements corroborate these findings by detecting sulfilimine mass adducts on the exact (*S*)- or (*R*)-MetO site when treated with the respective ChURRO enantiomer (Supplementary Fig. 3g).

Next, we envisioned validating if ChURRO could detect pro-(*S*) or pro-(*R*) Met oxidation sites across proteomes by performing quantitative proteomic studies in a model of (*S*)- or (*R*)-Met oxidation in HEK-293T cells. We generated these mirrored models through genetic knock-down of the stereospecific reductases present in mitochondria, MsrA or MsrB2, followed by oxidative stress through mitochondrial uncoupling to exacerbate (*S*)-or (*R*)-MetO formation, respectively (Fig. 1d). We observed stereochemically-biased protein labeling when a prochiral Met oxidation site is blocked, establishing the ability of the ChURRO platform to systematically detect prochiral Met oxidation sites in proteomes using the respective enantioprobe in cell lysates (Fig. 1d). Live-cell ChURRO labeling was examined but found to be incompatible due to cytotoxicity and membrane permeability constraints (Supplementary Fig. 4a-c). Armed with this information, we performed quantitative proteomic experiments in native HEK-293T cell lysates to identify 175 redox-sensitive methionines comprised of 133 pro-(*S*) and 42 pro-(*R*) Met oxidation sites that were highly enriched (>1.5-fold) (Supplementary Fig. 4d). Critically, no prochiral oxidation site displayed cross-reactivity with the opposite ChURRO enantiomer and nearly 50% of all prochiral Met oxidation sites found reside on undruggable proteins according to the ChEMBL database^37^, underscoring the selectivity and therapeutic potential of these sites (Supplementary Fig. 4e,f). Together, these data establish the ability of the ChURRO platform to reveal and discriminate prochiral methionine oxidation sites at the protein and proteome-wide level.

### ChURRO profiling in the mitochondria reveals structural and functional characteristics of prochiral methionine oxidation sites

After establishing that ChURRO can discriminate prochiral Met oxidation sites in a high-throughput manner, we applied this method to profile stereospecific Met oxidation in isolated mitochondria derived from liver tissue. The liver is an essential organ for maintaining metabolic homeostasis and is highly enriched in mitochondria, a subcellular hotspot for redox-mediated processes. We reasoned that this model would reveal a potential wealth of functional prochiral Met oxidation sites and that organelle-specific enrichment would both minimize background from other cellular compartments and increase signal by providing a proteome stimulated by the redox environment present in the mitochondria. A combination of sucrose and Percoll gradients afforded enriched mitochondria via density ultracentrifugation from wild-type mouse livers with enhanced purity compared to traditional sucrose gradient- or immunoprecipitation-based techniques (Fig. 2a, Supplementary Fig. 5a). The enrichment of mitochondria was confirmed by immunoblot analyses of validated subcellular protein markers as well as shotgun proteomics to determine the subcellular localization of the protein content (Fig. 2b, Supplementary Fig. 5b). With these purified mitochondria in hand, we performed a rapid assessment of putative pro-(*S*) and pro-(*R*) Met oxidation sites by protein gel assays which displayed sufficient resolution to distinguish prochiral sites (Fig. 2c). Encouraged by these qualitative results, we then employed quantitative proteomics to identify prochiral Met oxidation sites in the mitochondria by equimolar treatment of both ChURRO-2 enantiomers to detect pro-(*S*) and pro-(*R*) Met oxidation sites and identified 138 redox-sensitive Met oxidation sites, of which 60 were highly enriched (>1.5-fold) with 39 pro-(*S*) and 21 pro-(*R*) sites described (Fig. 2d,e). We also identified hyper-reactive Met sites in these mitochondria by treating with low and high doses of racemic ChURRO-2 probe and determined that the previously identified prochiral Met oxidation sites were not simply hyper-reactive Met sites (Supplementary Fig. 5c,d), showcasing a stereochemical distinction in asymmetric Met oxidation that supersedes hyper-reactivity.

**Fig 2:**
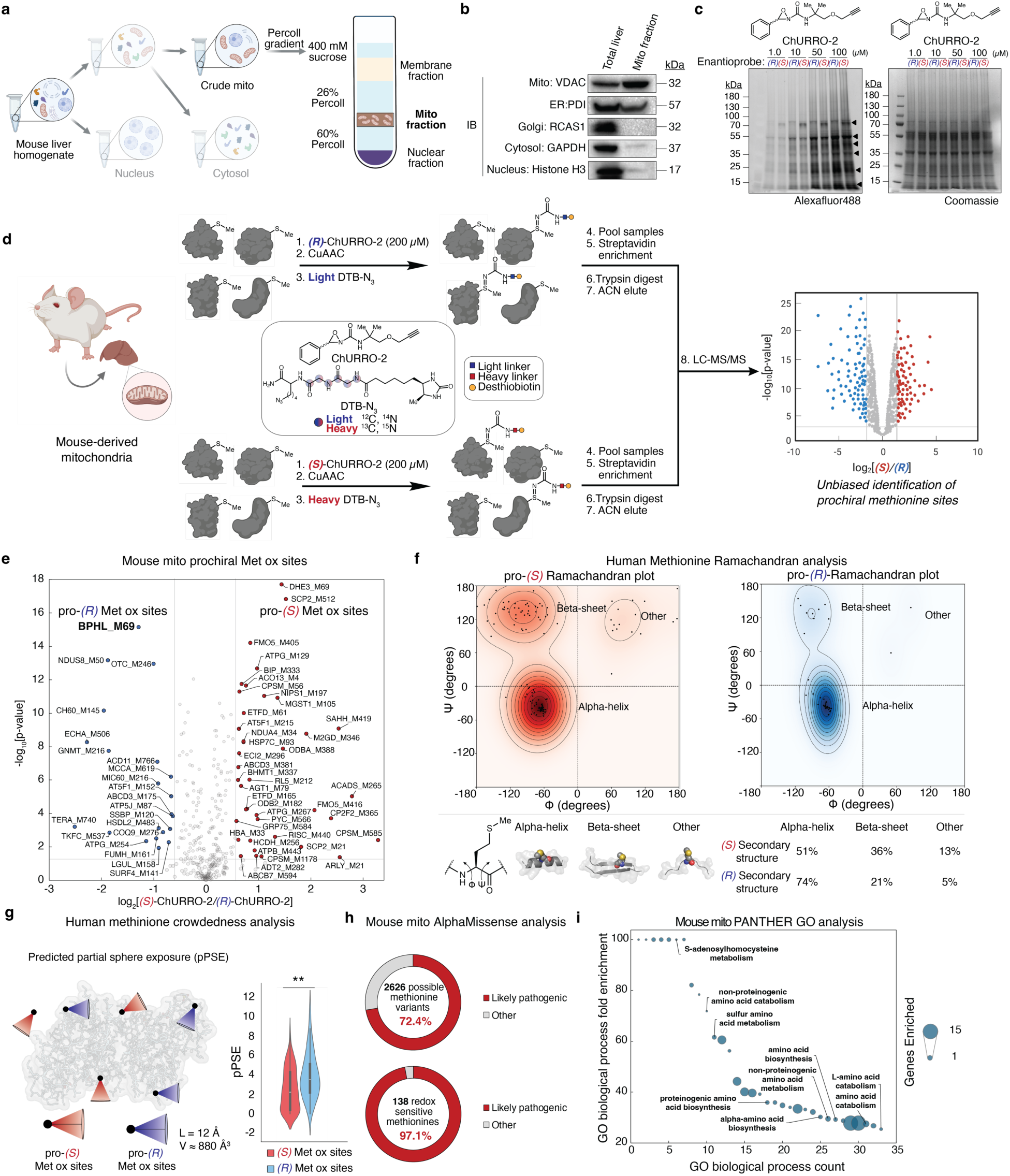
Chemoproteomic detection and stratification of prochiral methionine oxidation sites in cellular proteomes and the mitochondria. **a**, Density centrifugation and Percoll gradient scheme for isolation of highly pure mitochondria from mouse liver tissue. **b**, Immunoblot analysis of mitochondrial purity using various subcellular protein markers. **c**, (*R*)- and (*S*)-ChURRO-2 prochiral Met profile in purified mitochondria with increasing doses of reagent visualized by in-gel fluorescence. **d**, ChURRO proteomic strategy to identify prochiral Met oxidation sites. Mitochondria were incubated with equimolar doses of (*R*)- or (*S*)-ChURRO-2 and labeled using isotopically-encoded desthiobiotin tags through Cu-mediated click chemistry. Proteomes were then combined, enriched using streptavidin, tryptically digested, and analyzed by LC-MS/MS. **e**, Volcano plot analysis of statistically significant and highly enriched pro-(*R*) and pro-(*S*) Met oxidation sites in the mitochondria (n = 12 technical MS replicates). **f**, Ramachandran plots of enriched (*S*)- and (*R*)-Met oxidation sites determined from ChURRO-2 proteomic experiments in HEK-293T lysates. Phi and psi angles of identified prochiral Met oxidation sites were extracted from the AlphaFold database to visualize and quantify local environment. **g**, Met crowdedness of pro-(*S*)- and pro-(*R*)-Met oxidation sites determined from ChURRO-2 proteomic experiments in HEK-293T lysates. Crowdedness was determined by counting the number of residues within a projected cone using part-sphere exposure (pPSE) analysis for each individual Met site. **h**, AlphaMissense prediction of 2626 missense variants of identified mitochondrial redox sensitive Met oxidation sites. **i**, Gene ontology analysis of mitochondrial redox sensitive Met oxidation sites.

With this landscape of newly revealed prochiral Met oxidation sites across both human and mouse proteomes, we next sought to identify key biochemical parameters that promote asymmetric Met oxidation and assess their impact on protein function. In this context, a protein’s fold is described by its amino acid torsion angles, phi and psi, which provide a quantitative stereochemical assessment of local fold. Using the AlphaFold database^38^, we extracted the phi and psi angles of each uniquely identified pro-(*S*) and pro-(*R*) Met oxidation site to generate (*S*)- and (*R*)-Ramachandran plots (Fig. 2f). We then determined the distribution of prochiral Met sites in the phi and psi regions for common secondary protein structures. Interestingly, the pro-(*R*) sites display strong right-handed alpha helical character, while pro-(*S*) sites display a preference for beta sheet folds, which stratify these sites from average Met residues which display more equal helical and beta sheet distributions^39^. We further calculated the steric environment of all identified Met sites with ChURRO probe pair using a predicted partial sphere exposure (pPSE) analysis, where a greater pPSE value denotes more steric encumbrance (Fig. 2g). This analysis revealed that the ChURRO-2 probe can discriminate between protein steric environments, with pro-(*S*) sites residing in more encumbered environments relative to pro-(*R*). Taken together, these results suggest that local secondary structure and steric environment provide the scaffolding for asymmetric Met oxidation on proteins.

The list of prochiral Met oxidation sites identified from our mitochondrial dataset were validated against previously annotated mitochondrial proteins^40^, enabling a more selective proteome analysis to stratify and prioritize Met redox sites of putative function for further study. Predicted mutation pathogenicity scoring through AlphaMissense^41^ analysis of the 138 redox-sensitive Met oxidation sites revealed that 72.4% of all possible 2626 missense mutations are predicted to be pathogenic, and that 97.1% of the 138 sites have at least one predicted pathogenic mutation, highlighting the biological significance of these privileged residues in mitochondria (Fig. 2h). Along these lines, gene ontology (GO) analysis revealed a strong enrichment in proteins involved with amino acid biosynthesis, particularly proteinogenic and non-proteinogenic sulfur amino acids, pointing towards potential regulatory roles in amino acid metabolism for certain prochiral Met oxidation sites (Fig. 2i).

### Human BPHL contains a regulatory allosteric prochiral methionine oxidation site at M69

To showcase the consequences of asymmetric Met oxidation on protein function, we chose to characterize a highly enriched and significant pro-(*R*) M69 site present on mouse biphenyl hydrolase-like protein (BPHL) as a representative example of prochiral Met regulation. BPHL is a mitochondrially-localized serine hydrolase that is most commonly associated with prodrug activation but is also suggested to play a significant role in homocysteine thiolactone (HCTL) detoxification^42,43^. HCTL levels reciprocally correlate with those of homocysteine (HC), a non-proteinogenic amino acid involved in cysteine and Met biosynthesis, where HC is converted to HCTL through an error-editing mechanism of methionyl–tRNA synthetase^44,45^ (Fig. 3a). Aberrant overproduction of HCTL from HC introduces a highly electrophilic species that reacts with protein-bound amines via *N*-homocysteinylation. *N*-homocysteinylation adducts are known to induce protein damage, aggregation, immunogenicity, and cellular toxicity^46^. Analysis of available crystallographic data for human BPHL revealed that the identified M69 site lies in an allosteric hydrophobic pocket proximal to the catalytic triad^47^ (Fig. 3b), and a multiple-sequence alignment revealed sequence conservation of this Met site in humans, mice, and many orders of animal phyla, suggesting this site may have an evolved functional role (Supplementary Fig. 6a).

**Fig 3:**
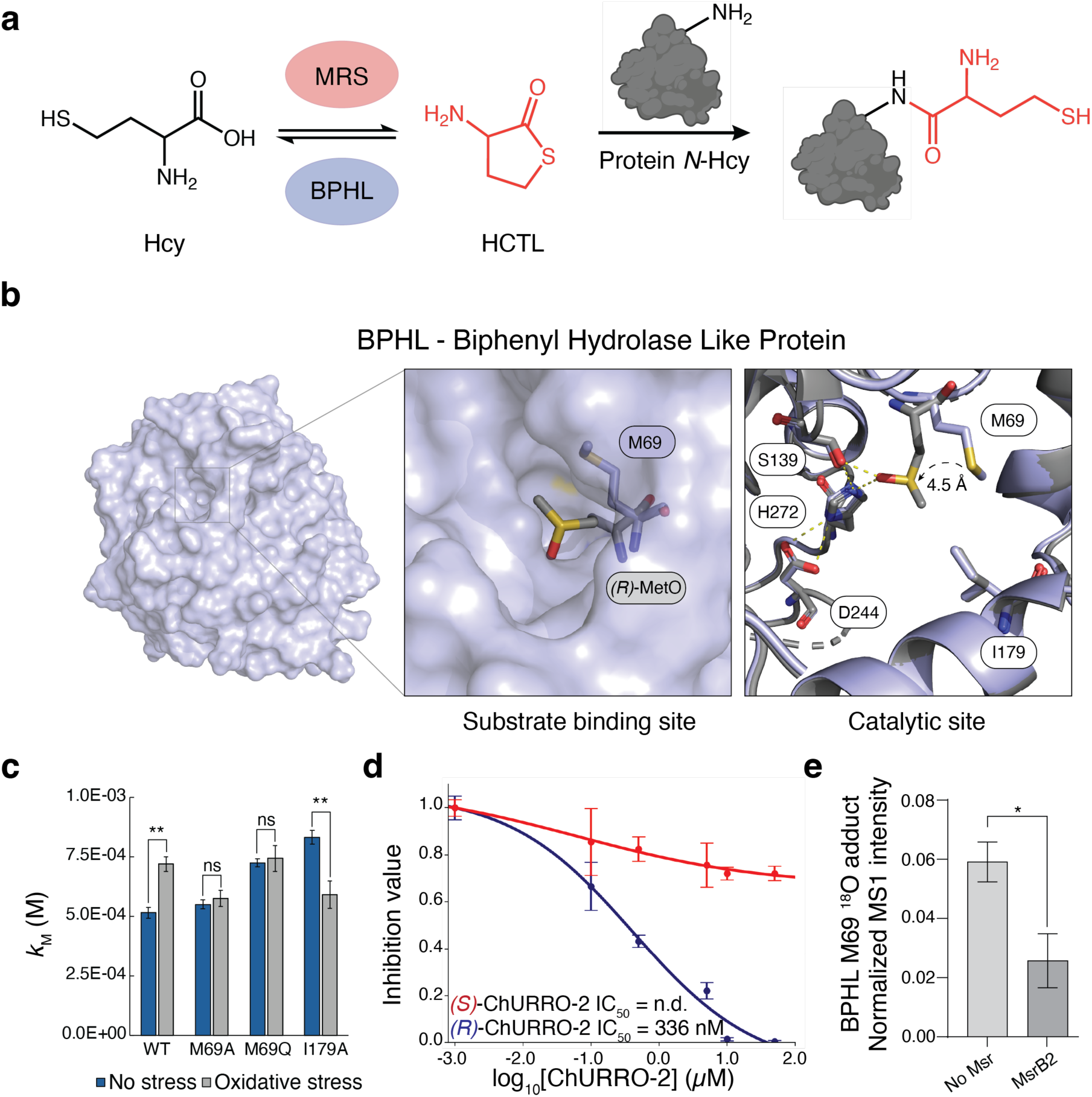
Human BPHL contains an allosteric regulatory M69 site. **a**, Homocysteine thiolactone (HCTL) regulation by BPHL and methionyl–tRNA synthetase (MRS) and subsequent protein *N*-homocysteinylation with HCTL. **b**, RoseTTAFold All-Atom structure of human BPHL with modeled (*R*)-MetO at the 69 site. **c**, *k*_M_ of BPHL variants under conditions of no stress or oxidative stress (n = 3 technical replicates, error bars represent standard deviation). **d**, Dose-response curve of WT BPHL treated with (*S*)- and (*R*)-ChURRO-2 with calculated IC_50_ values (n = 3 technical replicates, error bars represent standard deviation). **e**, LC-MS/MS quantification of ^18^O-H_2_O_2_ adducts at BPHL M69 with MsrB2 treatment (n = 3 technical MS replicates, error bars represent standard deviation).

To assess the impact of pro-(*R*) M69 oxidation on BPHL, we modeled an (*R*)-MetO at the 69 site using Baker’s RoseTTA-Fold All-Atom platform^48^ (Fig. 3b). These results indicate that pro-(*R*) M69 oxidation induces sequestration of the sulfoxide zwitterion by hydrogen bonding interactions with the catalytic serine leading to obstruction of the substrate binding site. To further understand the functional consequences of M69 oxidation, we generated a panel of BPHL variants by installing a small hydrophobic moiety (M69A), a MetO bioisostere (M69Q), and a second sphere hole to increase the binding pocket size (I179A) (Supplementary Fig. 6b). Michaelis-Menten kinetic studies on these purified protein variants using the native substrate HCTL reveal a conserved decrease in *k*_cat_ upon oxidation irrespective of M69 alluding to high redox-sensitivity of the enzyme (Supplementary Fig. 6c-f). However, the measured binding affinity *k*_M_ of HCTL is significantly altered in an M69-dependent manner that increases upon oxidation for the wild-type enzyme but is rescued by either M69A or I179A mutation. Furthermore, the bioisostere M69Q displays a similar *k*_M_ to that of oxidatively stressed wild-type enzyme, suggesting an inhibitory pathway through substrate occlusion (Fig. 3c). Together, these data support a mixed inhibition mechanism where at one axis is driven by M69 oxidation-based obstruction of HCTL binding (Supplementary Fig. 6g,h). We further probed the role of stereospecific M69 oxidation on protein inhibition by in-gel fluorescence, shotgun proteomics, and dose-response studies on purified wild-type BPHL showcasing that (*R*)-ChURRO-2 ligation or (*R*)-MetO formation using asymmetric oxygen-atom transfer reagents preferentially inhibit the protein (Fig. 3d, Supplementary Fig. 6i-k). These rigorous stereochemical studies are punctuated by directly detecting formation of (*R*)-MetO at the 69 position by measuring MetO adducts with isotopically-labeled H_2_O_2_, which controlled for non-specific oxidation events and were significantly attenuated following MsrB2 treatment (Fig. 3e).

### Protein *N*-homocysteinylation is regulated by a BPHL/MsrB2 axis in cell and tissue models of oxidative stress

Following *in vitro* characterization of BPHL’s prochiral regulatory M69 site on purified protein models, we explored how this stereospecific single-atom modification impacts downstream protein *N*-homocysteinylation at the proteome level. To this end, we devised a cellular *N*-homocysteinylation assay that utilizes mitochondrial uncoupling reagents for *in situ* ROS generation, followed by dosing of HCTL to examine the extent of protein *N*-homocysteinylation by leveraging a chemoselective native chemical ligation strategy reported by Wang^49^ (Fig. 4a). Screening a panel of hepatic and renal cell lines, we found that HepG2 and HEK-293T cells showed significantly high relative expression levels of BPHL compared to other models (Supplementary Fig. 7a). We chose HEK-293T cells for *in cellulo* characterization of protein *N*-homocysteinylation levels owing to their ease of transfection for subsequent genetic perturbation studies. Expression of an epitope-tagged BPHL confirmed selective ligation of (*R*)-ChURRO-2 in these cells by immunoprecipitation studies (Supplementary Fig. 7b). To further validate (*R*)-MetO formation and regulation, we measured MsrA or MsrB2 engagement of BPHL *in cellulo*, revealing a significant increase of MsrB2 co-precipitating with BPHL under conditions of oxidative stress in an M69-dependent manner, suggesting that MsrB2 is an eraser of BPHL (*R*)-Met69O (Fig. 4b, Supplementary Fig. 7c). We next measured the cellular consequences of BPHL (*R*)-Met69O on proteome *N*-homocysteinylation levels by generating CRISPR knockouts of MsrA or MsrB2 in HEK-293T cells followed by treatment with paraquat to induce oxidative stress (Supplementary Fig. 7d). Global protein *N*-homocysteinylation levels were comparable for control and MsrA KO lines but displayed a significant 2-fold increase upon KO of MsrB2. Elevated proteome *N*-homocysteinylation levels can be rescued by over-expression of wild-type BPHL in native and MsrA KO genotypes but not in MsrB2 KO lines (Fig. 4c, Supplementary Fig. 7e). These cellular data support our *in vitro* studies suggesting BPHL activity is stereospecifically regulated by Met redox status, which in turn amplifies downstream protein *N*-homocysteinylation across proteomes.

**Fig 4:**
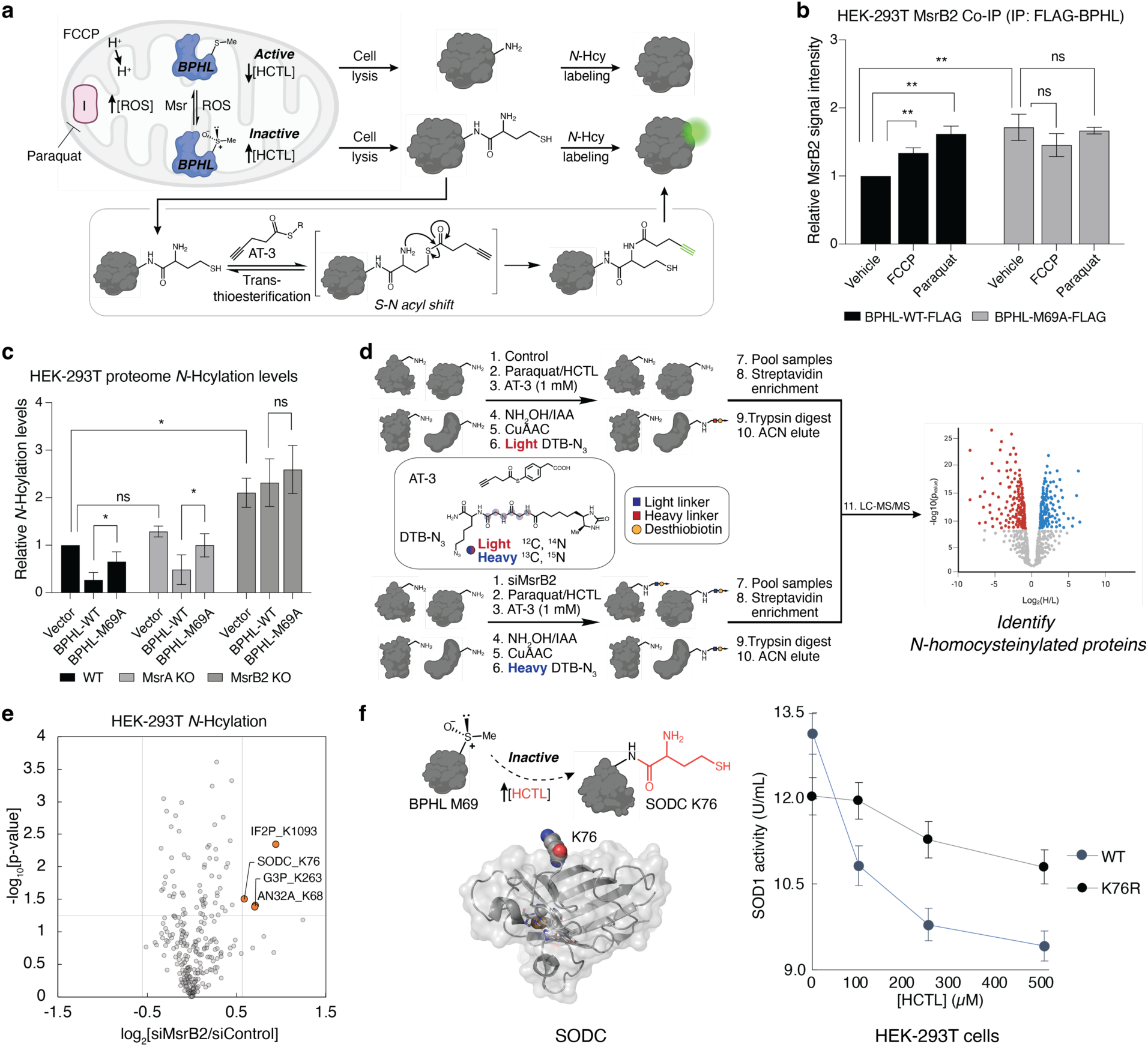
MsrB2 regulates BPHL through stereospecific methionine redox regulation to amplify cellular protein *N*-homocysteinylation. **a**, Mitochondrial uncoupler assay to assess protein *N*-homocysteinylation status as a function of BPHL Met redox status using thioester probe AT-3 to label sites of protein *N*-homocysteinylation. **b**, BPHL and Msr co-immunoprecipitation analysis following mitochondrial uncoupling in HEK-293T cells (n = 3 cell technical replicates, error bars represent standard deviation). **c**, Relative protein *N*-homocysteinylation levels following loss of mitochondrial (*S*)- or (*R*)-MetO rescue mechanisms (n = 3 cell technical replicates, error bars represent standard deviation). **d**, Chemoproteomic scheme for assessing global *N*-homocysteinylation changes in MsrB2 deficient cell models. HEK-293T cells were incubated with siRNA (50 nM) to knock-down endogenous MsrB2 addition of paraquat (100 μM) and HCTL (500 μM). Following lysis, lysates were treated with equimolar (1 mM) doses of AT-3 and labeled with isotopically-encoded desthiobiotin tags through Cu-catalyzed click chemistry. Proteomes were then combined, enriched using streptavidin, tryptically digested, and analyzed by LC-MS/MS. **e**, Volcano plot of hyper *N*-homocysteinylated sites detected in oxidatively-stressed HEK-293T lysates containing or lacking (*R*)-MetO rescue mechanisms (n = 4 technical MS replicates). **f**, (Left) Alphafold2 model of SOD1 with K76 site shown. (Right) Cell activity assays of HEK-293T cells expressing wild-type or K76R SOD1 upon titration of HCTL (n = 3 cell technical replicates, error bars represent standard deviation).

The crosstalk between a redox-regulated, chiral single-atom modification and global proteome *N*-homocysteinylation prompted us to investigate its biochemical genesis in models of oxidative stress. In particular, hyperhomocysteinemia is a metabolic disorder characterized by elevated levels of sulfur-containing metabolites HC and HCTL, which has been linked to exacerbation of oxidative stress and progression of neurodegenerative, cardiac, and hepatic diseases^50–54^. Despite observed correlations between higher levels of protein *N*-homocysteinylation and disease, underlying biochemical mechanisms of how hyperhomocysteinemic pathologies arise remain insufficiently understood^55^. We therefore sought to identify and characterize specific protein targets of *N*-homocysteinylation in a cellular model of elevated (*R*)-MetO sites.

We elected for a quantitative chemoproteomic approach to identify protein targets of *N*-homocysteinylation by increasing (*R*)-MetO levels through knock-down of MsrB2, followed by mitochondrial uncoupling (Fig. 4d, Supplementary Fig. 7f). We identified a set of highly enriched (>1.5-fold) *N*-homocysteinylated sites. Included is a K76 site on the enzyme SOD1, a crucial ROS maintenance enzyme that catalyzes disproportionation of superoxide to hydrogen peroxide and oxygen. Since SOD1 activity is highly regulated through a palette of PTMs^56,57^, we measured the impact of K76 *N*-homocysteinylation on SOD1 activity in HEK-293T cells upon titration of HCTL. These assays reveal that SOD1 activity is indeed attenuated at higher concentrations of HCTL which may be rescued by mutation to K76R, suggesting that K76 *N*-homocysteinylation is a negative regulator of SOD1 activity (Fig. 4f). Taken together, these data support a model in which a stereospecific single-atom modification on M69 of BPHL, a biocatalyst for HCTL degradation, inactivates its regulatory function leading to higher levels of HCTL and amplification of downstream protein *N*-homocysteinylation across the proteome.

## Discussion

Reversible, regulated oxidation-reduction cycles on protein Met residues give rise to chiral single-atom modifications across the proteome. The stereochemistry that engenders these events has been partially elucidated through characterization of MetO reduction events catalyzed by stereospecific Msr enzymes. Here, we show an inherent stereospecificity in PTM oxidation events that occur at privileged prochiral Met oxidation sites driven by the structural asymmetry present in protein pockets. We challenge the dogma of non-specific MetO epimer formation in proteins through development of a biomimetic chiral oxidation reagent platform to detect these privileged protein sites. The ChURRO method enables systematic identification of prochiral Met oxidation sites on purified proteins and in whole proteomes. Using this approach, we systematically identified new prochiral Met oxidation sites in human whole-cell proteomes, as well as in mouse liver-derived mitochondria, a subcellular locus of redox homeostasis and regulatory processes. In-depth analysis of the local microenvironments of these prochiral Met oxidation sites revealed a bias for right-handed alpha helices around pro-(*R*) sites, and beta sheets for pro-(*S*) sites, which we posit favor asymmetric Met oxidation. Together, these data elucidate foundational local environment preferences as well as protein functions that stratify and distinguish prochiral Met oxidation sites from other Met residues within the proteome.

To establish the functional consequences for regulation of prochiral Met oxidation sites, we performed biochemical studies of a representative pro-(*R*) Met oxidation site, M69, on the mitochondrial serine hydrolase BPHL. We show that this prochiral redox site plays an important role in HC metabolism through the hydrolysis of the reactive intermediate HCTL. Through site-directed mutagenesis studies, we revealed that HCTLase activity is dependent upon M69 redox status and its oxidation to (*R*)-MetO can function as a negative allosteric regulator of protein activity, resulting in increased HCTL levels and protein *N*-homocysteinylation in cells. We also found that BPHL is a substrate for the (*R*)-MetO reductase MsrB2, but not the corresponding (*S*)-MetO reductase MsrA when subjected to elevated ROS levels, suggesting stereochemically privileged (*R*)-MetO formation and reduction on this protein. We further characterized substrates of protein *N*-homocysteinylation in cell models of elevated (*R*)-MetO, where loss of MsrB2 leads to K76 *N*-homocysteinylation and downregulation of SOD1 activity. Indeed, these findings support a newly identified PTM crosstalk pathway between Met oxidation and protein *N*-homocysteinylation, where chiral regulation of oxidation-reduction events at a specific M69 site on BPHL leads to formation of (*R*)-MetO and allosteric inhibition of BPHL activity, resulting in elevated HCTL levels and downstream protein *N*-homocysteinylation. This work presages that chiral protein Met modifications can serve as privileged allosteric signaling moieties that reveal new native physiology and disease vulnerabilities at the single-atom level for potential detection and intervention.

## Methods

### Stereospecifically reduced protein Gel-ABPP

Proteins (Alpha 1 antitrypsin: Sigma, Cat. 178251; Glutathione S transferase: abcam, Cat. Ab89494) were diluted to 2 µM in PBS. To 50 µL of protein was added 1 µL of a stock of either (*R*)- or (*S*)-ChURRO-2 in DMSO for a final concentration of 0, 0.2, 1.0, 2.0, 4.0, 10, or 20 µM. Samples were allowed to incubate at 23 °C for 1 hour. Excess oxaziridine was then quenched by addition of 1 µL *N*-acetyl methionine (100 µM) and incubation at 23 °C for 30 minutes. 1 µL of Alexafluor488-N_3_ (1.2 equiv.) was added to each sample with 3 µL TBTA (stock in 1:4 DMSO:*^t^*BuOH, 100 µM), 1 µL TCEP (1 mM), and 1 µL CuSO_4_ (1 mM). Proteins were unfolded by addition of 5 µL 1.2% SDS in PBS (0.1% w/v) and CuAAC was allowed to proceed at 23 °C for 1 hour with rocking. Then, 20 µL of 4X Laemmli’s buffer (Bio-Rad Laboratories, Inc.; 1610747) + 10% BME was added, and samples were boiled at 95 °C for 8 minutes. 30 µL of sample was loaded per well and separated on precast 4–20% Novex Tris-Gly SDS-PAGE gels (Invitrogen) run at 160 V for 70 minutes. The gel was fixed in MQ water containing 10% acetic acid, 30% EtOH overnight then visualized using ChemiDoc MP (Bio-Rad) for measuring in-gel fluorescence (ex = 499 nm, em = 520 nm). The gel was rehydrated in MQ water for 1 hour then total protein level was assessed by Pierce Silver Staining (Thermo Fisher Scientific; 24612) according to the manufacturer’s protocol or Safestain incubation (Invitrogen; LC6065) and visualized under brightfield.

### Msr rescue of oxidized protein Gel-ABPP

Proteins (Alpha 1 antitrypsin: Sigma, Cat. 178251; Glutathione S transferase: abcam, Cat. Ab89494) were diluted to 2 µM in PBS. To 900 µL of protein was added 9 µL of a stock of H_2_O_2_ for the indicated final concentration. Samples were allowed to incubate at 37 °C for 1 hour. Then, to 900 µL of protein was added 9 µL of a stock of DTT for a final concentration of 1.5 mM. This parent stock was then split into 3 x 300 µL stocks, one stock for DTT only treatment, one was treated with 3 µL of a stock of human MsrA for a final concentration of 2 µM, and the final was treated with 3 µL of a stock *E. coli* MsrB for a final concentration of 4 µM. Samples were allowed to incubate at 37 °C for 1 hour. Samples were then desalted using Zeba™ Spin Desalting Columns (Thermo Fisher Scientific; 89882) according to manufacturer’s protocol. Following desalting, samples were split into 50 µL aliquots in PCR tubes then treated with 1 µL of a stock of either (±)-, (*R*)-, or (*S*)-ChURRO-2 in DMSO for a final concentration of 2 µM. Samples were allowed to incubate at 23 °C for 1 hour. Excess oxaziridine was then quenched by addition of 1 µL *N*-acetyl methionine (100 µM) and incubation at 23 °C for 30 minutes. 1 µL of Alexafluor488-N_3_ (1.2 equiv.) was added to each sample with 3 µL TBTA (stock in 1:4 DMSO:*^t^*BuOH, 100 µM), 1 µL TCEP (1 mM), and 1 µL CuSO_4_ (1 mM). Proteins were unfolded by addition of 5 µL 1.2% SDS in PBS (0.1% w/v) and CuAAC was allowed to proceed at 23 °C for 1 hour with rocking. Then, 20 µL of 4X Laemmli’s buffer (Bio-Rad Laboratories, Inc.; 1610747) + 10% BME was added, and samples were boiled at 95 °C for 8 minutes. 30 µL of sample was loaded per well and separated on precast 4–20% Novex Tris-Gly SDS-PAGE gels (Invitrogen) run at 160 V for 70 minutes. The gel was fixed in MQ water containing 10% acetic acid, 30% EtOH overnight then visualized using ChemiDoc MP (Bio-Rad) for measuring in-gel fluorescence (ex = 499 nm, em = 520 nm). The gel was rehydrated in MQ water for 1 hour then total protein level was assessed by Pierce Silver Staining (Thermo Fisher Scientific; 24612) according to the manufacturer’s protocol or Safestain incubation (Invitrogen; LC6065) and visualized under brightfield.

### hBPHL Gel-ABPP

hBPHL was diluted to 2 µM in PBS. To 50 µL of protein was added 1 µL of a stock of either (*R*)- or (*S*)-ChURRO-2 in DMSO for a final concentration of 0, 0.1, 0.5, 1.0, 2.5, 5.0, or 10 µM. Samples were allowed to incubate at 23 °C for 1 hour. Excess oxaziridine was then quenched by addition of 1 µL *N*-acetyl methionine (100 µM) and incubation at 23 °C for 30 minutes. 1 µL of Alexafluor488-N_3_ (1.2 equiv.) was added to each sample with 3 µL TBTA (stock in 1:4 DMSO:*^t^*BuOH, 100 µM), 1 µL TCEP (1 mM), and 1 µL CuSO_4_ (1 mM). Proteins were unfolded by addition of 5 µL 1.2% SDS in PBS (0.1% w/v) and CuAAC was allowed to proceed at 23 °C for 1 hour with rocking. Then, 20 µL of 4X Laemmli’s buffer (Bio-Rad; 1610747) + 10% BME was added, and samples were boiled at 95 °C for 8 minutes. 30 µL of sample was loaded per well and separated on precast 4–20% Novex Tris-Gly SDS-PAGE gels (Invitrogen) run at 160 V for 70 minutes. The gel was fixed in MQ water containing 10% acetic acid, 30% EtOH overnight then visualized using ChemiDoc MP (Bio-Rad) for measuring in-gel fluorescence (ex = 499 nm, em = 520 nm). The gel was rehydrated in MQ water for 1 hour then total protein level was assessed by coomassie brilliant blue staining and visualized under brightfield.

### Mitochondrial proteome Gel-ABPP

Mitochondrial lysate was diluted to 2.0 mg/mL in PBS. To 100 µL of protein was added 1 µL of a stock of either (*R*)- or (*S*)-ChURRO-2 in DMSO for a final concentration of 0, 1.0, 10, 50, or 100 µM. Samples were allowed to incubate at 23 °C for 1 hour. Excess oxaziridine was then quenched by addition of 1 µL *N*-acetyl methionine (100 µM) and incubation at 23 °C for 30 minutes. 2 µL of Alexafluor488-N_3_ (1.2 equiv.) was added to each sample with 6 µL TBTA (stock in 1:4 DMSO:*^t^*BuOH, 100 µM), 2 µL TCEP (1 mM), and 2 µL CuSO_4_ (1 mM). Proteins were unfolded by addition of 10 µL 1.2% SDS in PBS (0.1% w/v) and CuAAC was allowed to proceed at 23 °C for 1 hour with rocking. Then, 40 µL of 4X Laemmli’s buffer (Bio-Rad; 1610747) + 10% BME was added, and samples were boiled at 95 °C for 8 minutes. 30 µL of sample was loaded per well and separated on precast 4–20% Novex Tris-Gly SDS-PAGE gels (Invitrogen) run at 160 V for 70 minutes. The gel was fixed in MQ water containing 10% acetic acid, 30% EtOH overnight then visualized using ChemiDoc MP (Bio-Rad) for measuring in-gel fluorescence (ex = 499 nm, em = 520 nm). The gel was rehydrated in MQ water for 1 hour then total protein level was assessed by coomassie brilliant blue staining and visualized under brightfield.

### Michaelis-Menten activity assays of BPHL variants

All additions were performed using multi-channel pipettes. For proteins treated with oxidative stress, a stock of BPHL variant was diluted to 100 nM in PBS and treated with H_2_O_2_ (500 µM) at 37 °C for 30 mins. BPHL variants under no stress were treated with DMSO. To all samples, *N*-acetyl methionine (1.0 mM) was added and allowed to quench at 23 °C for 30 mins. 100 µL of a 100 nM protein solution containing BPHL was aliquoted into a black, clear flat-bottom 96-well plate in triplicate and allowed to equilibrate to 23 °C. Eight solutions containing 40, 20, 12, 8, 4, 2, 0.4, or 0 mM HCTL in PBS were freshly prepared and combined 1:1 with 4 mM DTNB in DMSO. 5 µL of the combined solutions containing varying concentrations of HCTL and equal concentrations of DTNB were administered to each well and absorbance at 412 nm was immediately monitored at 37 °C for 5 minutes using a plate reader (BioTek). Linear rates for the first 120 seconds were calculated to compute V_max_.

### SOD1 cell activity assays

SOD1 activity was assayed following transfection and HCTL treatment as above. Activity was determined using a lysate activity kit according to manufacturer’s protocols (Cayman; 706002).

### Dose response of WT BPHL against oxaziridines

All additions were performed using multi-channel pipettes. A stock of WT BPHL was diluted to 100 nM in PBS and aliquoted into a black, clear flat-bottom 96-well plate in triplicate and allowed to equilibrate to 23 °C. Wells were then treated by 1 µL addition of the appropriate concentration of (*R*)- or (*S*)-ChURRO-2, or (*R*)- or (*S*)-OT-Ox, ((*R*)-OT-Ox Sigma; 349003), ((*S*)-OT-Ox Sigma; 345350) while controls were treated with vehicle (DMSO). Oxidation was allowed to occur for 30 minutes at 23 °C, followed by quenching with 1 µL of *N*-acetyl methionine (1 mM) for 15 minutes at 23 °C. A solution containing both 40 mM HCTL and 4 mM DTNB was then prepared in the interim, and 5 µL was administered to each well. Absorbance at 412 nm was immediately monitored at 37 °C for 5 minutes using a plate reader (BioTek). Linear rates for the first 120 seconds were calculated to compute V_max_.

### General *N*-homocysteinylation assay

100 µL of desired proteome (4 mg/mL) was diluted to a concentration of 1 mg/mL using 8 M urea, 27 mM TCEP, pH 6.0 as the diluent for a final concentration of 6 M urea, 20 mM TCEP, pH 6.0 then incubated with 1 mM AT-3 thioester (*48*) for 30 minutes at 23 °C while rotating. Reaction was quenched by adding 20 µL NH_2_OH (50% v/v) and 50 µL iodoacetamide (400 mM) for 1 hour at 23 °C in the dark. Proteins were then precipitated by addition of 4 volumes of MeOH:CHCl_3_ (4:1.5), vortexed, and centrifuged at 12,000 xg for 15 minutes. The collected protein pellet was washed 1X with 500 µL chilled (-20 °C) MeOH via probe-tip sonication (5 s on, 20 s off, 20% A, 4-6 pulses) and resuspended in 200 µL 1.2% SDS/PBS via probe-tip sonication (5 s on, 20 s off, 20% A, 6-8 pulses). CuAAC was performed by addition of 4 µL Alexafluor488-N_3_ (1.2 equiv.), 12 µL TBTA (stock in 1:4 DMSO:*^t^*BuOH, 100 µM), 4 µL TCEP (1 mM), and 4 µL CuSO_4_ (1 mM). Then, 56 µL of 4X Laemmli’s buffer (Bio-Rad;1610747) + 10% BME was added and samples were boiled at 95 °C for 8 minutes.

20 µL of sample was loaded per well and separated on precast 4–20% Novex Tris-Gly SDS-PAGE gels (Invitrogen) run at 160 V for 70 minutes. The gel was fixed in MQ water containing 10% acetic acid, 30% EtOH overnight then visualized using ChemiDoc MP (Bio-Rad) for measuring in-gel fluorescence (ex = 499 nm, em = 520 nm). The gel was rehydrated in MQ water for 1 hour then total protein level was assessed by coomassie brilliant blue staining and visualized under brightfield. Band intensities were quantified via densitometry (ImageJ).

### rhBPHL cloning and M69 mutagenesis

WT Sequence (hBPHL, Uniprot ID: Q86WA6)

MVAVLGGRGVLRLRLLLSALKPGIHVPRAGPAAAFGTSVTSAKVAVNGVQLHYQQTGEGDHAVLLLPGMLGSGETDFGPQLKNLNKKLFTVVAWDPRGYGHSRPPDRDFPADFFERDAKDAVDLMKALKFKKVSLLGWSDGGITALIAAAKYPSYIHKMVIWGANAYVTDEDSMIYEGIRDVSKWSERTRKPLEALYGYDYFARTCEKWVDGIRQFKHLPDGNICRHLLPR VQCPALIVHGEKDPLVPRFHADFIHKHVKGSRLHLMPEGKHNLHLRFADEFNKLAEDFL Q

Cloned Sequence (rhBPHL)

MVAVLGGRGVLRLRLLLSALKPGIHVPRAGPAAAFGTSVTSAKVAVNGVQLHYQQTGEGDHAVLLLPG**M**LGSGETDFGPQLKNLNKKLFTVVAWDPRGYGHSRPPDRDFPADFFERDAKDAVDLMKALKFKKVSLLGWSDGGITALIAAAKYPSYIHKMVIWGANAYVTDEDSMIYEGIRDVSKWSERTRKPLEALYGYDYFARTCEKWVDGIRQFKHLPDGNICRHLLPRVQCPALIVHGEKDPLVPRFHADFIHKHVKGSRLHLMPEGKHNLHLRFADEFNKLAEDFLQ<u>HHHHHH</u>

<u>5’ – BPHL XbaI FOR– 3’</u>

CCCCTCTAGAATAATTTTGTTTAACTTTAAGAAGGAGATATACCATGGTAGCTGTGT TAGGCG

<u>5’ – BPHL XhoI REV– 3’</u>

GTGCTCGAGTTGTAAAAAGTCTTCGGCCAG

<u>5’ – BPHL M69A FOR– 3’</u>

GTATTACTGCTGCCCGGT**GC**GTTGGGTAGCGGTGAAAC

<u>5’ – BPHL M69A REV– 3’</u>

GTTTCACCGCTACCCAAC**GC**ACCGGGCAGCAGTAATAC

<u>5’ – BPHL M69L FOR– 3’</u>

GTATTACTGCTGCCCGGT**C**TGTTGGGTAGCGGTGAAAC

<u>5’ – BPHL M69L REV– 3’</u>

GTTTCACCGCTACCCAACA**G**ACCGGGCAGCAGTAATAC

<u>5’ – BPHL M69Q FOR– 3’</u>

GTATTACTGCTGCCCGGT**CA**GTTGGGTAGCGGTGAAAC

<u>5’ – BPHL M69Q REV– 3’</u>

GTTTCACCGCTACCCAAC**TG**ACCGGGCAGCAGTAATAC

<u>5’ – BPHL I179A FOR– 3’</u>

CAGCATGATCTACGAAGGG**GC**CCGCGATGTCAGCAAATG

<u>5’ – BPHL I179A REV– 3’</u>

CATTTGCTGACATCGCGG**GC**CCCTTCGTAGATCATGCTG

Molecular cloning was performed using a pET24a vector via traditional double-restriction digest using a codon-optimized synthetic gene-block of rhBPHL for *E. coli* expression (IDT) and overhang oligonucleotide primers (IDT). Insert PCR product was amplified and purified using QIAquick PCR purification (Qiagen). Both insert and vector were then double-restriction digested using XbaI and XhoI endonucleases (NEB). Plasmid was gel-purified using QIAquick gel purification (Qiagen) and insert was PCR purified. The digested insert and vector were then mixed at 0:1, 3:1, and 5:1 insert:vector ratios and ligated using T4 DNA ligase in 1X T4 DNA ligase buffer (NEB). Ligation was incubated overnight at 23 °C then transformed into chemically-competent XL1-blue cells and plated on 50 µg/mL kanamycin agar plates. Cells were grown at 37 °C overnight and colonies were picked for culturing and Sanger sequencing (QuintaraBiosciences).

Site-directed mutagenesis was performed via Quikchange using complementary oligonucleotide primers (IDT) that change a single codon from M to X shown in bold. Forward and reverse reactions were carried out in separate PCR tubes for 5 cycles, then combined 1:1 for 13 additional cycles. Methylated DNA was digested via addition of 20 U DpnI (NEB) and incubation at 37 °C for 1 hour. 5 µL of PCR reaction was directly transformed into chemically-competent XL1-blue cells and plated on 50 µg/mL kanamycin agar plates. Cells were grown at 37 °C overnight and colonies were picked for culturing and Sanger sequencing (QuintaraBiosciences).

### FLAG-BPHL cloning and M69 mutagenesis

WT Sequence (hBPHL, Uniprot ID: Q86WA6)

MVAVLGGRGVLRLRLLLSALKPGIHVPRAGPAAAFGTSVTSAKVAVNGVQLHYQQTGEGDHAVLLLPGMLGSGETDFGPQLKNLNKKLFTVVAWDPRGYGHSRPPDRDFPADFFERDAKDAVDLMKALKFKKVSLLGWSDGGITALIAAAKYPSYIHKMVIWGANAYVTDEDSMIYEGIRDVSKWSERTRKPLEALYGYDYFARTCEKWVDGIRQFKHLPDGNICRHLLPRVQCPALIVHGEKDPLVPRFHADFIHKHVKGSRLHLMPEGKHNLHLRFADEFNKLAEDFLQ

### Cloned Sequence (FLAG-BPHL)

<u>MGDYKDDDDKENLYFQSGGSGG</u>MVAVLGGRGVLRLRLLLSALKPGIHVPRAGPAAAFGTSVTSAKVAVNGVQLHYQQTGEGDHAVLLLPG**M**LGSGETDFGPQLKNLNKKLFTVVAWDPRGYGHSRPPDRDFPADFFERDAKDAVDLMKALKFKKVSLLGWSDGGITALIAAAKYPSYIHKMVIWGANAYVTDEDSMIYEGIRDVSKWSERTRKPLEALYGYDYFARTCEKWVDGIRQFKHLPDGNICRHLLPRVQCPALIVHGEKDPLVPRFHADFIHKHVKGSRLHLMPEGKHNLHLRFADEFNKLAEDFLQ*

<u>5’ – BPHL HindIII FOR– 3’</u>

CGACAAAGCTTGCCACCGCCACCATGGGAGACTATAAG

<u>5’ – BPHL XbaI REV– 3’</u>

AATATTCTAGACTCGAGTCACTGAAGAAAATCTTCAGCC

<u>5’ – BPHL M69A FOR– 3’</u>

CCTTTTGTTGCCGGGC**GC**GCTGGGGAGCGGGG

<u>5’ – BPHL M69A REV– 3’</u>

CCCCGCTCCCCAGC**GC**GCCCGGCAACAAAAGG

### SOD1 cloning and K76 mutagenesis

WT Sequence (SODC, Uniprot ID: P00441)

MATKAVCVLKGDGPVQGIINFEQKESNGPVKVWGSIKGLTEGLHGFHVHEFGDNTAGCTSAGPHFNPLSRKHGGPKDEERHVGDLGNVTADKDGVADVSIEDSVISLSGDHCIIGRTLVVHEKADDLGKGGNEESTKTGNAGSRLACGVIGIAQ*

Cloned sequence (V5-SOD1)

<u>MGKPIPNPLLGLDSTGGSSGG</u>ATKAVCVLKGDGPVQGIINFEQKESNGPVKVWGSIKGLTEGLHGFHVHEFGDNTAGCTSAGPHFNPLSRKHGGPKDEERHVGDLGNVTADKDGVADVSIEDSVISLSGDHCIIGRTLVVHEKADDLGKGGNEESTKTGNAGSRLACGVIGIAQ*

5’ – SOD1 HindIII FOR– 3’

CGACACAAAGCTTGCCACCGCCACCATGGGGAAACCGATACCG

5’ – SOD1 XbaI REV – 3’

TATATATCTAGACTCGAGTTACTGAGCTATACCTATTACCCCAC

5’ – SOD1 K76R FOR– 3’

CACGGTGGGCCTA**G**AGACGAAGAGCGCCAC

5’ – SOD1 K76R REV– 3’

GTGGCGCTCTTCGTCT**C**TAGGCCCACCGTG

Molecular cloning was performed using a pcDNA3.1(+) vector via traditional double-restriction digest using a codon-optimized synthetic gene-block of SOD1 for mammalian expression (IDT) and overhang oligonucleotide primers (IDT). Insert PCR product was amplified and purified using QIAquick PCR purification (Qiagen). Both insert and vector were then double-restriction digested using HindIII and XbaI endonucleases (NEB). Plasmid was gel-purified using QIAquick gel purification (Qiagen) and insert was PCR purified. The digested insert and vector were then mixed at 0:1, 3:1, and 5:1 insert:vector ratios and ligated using T4 DNA ligase in 1X T4 DNA ligase buffer (NEB). Ligation was incubated overnight at 23 °C then transformed into chemically- competent XL1-blue cells and plated on 100 μg/mL ampicillin agar plates. Cells were grown at 37 °C overnight and colonies were picked for culturing and Sanger sequencing (QuintaraBiosciences). Plasmids were bacterial endotoxing purified using Endofree purification kit (Qiagen) prior to mammalian transfection.

Site-directed mutagenesis was performed via Quikchange using complementary oligonucleotide primers (IDT) that change a single codon from M to L shown in bold. Forward and reverse reactions were carried out in separate PCR tubes for 5 cycles, then combined 1:1 for 13 additional cycles. Methylated DNA was digested via addition of 20 U DpnI (NEB) and incubation at 37 °C for 1 hour. 5 μL of PCR reaction was directly transformed into chemically-competent XL1-blue cells and plated on 50 μg/mL kanamycin agar plates. Cells were grown at 37 °C overnight and colonies were picked for culturing and Sanger sequencing (QuintaraBiosciences).

### Recombinant protein expression and purification

pET24a-rhBPHL and pGro7-GroEL/GroES were co-transformed into a chemically-competent BL21 (DE3) expression host. Following kanamycin (Kan) and chloramphenicol (Cm) selection on agar plates (50 µg/mL Kan, 25 µg/mL Cm), three separate 5 mL starter cultures were grown in LB media containing Kan/Cm at 37 °C overnight. These cultures were then inoculated into three 1 L cultures of LB media containing Kan/Cm and grown at 37 °C to OD_600_ = 0.6 then induced with 0.5 mM IPTG and 0.1% (w/v) L-arabinose. Protein was expressed for 18 hours at 37 °C then pelleted by centrifugation at 5,000 xg, 4 °C for 25 minutes. Overexpressed pellets were then flash-frozen in liquid nitrogen and stored at -80 °C. Protein pellets were thawed on ice then resuspended in 35 mL lysis buffer containing EDTA-free protease inhibitors (Roche). Cells were kept on ice and lysed via sonication (5 s on, 20 s off, 40% A, 6 pulses) then centrifuged at 15,000 xg, 4 °C for 25 minutes. Lysate was further clarified by 0.45 µm syringe filtration. Clarified lysate was then loaded onto 2 mL pre-equilibrated Ni-NTA resin (ThermoFisher Scientific; 88222) and batch incubated at 4 °C for 1 hour. Following incubation, the resin was added to a gravity filter and bound protein was washed by addition of 3 × 10 mL wash buffer, then eluted with 2 x 3 mL elution buffer. Protein was then dialyzed against 3 L SEC buffer at 4 °C for 4 hours before being loaded a HiLoad 16/600 Superdex 75 pg size-exclusion column and separated at a 0.25 mL/min flow rate. Fractions in the peak after GroEL/ES elution were pooled, concentrated, and quantified by A280.

<u>Buffers</u>

Lysis – 20 mM HEPES, 300 mM NaCl, 5 mM imidazole, 5% glycerol, pH 8.0

Wash – 20 mM HEPES, 300 mM NaCl, 15 mM imidazole, 5% glycerol, pH 8.0

Elution – 20 mM HEPES, 300 mM NaCl, 300 mM imidazole, 5% glycerol, pH 8.0

SEC – 20 mM HEPES, 150 mM NaCl, 5% glycerol, pH 8.0

### Treatment of FLAG-BPHL with ChURRO-2 enantiomers for Strep-IP

Adapted from Zanon et al.^57^ Transfected HEK-293T cells bearing pcDNA3.1(+)-FLAG-BPHL were prepared as described below (see FLAG-BPHL transfection). Cells were resuspended in 200 μL ice-cold PBS + 1% Triton X-100 (v/v) containing EDTA-free protease inhibitors (Roche). Cells were lysed at 4 °C for 30 minutes while rotating. Lysate was clarified by centrifugation at 12,000 xg at 4 °C for 15 minutes. The supernatant was then transferred to a fresh prechilled 1.7 mL microcentrifuge tube. Protein concentration was normalized to 2.0 mg/mL via BCA assay (Thermo; 23225). 100 µL of this sample was then aliquoted into separate 2.0 mL tubes and allowed to equilibrate to 23 °C. Samples were treated with 1 µL of either (*R*)- or (*S*)-ChURRO-2 at the desired concentration in DMSO and incubated at 23 °C for 1 hour. 2 µL of DTB-N_3_ (1.2 equiv.) was added to each sample with 6 µL TBTA (stock in 1:4 DMSO:*^t^*BuOH, 100 µM), 2 µL TCEP (1 mM), and 2 µL CuSO_4_ (1 mM).

Proteins were unfolded by addition of 20 µL 1.2% SDS in PBS (0.1% w/v) and CuAAC was allowed to proceed at 23 °C for 1 hour with rocking. Proteins were then precipitated by addition of 4 volumes of MeOH:CHCl_3_ (4:1.5), vortexed, and centrifuged at 12,000 xg for 15 minutes. The collected protein pellet was washed 1X with 500 µL chilled (-20 °C) MeOH via probe-tip sonication (5 s on, 20 s off, 20% A, 4-6 pulses) and resuspended in 100 µL 1.2% SDS/PBS via probe-tip sonication (5 s on, 20 s off, 20% A, 6-8 pulses).

### General Strep-IP protocol

High-capacity streptavidin agarose beads (50 µL per sample) (Thermo Fisher Scientific; 20357) were washed 3X in PBS then transferred to a fresh 2.0 mL microcentrifuge tube containing 1.1 mL PBS. The reconstituted protein samples (100 µL) were then transferred to tubes containing streptavidin beads and allowed to incubate at 23 °C for 2 hours. The samples were centrifuged at 1,400 xg for 3 minutes and the supernatant was removed. The beads were then washed with 500 µL PBS (5X) and 500 µL MQ (5X) in micro bio-spin columns (Bio-Rad; 7326204) on a vacuum manifold. With the final wash, samples were transferred to a fresh 1.7 mL microcentrifuge tube, centrifuged at 1,400 xg for 3 minutes, and the supernatant was removed. Beads were resuspended in 100 µL of a premixed 1X Laemmli’s buffer (Bio-Rad; 1610747) + 10% BME diluted with PBS and boiled at 95 °C for 8 minutes. 20 µL of bead slurry was loaded per well and separated on precast 4–20% Novex Tris-Gly SDS-PAGE gels (Invitrogen) run at 160 V for 70 minutes. Proteins were then electro-transferred to a PVDF membrane (25 V/2.5 A for 30 minutes). Membranes were then blocked in a solution of TBST + 5% BSA (w/v) and rocked at 23 °C for 1 hour. Membranes were washed 3X in TBST for 5 minutes each while rocking and cut using scissors along the protein ladder for separate antibody incubation. Membranes were then blotted with primary mouse anti-FLAG M2 (Sigma; F1804), rabbit anti-BPHL (Sigma; HPA036752), rabbit anti-β actin (CST; 4970), or rabbit anti-GAPDH (CST; 5174) (1:1000 TBST + 5% BSA) at 4 °C overnight. The following morning membranes were washed 3X in TBST for 5 minutes then incubated with either anti-mouse IgG HRP conjugated secondary (CST; 7076) or anti-rabbit IgG HRP conjugated secondary (CST; 7074) (1:3000 TBST) for 2 hours at 23 °C. Blots were quickly washed 3X with TBST prior to incubation with ECL western blotting substrates (Sigma; 34580) for 5 minutes and imaged with ChemiDoc MP. Band intensities were quantified via densitometry when applicable (ImageJ).

### Co-IP of BPHL and Msr proteins

Transfected HEK-293T cells bearing pcDNA3.1(+)-FLAG-BPHL were prepared as described below (see Co-IP transfection and uncoupler treatment). Cells were resuspended in 200 μL ice-cold PBS + 1% Triton X-100 (v/v) containing EDTA-free protease inhibitors (Roche). Cells were lysed at 4 °C for 30 minutes while rotating. Lysate was clarified by centrifugation at 12,000 xg at 4 °C for 15 minutes. The supernatant was then transferred to a fresh prechilled 1.7 mL microcentrifuge tube. Protein concentration was normalized to 4.0 mg/mL via BCA assay (Thermo; 23225). 100 µL of input sample was boiled with 4X Laemmli’s buffer (Bio-Rad; 1610747) + 10% BME at 95 °C for 8 minutes and stored at -20 °C for downstream analysis. Anti-FLAG agarose beads (50 µL per sample) (Sigma; A2220) were aliquoted into 1.7 mL microcentrifuge tubes and washed 3X in 500 µL PBS by centrifugation at 1,400 xg for 3 minutes and removal of the supernatant. The resulting beads were then resuspended in 300 µL PBS and 100 µL of protein sample was added for a final protein concentration of 1.0 mg/mL. The samples were incubated for 2 hours at 23 °C while rocking then washed 3X with 500 µL PBS. Beads were resuspended in 100 µL of a premixed 1X Laemmli’s buffer (Bio-Rad; 1610747) + 10% BME diluted with PBS and boiled at 95 °C for 8 minutes. 20 µL of bead slurry was loaded per well and separated on precast 4–20% Novex Tris-Gly SDS-PAGE gels (Invitrogen) run at 160 V for 70 minutes. Proteins were then electro-transferred to a PVDF membrane (25 V/2.5 A for 30 minutes). Membranes were then blocked in a solution of TBST + 5% BSA (w/v) and rocked at 23 °C for 1 hour. Membranes were washed 3X in TBST for 5 minutes each while rocking and cut using scissors along the protein ladder for separate antibody incubation. Membranes were then blotted with primary mouse anti-FLAG M2 (Sigma; F1804), rabbit anti-MsrA (Invitrogen; PA5-88562), rabbit anti-MsrB2 (abcam; ab229940), or rabbit anti-β actin (CST; 4970) (1:1000 TBST + 5% BSA) at 4 °C overnight. The following morning membranes were washed 3X in TBST for 5 minutes then incubated with either anti-mouse IgG HRP conjugated secondary (CST; 7076) or anti-rabbit IgG HRP conjugated secondary (CST; 7074) (1:3000 TBST) for 2 hours at 23 °C. Blots were quickly washed 3X with TBST prior to incubation with ECL western blotting substrates (Sigma; 34580) for 5 minutes and imaged with ChemiDoc MP.

### Cell line maintenance

Cells were maintained by the UC Berkeley Tissue Culture Facility. All cells were maintained as a monolayer in exponential growth at 37 °C in a 5% CO_2_ atmosphere. HEK-293T cells were maintained in Dulbecco’s Modified Eagle Medium + GlutaMAX (DMEM, Gibco) supplemented with 10% (v/v) fetal bovine serum (FBS, Seradigm), referred to below as complete media.

### Cytotoxicity measurements of ChURRO-2

Poly-lysine treated HEK-293T cells were grown to 80% confluency in 100 µL complete media in a 96-well plate at 37 °C, 5% CO_2_. The media was aspirated and replaced in either 100 µL of complete media or 100 µL of media containing no FBS. Cells were then treated with 1 µL of a stock of either (±)-, (*R*)-, or (*S*)-ChURRO-2 in DMSO for a final concentration of 500, 250, 100, 50, 25, 10, 1, or 0 µM in triplicate and incubated at 37 °C, 5% CO_2_ for 1 hour. Following incubation, the media was then aspirated and replaced with 100 µL fresh complete media. Cell viability was then assessed by CCK-8 assay (Dojindo; CK04) according to the manufacturer’s protocol.

### Live cell and lysate ChURRO-2 labeling comparison by Gel-ABPP

HEK-293T cells were grown to 80% confluency in 3 mL complete media in a 6-well plate at 37 °C, 5% CO_2_. The media was aspirated and replaced in either 3 mL of complete media or 3 mL of media containing no FBS. Cells were then treated with 30 µL of a stock of either (±)-, (*R*)-or (*S*)-ChURRO-2, or vehicle control in DMSO for a final concentration of 50 µM and incubated at 37 °C, 5% CO_2_ for 1 hour. Cells were then harvested by trypsinization and washed 1X with 1 mL of ice-cold PBS, pelleted at 300 xg, and flash-frozen in LN_2_. The same day, cells were resuspended in 100 μL ice-cold PBS + 1% Triton X-100 (v/v) containing EDTA-free protease inhibitors (Roche). Cells were lysed at 4 °C for 30 minutes while rotating. Lysate was clarified by centrifugation at 12,000 xg at 4 °C for 15 minutes. The supernatant was then transferred to a fresh prechilled 1.7 mL microcentrifuge tube. Protein concentration was normalized to 1.0 mg/mL via BCA assay (Thermo; 23225). 100 µL of these samples were then aliquoted into separate 1.7 mL tubes and allowed to equilibrate to 23 °C. The vehicle control lysate samples were then treated with 1 µL of either (±)-, (*R*)-, or (*S*)-ChURRO-2 in DMSO for a final concentration of 50 µM. Samples were allowed to incubate at 23 °C for 1 hour. Excess oxaziridine was then quenched by addition of 1 µL *N*-acetyl methionine (100 µM) and incubation at 23 °C for 30 minutes. To all samples, 2 µL of Alexafluor488-N_3_ (1.2 equiv.) was added to each sample with 6 µL TBTA (stock in 1:4 DMSO:*^t^*BuOH, 100 µM), 2 µL TCEP (1 mM), and 2 µL CuSO_4_ (1 mM).

Proteins were unfolded by addition of 10 µL 1.2% SDS in PBS (0.1% w/v) and CuAAC was allowed to proceed at 23 °C for 1 hour with rocking. Then, 40 µL of 4X Laemmli’s buffer (Bio-Rad; 1610747) + 10% BME was added, and samples were boiled at 95 °C for 8 minutes. 30 µL of sample was loaded per well and separated on precast 4–20% Novex Tris-Gly SDS-PAGE gels (Invitrogen) run at 160 V for 70 minutes. The gel was fixed in MQ water containing 10% acetic acid, 30% EtOH overnight then visualized using ChemiDoc MP (Bio-Rad) for measuring in-gel fluorescence (ex = 499 nm, em = 520 nm). The gel was rehydrated in MQ water for 1 hour then total protein level was assessed by coomassie brilliant blue staining and visualized under brightfield.

### Dot blot assessment of ChURRO-2 labeling

HEK-293T cells were grown to 80% confluency in 3 mL complete media in a 6-well plate at 37 °C, 5% CO_2_. The media was aspirated and replaced in 3 mL of media containing no FBS. Cells were then treated with 30 µL of a stock of either (±)-, (*R*)- or (*S*)-ChURRO-2, or vehicle control in DMSO for a final concentration of 50 µM and incubated at 37 °C, 5% CO_2_ for 1 hour. Cells were then harvested by trypsinization and washed 1X with 1 mL of ice-cold PBS, pelleted at 300 xg, and flash-frozen in LN_2_. The same day, cells were resuspended in 100 μL ice-cold PBS + 1% Triton X-100 (v/v) containing EDTA-free protease inhibitors (Roche). Cells were lysed at 4 °C for 30 minutes while rotating. Lysate was clarified by centrifugation at 12,000 xg at 4 °C for 15 minutes. The pellet was collected for plasma membrane analysis and resuspended in 100 µL 1.2% SDS/PBS via probe-tip sonication (5 s on, 20 s off, 20% A, 6-8 pulses). Protein concentration for all samples were normalized to 1.0 mg/mL via BCA assay (Thermo; 23225). 100 µL of these samples were then aliquoted into separate 1.7 mL tubes and allowed to equilibrate to 23 °C. The vehicle control lysate samples were then treated with 1 µL of either (±)-, (*R*)- or (*S*)-ChURRO-2 in DMSO for a final concentration of 50 µM. Samples were allowed to incubate at 23 °C for 1 hour. Excess oxaziridine was then quenched by addition of 1 µL *N*-acetyl methionine (100 µM) and incubation at 23 °C for 30 minutes. To all samples, 2 µL of DTB-N_3_ (1.2 equiv.) was added to each sample with 6 µL TBTA (stock in 1:4 DMSO:*^t^*BuOH, 100 µM), 2 µL TCEP (1 mM), and 2 µL CuSO_4_ (1 mM). Proteins were unfolded by addition of 20 µL 1.2% SDS in PBS (0.1% w/v) and CuAAC was allowed to proceed at 23 °C for 1 hour with rocking. Proteins were then precipitated by addition of 4 volumes of MeOH:CHCl_3_ (4:1.5), vortexed, and centrifuged at 12,000 xg for 15 minutes. The collected protein pellet was washed 1X with 500 µL chilled (-20 °C) MeOH via probe-tip sonication (5 s on, 20 s off, 20% A, 4-6 pulses) and resuspended in 100 µL 1.2% SDS/PBS via probe sonication (5 s on, 20 s off, 20% A, 6-8 pulses). Dot blot assessment was conducted using Bio-Dot apparatus (Bio-Rad; 1706545). Briefly, a nitrocellulose membrane was rehydrated by rocking in PBS for 10 mins. The membrane was then added to the Bio-Dot apparatus connected to house vacuum, and 10 µL of sample was then added to each well. Vacuum drying was conducted for 1 hour. Following drying, the membrane was washed 3X in TBST for 5 minutes then blocked in a solution of TBST + 5% BSA (w/v) and rocked at 23 °C for 1 hour. Membrane was then washed 3X in TBST for 5 minutes each while rocking. Membrane was then blotted with primary rabbit anti-TRPC3 (CST; 77934) at 4 °C overnight. The following morning the membrane was washed 3X in TBST for 5 minutes then incubated with anti-rabbit IgG HRP conjugated secondary (CST; 7074) (1:3000 TBST) for 2 hours at 23 °C. Blots were quickly washed 3X with TBST prior to incubation with ECL western blotting substrates (Sigma; 34580) for 5 minutes and imaged with ChemiDoc MP. The membrane was then stripped by incubation with Restore™ PLUS Western Blot Stripping Buffer (Thermo Fisher Scientific; 46430) and reblotted with primary Strep-HRP conjugate (Jackson ImmunoResearch; AB_2337238) (1:10,000 TBST) for 2 hours at 23 °C. Blots were quickly washed 3X with TBST prior to incubation with ECL western blotting substrates (Sigma; 34580) for 5 minutes and imaged with ChemiDoc MP. Finally, membrane was incubated with SafeStain (Invitrogen; LC6060) for total protein loading imaging and imaged under white light with ChemiDoc MP. Band intensities were quantified via densitometry (ImageJ).

### FLAG-BPHL transfection

HEK-293T cells were grown to 70% confluency in 10 mL complete media in a 10 cm plate at 37 °C, 5% CO_2_. Transfection was then performed as per Lipofectamine 3000 protocol (Invitrogen). Briefly, 25 µg of pcDNA3.1(+)-FLAG-BPHL expression construct was introduced at 1:1 transfection reagent:DNA ratio. The lipid-DNA complex was incubated for 30 minutes at 23 °C in 2 mL Opti-MEM media (Gibco). Then, 2 mL complex was added to 8 mL complete media. 10 mL of this solution was then added to each 10 cm plate. Cells were incubated for 24 hours at 37 °C, 5% CO_2_. The media was then aspirated, and cells were harvested by trypsinization. Cells were washed 1X with 10 mL of ice-cold PBS, pelleted at 300 xg, and flash-frozen in LN_2_ for future analysis.

### Co-IP transfection and uncoupler treatment

HEK-293T cells were grown to 70% confluency in 10 mL complete media in a 10 cm plate at 37 °C, 5% CO_2_. Transfection was then performed as per Lipofectamine 3000 protocol (Invitrogen). Briefly, 25 µg of pcDNA3.1(+)-FLAG-BPHL expression construct was introduced at 1:1 transfection reagent:DNA ratio. The lipid-DNA complex was incubated for 30 minutes at 23 °C in 2 mL Opti-MEM media (Gibco). Then, 2 mL complex was added to 8 mL complete media. 10 mL of this solution was then added to each 10 cm plate. Cells were incubated for 24 hours at 37 °C, 5 % CO_2_. The media was then aspirated, and fresh complete media containing vehicle (DMSO), 100 µM paraquat, or 2.5 µM FCCP was added. Cells were incubated for 18 hours at 37 °C, 5 % CO_2_. The following day, cells were harvested by trypsinization. Cells were washed 1X with 10 mL of ice-cold PBS, pelleted at 300 xg, and flash-frozen in LN_2_ for future analysis.

### CRISPR KO of MsrA and MsrB2

Producer HEK-293T cells were grown to 70% confluency in 10 mL complete media in a 10 cm plate at 37 °C, 5% CO2. Producer cells were transfected using 1 µg of lentiviral plasmids psPAX2 and pMD2.G each as well as 3 µg of plasmid containing a lentiCas9-bleomycin cassette following the TransIT-293 transfection protocol (Mirusbio; MIR 2700). After 48 hours, the viral media was collected, filtered using a 0.45 um syringe filter, and 5 mL was mixed 1:1 with media containing 8 µg/mL polybrene. A separate 10 cm plate containing 70% confluent HEK-293T cells was transduced by switching the media with the viral mixture and incubated for 24 hours. After 3 days, selection for stable Cas9 expression was performed by 250 µg/mL Zeocin selection for an additional 10 days to generate Cas9-293T cells. MsrA and MsrB2 sgRNAs were designed using Feng Zhang’s protocol^58^ from a panel of 20 gene-specific sequences to target two discrete loci following PAM constraints. These sgRNAs were inserted into a lentiCRISPR v2 vector by PCR amplification and restriction digest cloning using BsmBI overhangs designed into the oligos below.

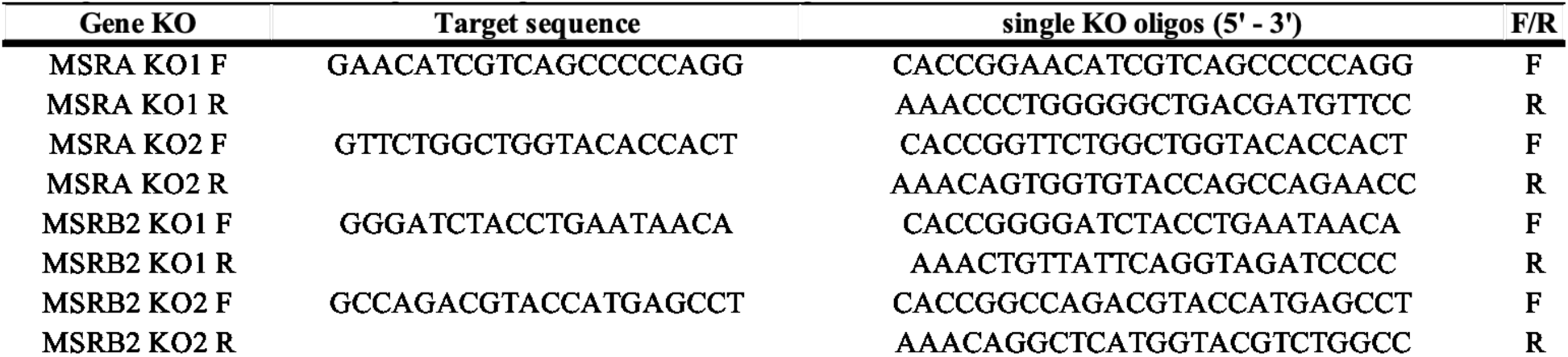

A separate batch of producer HEK-293T cells were grown to 70% confluency in 10 mL complete media in a 10 cm plate at 37 °C, 5% CO_2_. Separate producer cells were transfected using 1 µg of lentiviral plasmids psPAX2 and pMD2.G each as well as 3 µg of plasmid containing lentiCRISPR v2-sgRNA plasmid bearing either MsrA g1/g2, or MsrB2 g1/g2 following the TransIT-293 transfection protocol (Mirusbio; MIR 2700). After 48 hours, the viral media was collected, filtered using a 0.45 um syringe filter, and 5 mL was mixed 1:1 with a 8 µg/mL polybrene solution. A separate 10 cm plate containing 70% confluent Cas9-293T cells was transduced by switching the media with the viral mixture and incubated for 24 hours. Selection for stable Cas9 expression was performed after 3 days by 1 µg/mL puromycin selection for an additional 10 days to generate MsrA and MsrB2 KO lines.

### siRNA knock-down for *N*-homocysteinylation

HEK-293T cells were grown to 70% confluency in 10 mL complete media in a 10 cm plate at 37 °C, 5% CO_2_. Transfection was then performed as per Lipofectamine RNAiMax protocol. Briefly, 500 pmol of desired siRNA, siControl (Thermo; 4390843), siMsrA (Thermo; AM16708), or siMsrB2 (Thermo; AM16708), was introduced to 25 µL lipid. The lipid-siRNA complex was incubated for 30 minutes at 23 °C in 2 mL Opti-MEM media (Gibco). Then, 2 mL complex was added to 8 mL complete media. 10 mL of this solution was then added to each 10 cm plate containing a final siRNA concentration of 50 nM. Cells were incubated for 24 hours at 37 °C, 5 % CO_2_. The media was then aspirated, and fresh complete media containing a combination of 500 µM HCTL and either vehicle (DMSO) or 100 µM paraquat was added. Cells were incubated for 18 hours at 37 °C, 5 % CO_2_. The following day, cells were harvested by trypsinization. Cells were washed 1X with 10 mL of ice-cold PBS, pelleted at 300 xg, and flash-frozen in LN_2_ for future analysis.

### siRNA knock-down for immunoblotting

HEK-293T cells were grown to 70% confluency in 3 mL complete media in a 6-well plate at 37 °C, 5% CO_2_. Transfection was then performed as per Lipofectamine RNAiMax protocol. Briefly, 150 pmol of desired siRNA, siControl (Thermo; 4390843), siMsrA (Thermo; AM16708), or siMsrB2 (Thermo; AM16708), was introduced to 7.5 µL lipid. The lipid-siRNA complex was incubated for 30 minutes at 23 °C in 500 µL Opti-MEM media (Gibco). Then, 500 µL complex was added to 2.5 mL complete media. 10 mL of this solution was then added to each 10 cm plate containing a final siRNA concentration of 50 nM. Cells were incubated for 24 hours at 37°C, 5 % CO_2_. The following day, cells were harvested by trypsinization. Cells were washed 1X with 1 mL of ice-cold PBS, pelleted at 300 xg, and flash-frozen in LN_2_ for future analysis.

### siRNA knock-down for proteomics

HEK-293T cells were grown to 70% confluency in 25 mL complete media in a 15 cm plate at 37 °C, 5% CO_2_. Transfection was then performed as per Lipofectamine RNAiMax protocol. Briefly, 1250 pmol of desired siRNA, siControl (Thermo; 4390843), siMsrA (Thermo; AM16708), or siMsrB2 (Thermo; AM16708), was introduced to 62.5 µL lipid. The lipid-siRNA complex was incubated for 30 minutes at 23 °C in 5 mL Opti-MEM media (Gibco). Then, 5 mL complex was added to 20 mL complete media. 25 mL of this solution was then added to each 15 cm plate containing a final siRNA concentration of 50 nM. Cells were incubated for 24 hours at 37°C, 5 % CO_2_. The media was then aspirated, and fresh complete media containing 100 µM paraquat was added. Cells were incubated for 18 hours at 37 °C, 5 % CO_2_. The following day, cells were harvested by trypsinization. Cells were washed 2X with 10 mL of ice-cold PBS, pelleted at 300 xg, and flash-frozen in LN_2_ for future analysis.

### Determination of Msr knockdown out efficiency by immunoblotting

CRISPR KO or transfected HEK-293T cells bearing the corresponding siRNA were prepared as described above. Cells were resuspended in 200 μL ice-cold PBS + 1% Triton X-100 (v/v) containing EDTA-free protease inhibitors (Roche). Cells were lysed at 4 °C for 30 minutes while rotating. Lysate was clarified by centrifugation at 12,000 xg at 4 °C for 15 minutes. The supernatant was then transferred to a fresh prechilled 1.7 mL microcentrifuge tube. Protein concentration was normalized to 2.0 mg/mL via BCA assay (Thermo; 23225). 100 µL of sample was boiled with 4X Laemmli’s buffer (Bio-Rad; 1610747) + 10% BME diluted at 95 °C for 8 minutes. Approximately 30 µg of protein sample was loaded per well and separated on precast 4–20% Novex Tris-Gly SDS-PAGE gels (Invitrogen) run at 160 V for 70 minutes. Proteins were then electro-transferred to a PVDF membrane (25 V/2.5 A for 30 minutes). Membranes were then blocked in a solution of TBST + 5% BSA (w/v) and rocked at 23 °C for 1 hour. Membranes were washed 3X in TBST for 5 minutes each while rocking and cut using scissors along the protein ladder for separate antibody incubation. Membranes were then blotted with primary rabbit anti-BPHL (Sigma; HPA036752), rabbit anti-MsrA (Invitrogen; PA5-88562), rabbit anti-MsrB2 (abcam; ab229940), or rabbit anti-β actin (CST; 4970) (1:1000 TBST + 5% BSA) at 4 °C overnight. The following morning membranes were washed 3X in TBST for 5 minutes then incubated with anti-rabbit IgG HRP conjugated secondary (CST; 7074) (1:3000 TBST) for 2 hours at 23 °C. Blots were quickly washed 3X with TBST prior to incubation with ECL western blotting substrates (Sigma; 34580) for 5 minutes and imaged with ChemiDoc MP.

### Animal raising and feeding

All animal experiments were approved and performed under the guidelines and ethical regulations established by the UC Berkeley Animal Care and Use Committee. Male wildtype C57BL/6J mice (JAX, stock #000664) were fed a regular chow diet (LabDiet 5053) until 6-8 weeks of age under ambient temperature at 23°C. Mice were fasted for 4 – 5 hours prior to sacrificing and tissue harvesting. All animal studies were performed using age-matched male mice.

### Tissue homogenization

All steps and centrifugations were carried out at 4 °C. Mouse livers were thawed on ice and rinsed 2X with 5 mL of ice-cold PBS. Livers were cut into small pieces (size of a grain of rice) using a razor blade prior to homogenization. Chopped liver pieces were transferred to a glass homogenizer on ice and homogenized with 6 – 8 strokes of the glass pestle in 10 mL/gram tissue ice-cold PBS. Tissue homogenate was then sonicated via probe-tip sonication (5 s pulse, 20 s off, 40% A, 4-6 pulses) then strained through a 100 µM cell strainer into a 50 mL conical tube and centrifuged at 12,000 xg for 20 minutes. The resulting top fatty layer was removed by pipetting and the supernatant was collected.

### Mitochondrial purification

All steps and centrifugations were carried out at 4 °C. Mouse livers were thawed on ice and rinsed twice with 5 mL of ice-cold PBS. 3 mouse livers were cut into small pieces (size of a grain of rice) using a razor blade prior to homogenization. Chopped liver pieces were transferred to a glass homogenizer on ice and homogenized with 6 – 8 strokes of the glass pestle in 10 mL/gram tissue ice-cold Buffer A. Tissue homogenate was then strained through a 100 µM cell strainer into separate 50 mL conical tube and balanced by dilution with Buffer A. Samples were centrifuged for 30 minutes at 600 xg to remove nuclear components. The supernatant was collected and centrifuged an additional 10 minutes at 15,000 xg to yield a pellet of crude mitochondria. The crude mitochondria pellet was resuspended in 18 mL Buffer B and centrifuged for 30 minutes at 15,000 xg. The supernatant was discarded, and the pellet was resuspended in 9 mL of Buffer C. To three separate polycarbonate ultracentrifuge tubes (24 mL size) was added a 5 mL layer of Buffer D followed by careful addition (add to the side of tube and pipette extremely gently) of 9 mL Buffer E. Then, 3 mL of resuspended crude mitochondrial pellet was carefully added to the top layer of each Percoll gradient. The triphasic solutions were centrifuged with a SW 32Ti swinging-bucket rotor (Beckman; 369650) using an Optima XPN-100-IVD ultracentrifuge (Beckman; B10053) at 41,000 xg for 30 minutes. After centrifugation, enriched mitochondria were collected from the interface formed between the 26% and 60% Percoll and transferred to a 15 mL conical tube. Excess Percoll was removed from the mitochondrial fraction by resuspension in 10 mL of ice-cold PBS followed by centrifugation at 7,000 xg for 10 minutes. The resulting pellet of enriched mitochondria was then flash-frozen in LN_2_ for future analysis.

<u>Buffers</u>

A – 225 mM mannitol, 75 mM sucrose, 0.1 mM EGTA, 1 mg/mL fatty acid-free BSA, 10 mM HEPES, pH 7.4

B – 225 mM mannitol, 75 mM sucrose, 0.1 mM EGTA, 10 mM HEPES, pH 7.4

C – 395 mM sucrose, 0.1 mM EGTA, 10 mM HEPES, pH 7.4

D – 1.28 M sucrose, 0.4 mM EGTA, 40 mM HEPES, 60% Percoll (v/v), pH 7.4

E – 1.28 M sucrose, 0.4 mM EGTA, 40 mM HEPES, 26% Percoll (v/v), pH 7.4

### Determination of mitochondrial enrichment by immunoblotting

Mitochondrial lysate was normalized to 4.0 mg/mL in PBS. Samples were diluted with 4X Laemmli’s buffer (Bio-Rad Laboratories, Inc.; 1610747) + 10% BME, boiled at 95 °C for 8 minutes, and loaded at 40 µg per lane on a 4-20% Tris-Gly SDS-PAGE gel. The gel was run at 160 V for 80 minutes and semi-dry electrotransferred to a PVDF membrane at 25 V, 2.5 Å, for 10 minutes. Blots were blocked with 5% BSA/TBST for 1 hour, then washed 2X with TBST for 5 minutes, and cut for incubation with separate antibodies. Antibodies used were rabbit anti-VDAC (CST; 4661), rabbit anti-PDI (CST; 3501), rabbit anti-RCAS1 (CST; 12290), rabbit anti-GAPDH (CST; 5714), and rabbit anti-Histone H3 (CST; 4499). All antibodies were diluted at 1:1000 in 5% BSA/TBST at 4 °C overnight. The next morning the blots were washed 3X with TBST prior to incubation with anti-rabbit IgG HRP conjugated secondary (CST; 7074) (1:3000 TBST) for 2 hours at 23 °C. Blots were quickly washed 3X with TBST prior to incubation with ECL western blotting substrates (Promega; W1001) for 1 minute and imaging with ChemiDoc MP.

### Alpha 1 antitrypsin, GST, and BPHL ChURRO shotgun proteomics

Proteins (Alpha 1 antitrypsin: Sigma, Cat. 178251; Glutathione S transferase: abcam, Cat. Ab89494) were diluted to 10 µM. Recombinant human WT BPHL was diluted to 0.1mg/ml (3mM). To 100 µL of protein was added 1 µL of a stock of either (*R*)-, (*S*)-, or racemic ChURRO-2 in DMSO for a final concentration of 200 µM or 30 mM for human BPHL. Samples were allowed to incubate for 1 hour at 23 °C. While samples were incubating, 10 µL each of magnetic E3 (Cytiva; 44152105050250) and E7 (Cytiva; 24152105050250) SP3 beads were mixed 1:1 and washed 3X with 1 mL MQ using a magnetic rack. The washed beads were then resuspended in 20 µL of MQ and added to each protein sample. The samples were allowed to incubate at 23 °C for 5 minutes with shaking (1000 rpm). Protein binding was induced by addition of 800 µL absolute ethanol followed by incubation at 23 °C for 5 minutes with shaking (1000 rpm). The beads were then placed on a magnetic rack and allowed to settle. The supernatant was removed, and the beads were then resuspended and 3X with 400 µL of 80% (v/v) ethanol. The beads were then resuspended in 200 µL of 8 M urea in PBS and incubated with 10 µL TCEP (10 mM) at 65 °C for 15 minutes. Then, 10 µL of iodoacetamide (20 mM) was added and incubated at 37 °C for 30 minutes in the dark. Samples were cooled to 23 °C and incubated with 800 µL ethanol for 5 minutes with shaking (1000 rpm). The supernatant was removed, and the beads were then resuspended and washed 3X with 400 µL of 80% (v/v) ethanol. Then, the beads were resuspended in 200 µL of 8 M urea in PBS and 4 µL of reconstituted trypsin/Lys C solution (20 µg lyophilized Trypsin/LysC reconstituted in 40 µL Trypsin/LysC buffer) (Promega; V5073) was added and incubated at 37 °C overnight with shaking at 200 rpm. The next day, the digested samples were transferred to a 15 mL conical tube and incubated with 3.8 mL MeCN at 23 °C for 10 minutes with shaking (1000 rpm). The tubes were placed on a magnetic rack and the beads were allowed to settle. The supernatant was removed, and the beads were resuspended and washed 3X with 1 mL MeCN. Then, the beads were eluted by resuspension with 100 µL 2% (v/v) DMSO in MQ and incubated at 37 °C for 30 minutes with shaking at 1000 rpm. The tubes were placed on a magnetic rack and the supernatants were collected in a microcentrifuge tube. The elution step was repeated, and the supernatants were combined. Peptides were reconstituted in 100 µL LC-MS grade 50% MeCN/H_2_O + 0.1% FA (v/v) via bath sonication for 10 minutes and quantified using Pierce fluorometric assay kit (Thermo; 23290).

### 18O peroxide labeling shotgun proteomics

BPHL protein was diluted to 2 μM in PBS and 1300 µL of protein was treated with 13 μl of 100mM ^18^O-H_2_O_2_ at 37°C for 1 hour with end-over-end rotation. The oxidation was quenched by performing two desalting steps using Zeba 7 kDa columns after equilibrating the columns in PBS. The protein was then split into four 300μl aliquots to which was added either PBS vehicle, hMsrA, mMsrB2, or *E. coli* Msrb all at 4μM, and the samples were incubated at 37°C and 300 rpm overnight in a ThermoMixer. The next day, 5 µL each of magnetic E3 (Cytiva; 44152105050250) and E7 (Cytiva; 24152105050250) SP3 beads were mixed 1:1 and washed 3X with 1 mL MQ H_2_O using a magnetic rack. The washed beads were then resuspended in 20 µL of MQ H_2_O and added to each protein sample. The samples were allowed to incubate at 23 °C for 5 minutes with shaking at 1000 rpm. Protein binding was induced by addition of 800 µL absolute ethanol followed by incubation at 23 °C for 5 minutes with shaking at 1000 rpm. The beads were then placed on a magnetic rack and allowed to settle. The supernatant was removed, and the beads were then resuspended and washed 3X with 400 µL of 80% (v/v) ethanol. The beads were then resuspended in 200 µL of 2 M urea in PBS and incubated with 10 µL of 10 mM TCEP at 65 °C for 15 minutes. Then, 10 µL of 20 mM iodoacetamide was added and incubated at 37 °C for 30 minutes in the dark. Samples were cooled to 23 °C and incubated with 800 µL ethanol for 5 minutes with shaking at 1000 rpm. The supernatant was removed, and the beads were then resuspended and washed 3X with 400 µL of 80% (v/v) ethanol. Then, the beads were resuspended in 200 µL of 1 M urea in PBS and 1 µL of reconstituted trypsin/Lys C solution (20 µg lyophilized Trypsin/LysC reconstituted in 40 µL Trypsin/LysC buffer) (Promega; V5073) was added and incubated at 37 °C overnight with shaking at 200 rpm. The next day, 1 μl of 0.5 mg/mL GluC (Promega; V1651) was added and incubated at 37°C for 3 hours. The digested samples were transferred to a 15 mL conical tube and incubated with 3.8 mL MeCN at 23 °C for 10 minutes with shaking at 1000 rpm. The tubes were placed on a magnetic rack and the beads were allowed to settle. The supernatant was removed, and the beads were washed 3X with 1 mL MeCN. Then, the beads were eluted by resuspension with 100 µL 2% (v/v) DMSO in MQ H_2_O and incubated at 37 °C for 30 minutes with shaking at 1000 rpm. The tubes were placed on a magnetic rack and the supernatants were collected in a microcentrifuge tube. The elution step was repeated, and the supernatants were combined. Peptide concentration was determined by Pierce fluorometric assay kit (Thermo; 23290). Peptides were dried by SpeedVac and reconstituted in 0.1% formic acid at 200 ng/μl for LC-MS/MS analysis.

### ChURRO-ABPP

500 µL of desired proteome was normalized to 4.0 mg/mL and allowed to equilibrate to 23 °C. To 500 µL of lysate was added 5 µL of (*R*)- or (*S*)-ChURRO-2 (200 µM) and incubated at 23 °C for 1 hour. Excess oxaziridine was then quenched by addition of 5 µL *N*-acetyl methionine (final concentration 200 µM) and incubation at 23 °C for 30 minutes. 10 µL of isotopically light isoDTB-N_3_ (1.2 eq) (Vector labs; CCT-1565) tag was added to samples labeled with (*R*)-ChURRO-2 while 10 µL isotopically heavy isoDTB-N_3_ (1.2 eq) (Vector labs; CCT-1565) tag was added to samples labeled with (*S*)-ChURRO-2. 30 µL TBTA (stock in 1:4 DMSO:*^t^*BuOH, 100 µM), 10 µL TCEP (1 mM), and 10 µL CuSO_4_ (1 mM) were added and proteins were unfolded by addition of 50 µL 1.2% SDS in PBS (0.1% w/v). CuAAC was allowed to proceed at 23 °C for 1 hour with rocking. Light and heavy samples were combined in a 1:1 ratio and co-precipitated by addition of 4 volumes of MeOH:CHCl_3_ (4:1.5), vortexed, and centrifuged at 12,000 xg for 15 minutes. The collected protein pellet was washed 1X with 1 mL chilled (-20 °C) MeOH via probe-tip sonication (5 s on, 20 s off, 20% A, 4-6 pulses) on ice and transferred to a 2 mL microcentrifuge tube and centrifuged at 4 °C for 5 minutes at 6,500 xg. The resulting pellet was resuspended in 500 µL 1.2% SDS/PBS via probe-tip sonication (5 s on, 20 s off, 20% A, 6-8 pulses). Samples were then incubated at 90 °C for 5 minutes, cooled to 23 °C, then centrifuged for 5 minutes at 6,500 xg. High-capacity streptavidin agarose beads (200 µL per sample) (Thermo Fisher Scientific; 20357) were washed 3X in 1 mL PBS. The washed beads were then transferred by 2X 250 µL resuspension in PBS to a 15 mL conical tube containing 5 mL PBS (final volume 5.5 mL). The 500 µL resuspended sample was added to the conical tube containing beads and allowed to incubate at 4 °C overnight while rotating.

The following day, samples were allowed to rock at 23 °C for 30 minutes to resolubilize the SDS. The samples were centrifuged at 1,400 xg for 3 minutes and the supernatant was removed. The beads were then washed with 5 mL 0.2% SDS/PBS (w/v), incubated for 10 minutes on a rotator, and centrifuged for 3 minutes at 1,400 xg. The beads were then transferred to micro bio-spin columns (Bio-Rad; 7326204) by 2 x 1 mL resuspension in PBS, placed on a vacuum manifold, and washed thoroughly by 5 x 1 mL PBS washes and 5 x 1 mL MQ washes. The beads were then transferred to low-retention 2.0 mL microcentrifuge tubes by 2 x 250 µL resuspension in 6 M urea/PBS and incubated with 25 µL TCEP (110 mM) at 65 °C for 20 minutes, with gentle agitation every 10 minutes. Then, 25 µL iodoacetamide (400 mM) was added and incubated at 37 °C for 30 minutes in the dark. The beads were washed 2X by addition of 1 mL PBS and centrifuged at 1,400 xg for 3 minutes. The supernatant was removed, and a premixed solution of 200 µL 2 M urea/PBS, 2 µL CaCl_2_ (100 mM), and 8 µL Trypsin/LysC solution (20 µg lyophilized Trypsin/LysC reconstituted in 40 µL Trypsin/LysC buffer) (Promega; V5073) was added and incubated at 37 °C overnight.

The following day, the beads were transferred to micro bio-spin columns (Bio-Rad; 7326204) by 2 x 500 µL resuspension in PBS, placed on a vacuum manifold, and washed thoroughly by 5 x 1 mL PBS washes and 5 x 1 mL MQ washes. The bio-spin columns capped at the bottom, placed in fresh low-retention 2.0 mL microcentrifuge tubes, and peptides were eluted from beads via addition of 200 µL LC-MS grade 50% MeCN/H_2_O + 0.1% FA (v/v) and incubated for 5 minutes. The cap was removed, and samples were centrifuged at 3000 xg for 3 minutes to collect the eluent. The elution step was repeated once more, and elution fractions containing the same sample were combined. The eluent was dried via SpeedVac concentration. Peptides were reconstituted in 100 µL LC-MS grade 50% MeCN/H_2_O + 0.1% FA (v/v) via bath sonication for 10 minutes and quantified using Pierce fluorometric assay kit (Thermo; 23290).

### isoTOP-ABPP for methionine hyper-reactivity

Hyper-reactivity was assessed as previously described^58^. 500 µL of desired proteome was normalized to 4.0 mg/mL and allowed to equilibrate to 23 °C. To 500 µL of lysate was added 5 µL of racemic ChURRO-2 (50 or 500 µM final) and incubated at 23 °C for 1 hour. Excess oxaziridine was then quenched by addition of 5 µL *N*-acetyl methionine (50 µM or 500 µM final) and incubation at 23 °C for 30 minutes. 10 µL of isotopically light isoDTB-N_3_ (1.2 eq) (Vector labs; CCT-1565) tag was added to samples labeled with 500 µM racemic ChURRO-2 while 10 µL isotopically heavy isoDTB-N_3_ (1.2 eq) (Vector labs; CCT-1565) tag was added to samples labeled with 50 µM racemic ChURRO-2. 30 µL TBTA (stock in 1:4 DMSO:*^t^*BuOH, 100 µM), 10 µL TCEP (1 mM), and 10 µL CuSO_4_ (1 mM) were added and proteins were unfolded by addition of 50 µL 1.2% SDS in PBS (0.1% w/v). CuAAC was allowed to proceed at 23 °C for 1 hour with rocking. Light and heavy samples were combined in a 1:1 ratio and co-precipitated by addition of 4 volumes of MeOH:CHCl_3_ (4:1.5), then vortexed and centrifuged at 12,000 xg for 15 minutes. The collected protein pellet was washed 1X with 1 mL chilled (-20 °C) MeOH via probe-tip sonication (5 s on, 20 s off, 20% A, 4-6 pulses) on ice and transferred to a 2 mL microcentrifuge tube and centrifuged at 4 °C for 5 minutes at 6,500 xg. The resulting pellet was resuspended in 500 µL 1.2% SDS/PBS via probe sonication (5 s on, 20 s off, 20% A, 6-8 pulses). Samples were then incubated at 90 °C for 5 minutes, cooled to 23 °C, then centrifuged for 5 minutes at 6,500 xg. High-capacity streptavidin agarose beads (200 µL per sample) (Thermo Fisher Scientific; 20357) were washed 3X in 1 mL PBS. The washed beads were then transferred by 2 x 250 µL resuspension in PBS to a 15 mL conical tube containing 5 mL PBS (final volume 5.5 mL). The 500 µL resuspended sample was added to the conical tube containing beads and allowed to incubate at 4 °C overnight while rotating.

The following day, samples were allowed to rock at 23 °C for 30 minutes to resolubilize the SDS. The samples were centrifuged at 1,400 xg for 3 minutes and the supernatant was removed. The beads were then washed with 5 mL 0.2% SDS/PBS (w/v), incubated for 10 minutes on a rotator, and centrifuged for 3 minutes at 1,400 xg. The beads were then transferred to micro bio-spin columns (Bio-Rad; 7326204) by 2X 1 mL resuspension in PBS, placed on a vacuum manifold, and washed thoroughly by 5 x 1 mL PBS washes and 5 x 1 mL MQ washes. The beads were then transferred to low-retention 2.0 mL microcentrifuge tubes by 2 x 250 µL resuspension in 6 M urea/PBS and incubated with 25 µL TCEP (110 mM) at 65 °C for 20 minutes, with gentle agitation every 10 minutes. Then, 25 µL iodoacetamide (400 mM) was added and incubated at 37 °C for 30 minutes in the dark. The beads were washed 2X by addition of 1 mL PBS and centrifuged at 1,400 xg for 3 minutes. The supernatant was removed, and a premixed solution of 200 µL 2 M urea/PBS, 2 µL CaCl_2_ (100 mM), and 8 µL Trypsin/LysC solution (20 µg lyophilized Trypsin/LysC reconstituted in 40 µL Trypsin/LysC buffer) (Promega; V5073) was added and incubated at 37 °C overnight while rotating.

The following day, the beads were transferred to micro bio-spin columns (Bio-Rad; 7326204) by 2 x 500 µL resuspension in PBS, placed on a vacuum manifold, and washed thoroughly by 5 x 1 mL PBS washes and 5 x 1 mL MQ washes. The bio-spin columns were capped at the bottom, placed in fresh low-retention 2.0 mL microcentrifuge tubes, and peptides were eluted from beads via addition of 200 µL LC-MS grade 50% MeCN/H_2_O + 0.1% FA (v/v) and incubated for 5 minutes. The cap was removed and samples were centrifuged at 3000 xg for 3 minutes to collect the eluent. The elution step was repeated once more, and elution fractions containing the same sample were combined. The eluent was dried via SpeedVac concentration. Peptides were reconstituted in 100 µL LC-MS grade 50% MeCN/H_2_O + 0.1% FA (v/v) via bath sonication for 10 minutes and quantified using Pierce fluorometric assay kit (Thermo; 23290).

### isoTOP-ABPP for *N*-homocysteinylation

500 μL of desired proteome was normalized to 4.0 mg/mL and allowed to equilibrate to 23 °C. Then, the proteomes were diluted to 1 mg/mL using 8 M urea, 27 mM TCEP, pH 6.0 as the diluent for a final concentration of 6 M urea, 20 mM TCEP, pH 6.0 then incubated with 1 mM AT-3 thioester14 for 30 minutes at 23 °C while rotating. Reaction was quenched by adding 80 μL NH_2_OH (50% v/v) and 200 μL iodoacetamide (400 mM) for 1 hour at 23 °C in the dark. Proteins were then precipitated by addition of 4 volumes of MeOH:CHCl_3_ (4:1.5), vortexed, and centrifuged at 12,000 xg for 15 minutes. The collected protein pellet was washed 1X with 1 mL chilled (-20 °C) MeOH via probe- tip sonication (5 s on, 20 s off, 20% A, 4-6 pulses) and resuspended in 500 μL 1.2% SDS/PBS via probe-tip sonication (5 s on, 20 s off, 20% A, 6-8 pulses). 10 μL of isotopically light isoDTB-N_3_ (1.2 eq) (Vector labs; CCT-1565) tag was added to samples treated with no siRNA while 10 μL isotopically heavy isoDTB-N_3_ (1.2 eq) (Vector labs; CCT-1565) tag was added to samples treated with siMsrB2. 30 μL TBTA (stock in 1:4 DMSO:tBuOH, 100 μM), 10 μL TCEP (1 mM), and 10 μL CuSO_4_ (1 mM) were added and proteins were unfolded by addition of 50 μL 1.2% SDS in PBS (0.1% w/v). CuAAC was allowed to proceed at 23 °C for 1 hour with rocking. Light and heavy samples were combined in a 1:1 ratio and co-precipitated by addition of 4 volumes of MeOH:CHCl_3_ (4:1.5), then vortexed and centrifuged at 12,000 xg for 15 minutes. The collected protein pellet was washed 1X with 1 mL chilled (-20 °C) MeOH via probe-tip sonication (5 s on, 20 s off, 20% A, 4-6 pulses) on ice and transferred to a 2 mL microcentrifuge tube and centrifuged at 4 °C for 5 minutes at 6,500 xg. The resulting pellet was resuspended in 500 μL 1.2% SDS/PBS via probe sonication (5 s on, 20 s off, 20% A, 6-8 pulses). Samples were then incubated at 90 °C for 5 minutes, cooled to 23 °C, then centrifuged for 5 minutes at 6,500 xg. High-capacity streptavidin agarose beads (200 μL per sample) (Thermo Fisher Scientific; 20357) were washed 3X in 1 mL PBS. The washed beads were then transferred by 2X 250 μL resuspension in PBS to a 15 mL conical tube containing 5 mL PBS (final volume 5.5 mL). The 500 μL resuspended sample was added to the conical tube containing beads and allowed to incubate at 4 °C overnight while rotating.

The following day, samples were allowed to rock at 23 °C for 30 minutes to resolubilize the SDS. The samples were centrifuged at 1,400 xg for 3 minutes and the supernatant was removed. The beads were then washed with 5 mL 0.2% SDS/PBS (w/v), incubated for 10 minutes on a rotator, and centrifuged for 3 minutes at 1,400 xg. The beads were then transferred to micro bio-spin columns (Bio-Rad; 7326204) by 2 x 1 mL resuspension in PBS, placed on a vacuum manifold, and washed thoroughly by 5 x 1 mL PBS washes and 5 x 1 mL MQ washes. The beads were then transferred to low-retention 2.0 mL microcentrifuge tubes by 2 x 250 μL resuspension in 6 M urea/PBS and incubated with 25 μL TCEP (110 mM) at 65 °C for 20 minutes, with gentle agitation every 10 minutes. Then, 25 μL iodoacetamide (400 mM) was added and incubated at 37 °C for 30 minutes in the dark. The beads were washed 2X by addition of 1 mL PBS and centrifuged at 1,400 xg for 3 minutes. The supernatant was removed, and a premixed solution of 200 μL 2 M urea/PBS, 2 μL CaCl_2_ (100 mM), and 8 μL Trypsin/LysC solution (20 μg lyophilized Trypsin/LysC reconstituted in 40 μL Trypsin/LysC buffer) (Promega; V5073) was added and incubated at 37 °C overnight while rotating.

The following day, the beads were transferred to micro bio-spin columns (Bio-Rad; 7326204) by 2 x 500 μL resuspension in PBS, placed on a vacuum manifold, and washed thoroughly by 5 x 1 mL PBS washes and 5 x 1 mL MQ washes. The bio-spin columns were capped at the bottom, placed in fresh low-retention 2.0 mL microcentrifuge tubes, and peptides were eluted from beads via addition of 200 μL LC-MS grade 50% MeCN/H_2_O + 0.1% FA (v/v) and incubated for 5 minutes. The cap was removed and samples were centrifuged at 3000 xg for 3 minutes to collect the eluent. The elution step was repeated once more, and elution fractions containing the same sample were combined. The eluent was dried via SpeedVac concentration. Peptides were reconstituted in 100 μL LC-MS grade 50% MeCN/H_2_O + 0.1% FA (v/v) via bath sonication for 10 minutes and quantified using Pierce fluorometric assay kit (Thermo; 23290).

### LC-MS/MS data collection

Peptides were analyzed on either a Q Exactive Plus Hybrid Quadrupole-Orbitrap Mass Spectrometer (Thermo; IQLAAEGAAPFALGMBDK) or timsTOF fleX MALDI 2 (Bruker Scientific). Peptides analyzed on the Q Exactive Plus were reconstituted in LC-MS grade H_2_O + 0.1% FA (v/v) via bath sonication for 15 minutes for a final peptide concentration of 50 ng/µL. Approximately 500 ng of peptide was injected and separated via liquid chromatography using a VanquishNeo UHPLC (Thermo; VN-S10-A-01) in Trap-and-Elute configuration using a PepMap Neo (5 µM x 300 µM x 5 mm) trap (Thermo; 174500) and an Aurora Ultimate 25 cm C18 analytical column (IonOpticks; AUR3-25075C18) operating at a 350 nL/min flow rate at a column temperature of 50 °C. Peptides were eluted over a 120 minute gradient from 0-80% H_2_O/MeCN + 0.1% FA and data were collected in positive-ion mode using data-dependent acquisition mode with a default charge state of +2. MS1 was operated at 70,000 mass resolution, 1e6 AGC target, 30 ms maximum injection time, at a scan range of 400 – 1800 m/z. One MS1 scan was followed by 15 MS2 scans of the nth most abundant ions operated at 17,500 mass resolution, 5e4 AGC target, 50 ms maximum injection time, 27 (N)CE, with an isolation window of 1.5 m/z. A 30 s dynamic exclusion window was enabled. Nanospray voltage was set at 2.50 kV and heated capillary temperature at 200 °C.

Peptides analyzed on the timsTOF fleX MALDI 2 were reconstituted in LC-MS grade H_2_O + 0.1% formic acid by bath sonication for 5 mins at a final peptide concentration of 100ng/ml. 100 ng was injected and separated via liquid chromatography using a nanoElute2 UHPLC (Bruker Scientific) in a two column separation method using a PepMap Neo (5 µM x 300 µM x 5 mm) trap column (Thermo; 174500) and a PepSep Max (150μm, 1.5μm) C18 analytical column (Bruker; 1893483) operating at a 500 nL/min flow rate at a column temperature of 50 °C and introduced to the mass spectrometer via a captive spray nano-electrospray ion source operated at 1.6 kV. Peptides were eluted over a 60 minute gradient from 0-90% H_2_O/MeCN + 0.1% FA and data were collected in positive-ion mode using DDA PASEF acquisition. Mass spectra were collected from m/z range of 100-1700 and ion mobility range 1/K0 of 0.70-1.40 Vs/cm^2^, a 100 ms accumulation time, and 4 PASEF MS/MS cycles with an intensity threshold of 2500, charge of +3-+6, and a CID ramp of 20-59 eV.

### Byos analysis

Shotgun data of purified proteins were analyzed with Byologic (Protein Metrics Inc.). Raw files were searched directly against the FASTA sequences of the respective protein pulled from Uniprot database using the Byos HCP workflow, with decoys and common contaminants added. Peptides were assumed fully tryptic. All searches included the following modifications: Acetyl (+42.010565; Protein N-term; variable - rare1), carbamidomethyl (+57.021464; C; Fixed), oxidation (+15.994915; M; variable - common1) and ChURRO-2 labeling (+168.08988; M; variable - common1).

### ChURRO and AT-3 Fragpipe analysis

All samples were analyzed using Fragpipe v21.1, MSFragger v4.1, IonQuant v1.10.27, Philosopher v.5.1.0, and Python v3.9.12. Raw files were directly ported into the Fragpipe GUI under ‘Workflow’. Proteomes were directly downloaded from Uniprot database (human; UP000005640, mouse; UP000000589). The default workflow was used with the following modifications.

MSFragger: Protein digestion was set to enzyme 1: trypsin and enzyme 2: lysc. Variable modifications included oxidation (M, +15.9949, max occurrences 2), *N*-terminal acetylation ([^, +42.0106, max occurrences 1), light ChURRO-2 adduct (M, +649.3660, max occurrences 2), and heavy ChURRO-2 adduct (M, +655.3735, max occurrences 2), light AT-3 adduct (K, +735.3485, max occurrences 2), and heavy AT-3 adduct (M, +741.3561, max occurrences 2). A fixed carbamidomethyl modification (C, +57.021465) was enabled.

Validation: Run validation tools was enabled. Run MSBooster, predict RT, predict spectra was enabled. Run PSM validation, and run Percolator was enabled. Run ProteinProphet was enabled. Generate reports, remove contaminants, generate peptide-level summary, and generate protein-level summary was enabled.

Quant (MS1): Run MS1 quant was enabled. IonQuant Labeling was enabled with light modification of M649.3660 or K735.3485 and heavy modification of M655.3735 or K741.3561 and re-quantify enabled. Normalized intensity across runs was enabled.

After running the search, statistical significance was calculated using a two-tailed t-test of unequal variance with the first array set as the computed log_2_(H/L) ratios across multiple technical replicates (n = 12 for ChURRO studies, n = 4 for *N*-homocysteinylation studies), and the secondary array centered at {0,0}.

### BPHL shotgun proteomics Fragpipe analysis

Samples were analyzed using Fragpipe v23.1, MSFragger v4.3, IonQuant v1.11.11, Philosopher v.5.1.0, and Python v3.11.11. Raw files were directly ported into the Fragpipe GUI under ‘Workflow’. The BPHL fasta sequence was uploaded in the “Database” tab with decoys and contaminants added. The default workflow was used with the following modifications.

MSFragger: Protein digestion was set to enzyme 1: trypsin_gluc. Variable modifications included oxidation (M, +15.9949, max occurrences 1), *N*-terminal acetylation ([^, +42.0106, max occurrences 1, light ChURRO-2 adduct (for ChURRO-2 BPHL labeling) (M, +168.08988, max occurrences 1), and ^18^O-H_2_O_2_ adduct (for Msr rescue studies) (M, +17.99916, max occurrences 1). A fixed carbamidomethyl modification (C, +57.021465) was enabled.

Validation: Run validation tools was enabled. Run MSBooster, predict RT, predict spectra was enabled. Run PSM validation, and run Percolator was enabled. Run ProteinProphet was enabled. Generate reports, remove contaminants, generate peptide-level summary, and generate protein-level summary was enabled.

Quant (MS1): Run MS1 quant was enabled. IonQuant LFQ was enabled with Add MaxLFQ. Match between runs and Normalize intensity across runs were selected and MBR ion FDR set to 0.01.

After running the search, the LFQ intensities for all occurrences of each Met site were summed, and the summed LFQ intensity of peptides containing the indicated modification were then normalized by dividing by the total summed LFQ intensity of each Met site.

### Ramachandran analysis

For each amino acid residue, the φ (phi) and ψ (psi) dihedral angles were extracted from their protein’s predicted AlphaFold structure. The AlphaFold structures were queried from the Alpha Fold Protein Structure Database^61^ using the python package StructureMap (https://github.com/MannLabs/structuremap), adapted from Bludau et al.^62^ The BioPython package^63^ was then used to parse the queried CIFs and extract the psi and phi angles of the amino acids of interest. The dihedral angles of the modified methionine residues were then plotted and intensities were calculated using kernel density estimation to construct Ramachandran plots. The source code for replicating this analysis, along with detailed instructions, are available at https://github.com/ritster/ChURRO_ABPP.

### Crowdedness analysis

Amino acid crowdedness (pPSE) predictions were done using StructureMap (https://github.com/MannLabs/structuremap), adapted from Bludau et al.^62^ and White et al.^64^ In short, for each amino acid residue, a cone region is projected from the β-carbon (or the α-carbon for glycine) at length (12 Å). Within each cone, the number of α-carbon atoms from neighboring amino acids is counted, considering the tolerance of the predicted alignment error as reported by AlphaFold. This count is referred to as the pPSE (proximal PSE) value, which reflects the crowdedness/accessibility of the local environment around each amino acid. The source code for replicating this analysis, along with detailed instructions, are available at https://github.com/ritster/ChURRO_ABPP.

### Subcellular localization

An in-house python script was used to search for subcellular localization using subcellular localization data pulled from the Uniprot database and employing regex patterning with user- defined identifiers (e.g. Nucleus = ‘Nucle[a-z]*’, Cytoplasm = ‘Cytoplasm[a-z]*’). Percent localization was determined by dividing the number of proteins localized to a specific organelle over the total number of proteins.

### PANTHER GO

Biological activity was determined via “PANTHER GO biological process complete” analysis^65–67^ using Uniprot IDs of all statistically significant protein hits and plotted based if a biological process was deemed statistically significant and > 25% enriched.

### AlphaMissense

The public AlphaMissense database was used to determine predicted mutation outcomes from mitochondrial datasets. Briefly, mouse protein Met sites were aligned to their human isoforms and the human Met site information was used to map mutation outcome. https://console.cloud.google.com/storage/browser/dm_alphamissense;tab=objects?prefix=&forceOnObjectsSortingFiltering=false&pli=1

### DFT

DFT calculations were conducted at the Molecular Graphics and Computational Facility (MGCF) at the University of California, Berkeley, using the Gaussian 16 software package. All calculations, including geometry optimizations, were performed in H_2_O solvent using the SMD solvent continuum model reported by Truhlar and co-workers (scrf=smd)^68^. Initial geometries were constructed in GaussView16 and subjected to conformational searches using Maestro (Schrodinger). Conformers within a 5 kcal / mol window were subsequently optimized to stationary points (either minima or first-order saddle points) using the M06-2X functional and the basis set 6-31(d,p). The nature of each stationary point was evaluated by accompanying frequency calculations (all positive eigenvalues for minima and exactly one negative eigenvalue for transition states). Several isomers were considered, but only those representing the minimum-energy paths are presented here. Transition states were connected to ground states by following the intrinsic reaction coordinate (IRC) downhill from the transition state. All other parameters were left as their defaults unless otherwise stated.

### RoseTTAFold-All-Atom structure modeling

RoseTTAFold-All-Atom^48^ was downloaded and installed following instructions provided at https://github.com/baker-laboratory/RoseTTAFold-All-Atom. BPHL *(R)*-MetO 69 was modeled using the covalent config provided in the downloaded package with the following modifications. The FASTA file used was that of the human BPHL protein sequence (Uniprot Q86WA6) with amino acid 69 changed from M to G. The small molecule input was a .sdf file of 1-[*(R)*-methylsulfinyl]ethane. A covalent bond was designated between the alpha carbon of G69 and carbon 2 of 1-[*(R)*-methylsulfinyl]ethane.

### Statistical information

Replicate information is reported in the figure legends and relevant methods sections. P-values were calculated via student’s t-test using a two-tailed, two-sample test of unequal variance (P-values * < 0.05, ** < 0.01, *** < 0.001). Bar graphs with error bars represent mean + SD.

## Acknowledgments

We thank Dr. A. Killilea and her staff at the UC Berkeley Cell Culture Facility for cell culture support (RRID: SCR_017924). We thank Dr. A. Jain and Prof. R. Zoncu for providing cells for mito-IP comparison and suggestions for mitochondrial purification. We thank Dr. T. Xiao for providing mouse liver tissue. We thank Dr. H. Celik for assistance with NMR spectroscopy (NIH S10OD024998). We thank Drs. V. Roytman, K. Durkin, and D. Small for computational support (NIH S10OD034382). We thank Drs. Rodney Levine and Geumsoo Kim for providing purified MsrA and B proteins. We thank Drs. John Eng and Venu Vandavasi at the Princeton Mass Spectrometry and Small Instrument Core Facility for technical support.

## Author contributions

A.G.-V., A.C.S.P., A.G.R., F.D.T., and C.J.C. conceived of this study. A.C.S.P., A.G.-V., J.M.B., F.A., K.S., and T.G.C. synthesized the chemical probes used in this study. S.C.V. performed the SFC chiral separation. J.N. and R.D. supported VCD measurements. A.C.S.P. performed DFT computations. A.G.-V. and C.J.C. designed the proteomic experiments and A.G.-V., A.C.S.P., and J.M.B. performed the proteomic experiments. A.G.-V. and A.C.S.P. purified mitochondria. A.G.-V. performed biochemical experiments on purified proteins. J.K. generated CRISPR lines and performed cellular assays. A.L.G. and A.S. raised and fed mice and harvested tissue. R.R.S., A.A.A., X.X., and N.D. performed bioinformatic analyses for this study based on discussions with A.G.-V., A.C.S.P., J.M.B., and D.H. The initial manuscript was written by A.G.-V. and A.C.S.P. All authors contributed to the editing of this manuscript.

## Data and materials availability

All materials and methods, including proteomic datasets, are included in the Supplementary Materials. Raw MS datafiles are available on the MassIVE repository (https://massive.ucsd.edu/) under identifier MSV000099597. Code used in this study for pPSE and Ramachandran analysis are located on Github (https://github.com/ritster/ChURRO_ABPP). AlphaMissense Pathogenicity Prediction was performed by downloading from the public AlphaMissense database available at (https://console.cloud.google.com/storage/browser/dm_alphamissense;tab=objects?prefix=&forceOnObjectsSortingFiltering=false).

## Supplementary Materials

Supplemental Materials and Methods

Tables S1-3

NMR Spectra

---

**Supplementary Fig. 1:**
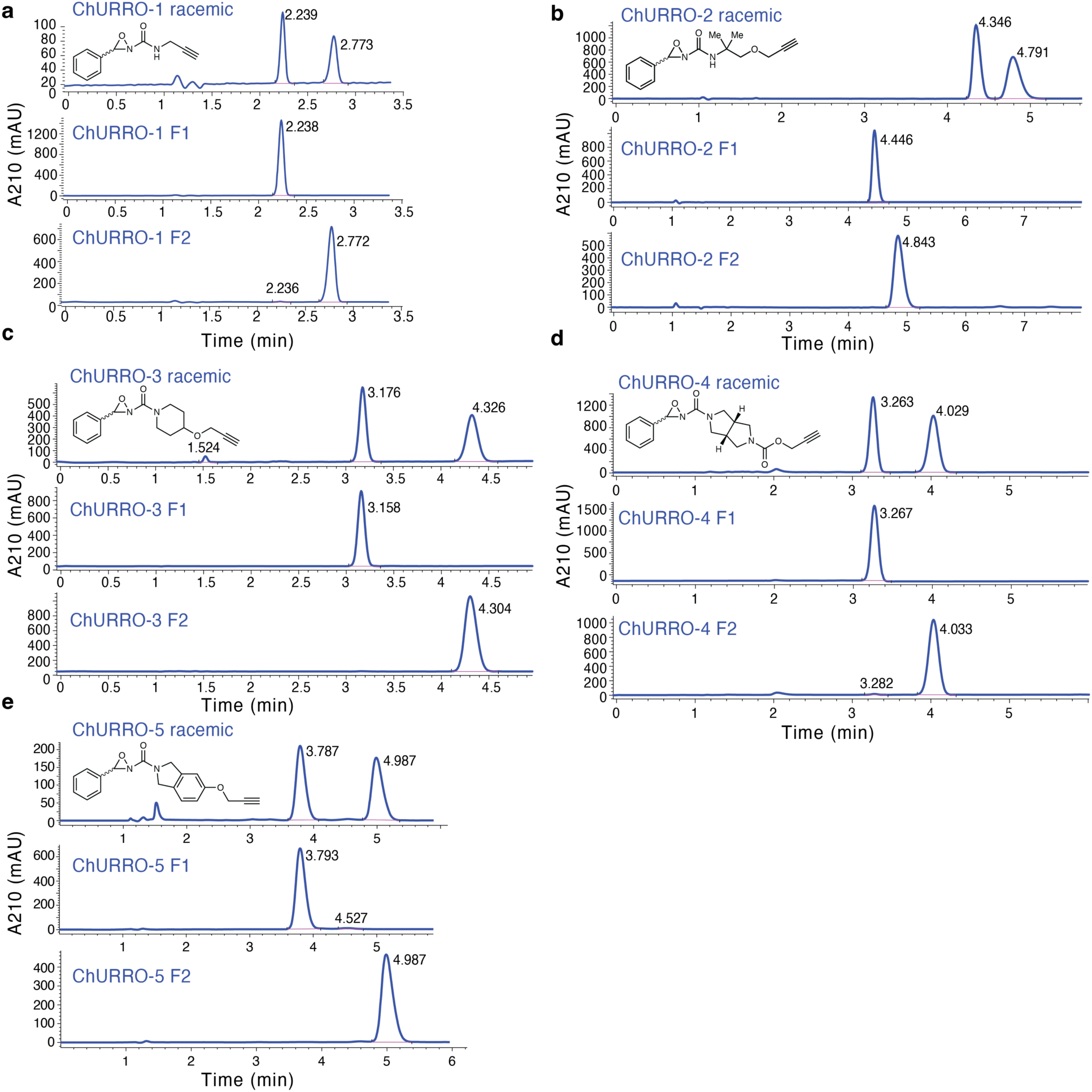
Chiral SFC separation of ChURRO probe enantiomers. **a**, ChURRO-1 preparative SFC. (*R*)-ChURRO-1 is the first eluting enantiomer and enantiopurity was determined by analytical chiral SFC retention time: 2.2 min (99%). (*S*)-ChURRO-1 is the second eluting enantiomer and enantiopurity was determined by analytical chiral SFC retention time: 2.8 min (99%). **b**, ChURRO-2 preparative SFC. (*R*)-ChURRO-2 is the first eluting enantiomer and enantiopurity was determined by analytical chiral SFC retention time: 4.4 min (99%). (*S*)-ChURRO-2 is the second eluting enantiomer and enantiopurity was determined by analytical chiral SFC retention time: 4.8 min (99%). **c**, ChURRO-3 preparative SFC. (*R*)-ChURRO-3 is the first eluting enantiomer and enantiopurity was determined by analytical chiral SFC retention time: 3.2 min (99%). (*S*)-ChURRO-3 is the second eluting enantiomer and enantiopurity was determined by analytical chiral SFC retention time: 4.3 min (99%). **d**, ChURRO-4 preparative SFC. (*R*)-ChURRO-4 is the first eluting enantiomer and enantiopurity was determined by analytical chiral SFC retention time: 3.2 min (99%). (*S*)-ChURRO-4 is the second eluting enantiomer and enantiopurity was determined by analytical chiral SFC retention time: 4.3 min (99%). **e**, ChURRO-5 preparative SFC. (*R*)-ChURRO-5 is the first eluting enantiomer and enantiopurity was determined by analytical chiral SFC retention time: 3.8 min (99%). (*S*)-ChURRO-5 is the second eluting enantiomer and enantiopurity was determined by analytical Chiral SFC retention time: 5.0 min (99%).

**Supplementary Fig. 2:**
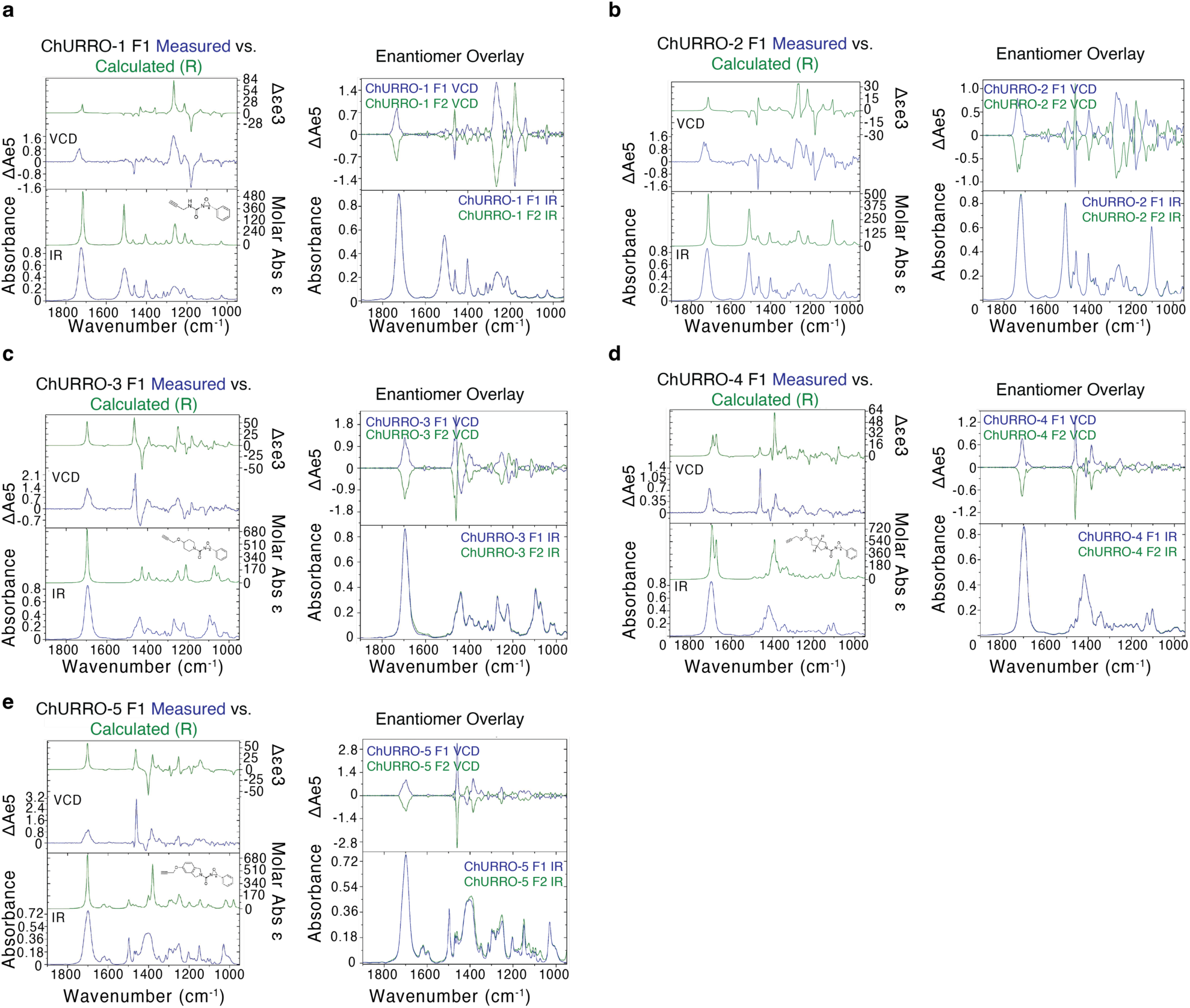
VCD and IR analysis of ChURRO enantiomers. **a-e**, (Left) measured and calculated IR and VCD spectra of first eluting enantiomer of ChURRO probe, assigned as (*R*)-ChURRO. (Right) overlay of measured VCD and IR spectra of both ChURRO enantiomers assigned as (*R*) and (*S*), respectively, in order of elution on the analytical SFC column.

**Supplementary Fig. 3:**
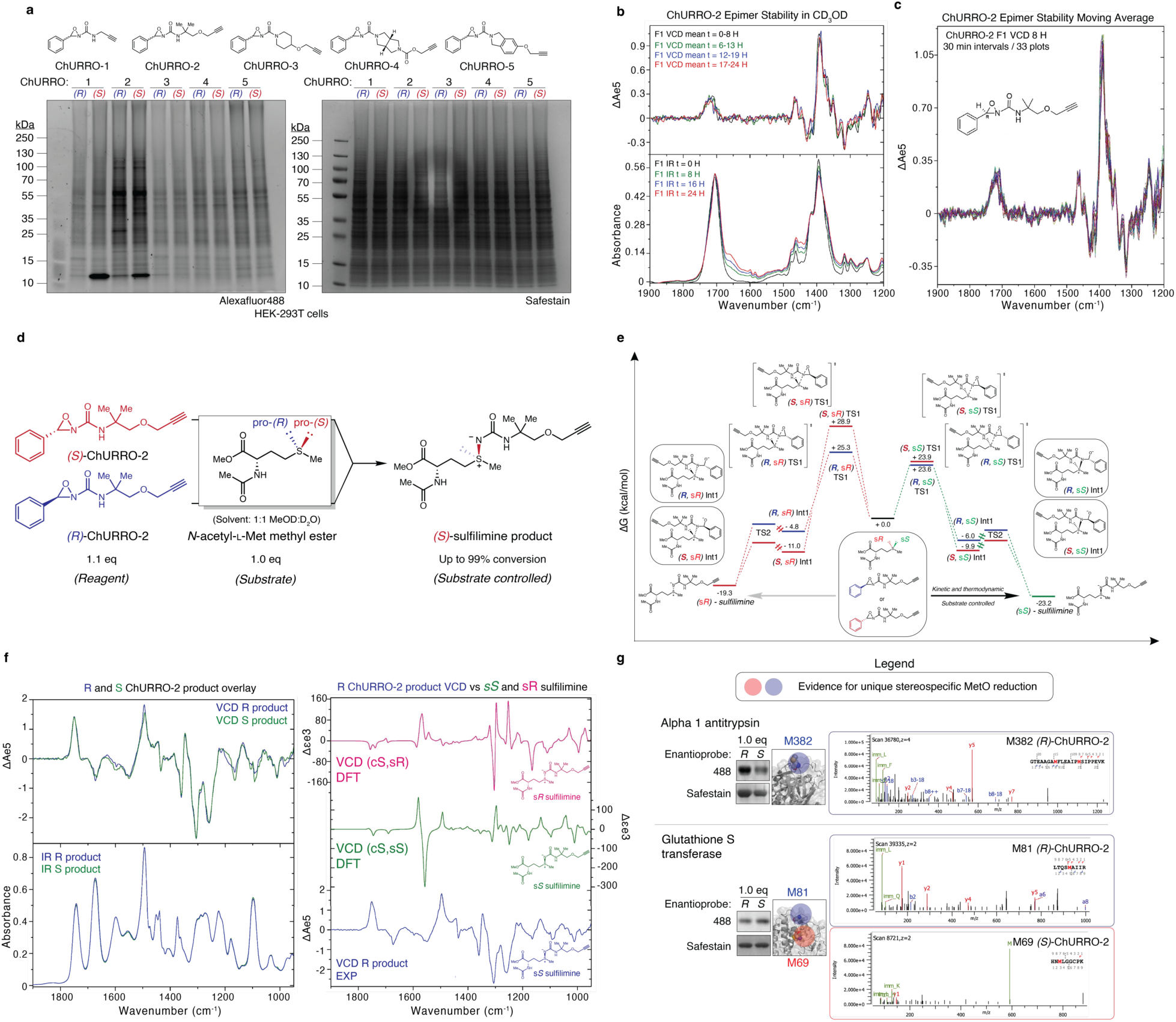
Assessing ChURRO reactivity on monomers, proteins, and proteomes. **a**, In-gel fluorescence labeling comparison of all 5 ChURRO probes with equimolar doses (100 µM) in HEK-293T lysates (4 mg/mL). **b**, (Top) 8 hour averaged VCD spectra of ChURRO-2 F1 in CD_3_OD taken over 24 hours at 37 °C. (Bottom) 8 hour averaged IR spectra of ChURRO-2 F1 in CD_3_OD taken over 24 hours. **c**, Half-difference baseline corrected VCD spectra representative of ChURRO-2 F1 and F2 (referred to in text as (*R*) and (*S*)) for the 24 hour time course. **d**, Model diastereoselectivity studies of (*R*)- or (*S*)-ChURRO-2 (50 mM) incubated with *N*-acetyl-L-methionine methyl ester (55 mM) in cosolvent (1:1 CD_3_OD:D_2_O) at room temperature for 30 minutes. Product was confirmed by ^1^H NMR and VCD measurement. **e**, Reaction coordinate diagram of (*R*)- or (*S*)-ChURRO-2 reacting with a model methionine substrate. **f**, (Left) overlay of measured VCD and IR spectra of products isolated from reaction of (*R*)-ChURRO-2 and (*S*)-ChURRO-2 with *N*-acetyl methionine methyl ester. (Right) measured VCD spectrum of isolated sulfilimine product from (*R*)-ChURRO-2 reaction vs calculated VCD spectra of (s*R*) and (s*S*) sulfilimine diastereomers. **g**, ChURRO labeling of previously characterized proteins that have evidence for stereospecific MetO reduction assessed by in-gel fluorescence and shotgun proteomics.

**Supplementary Fig. 4:**
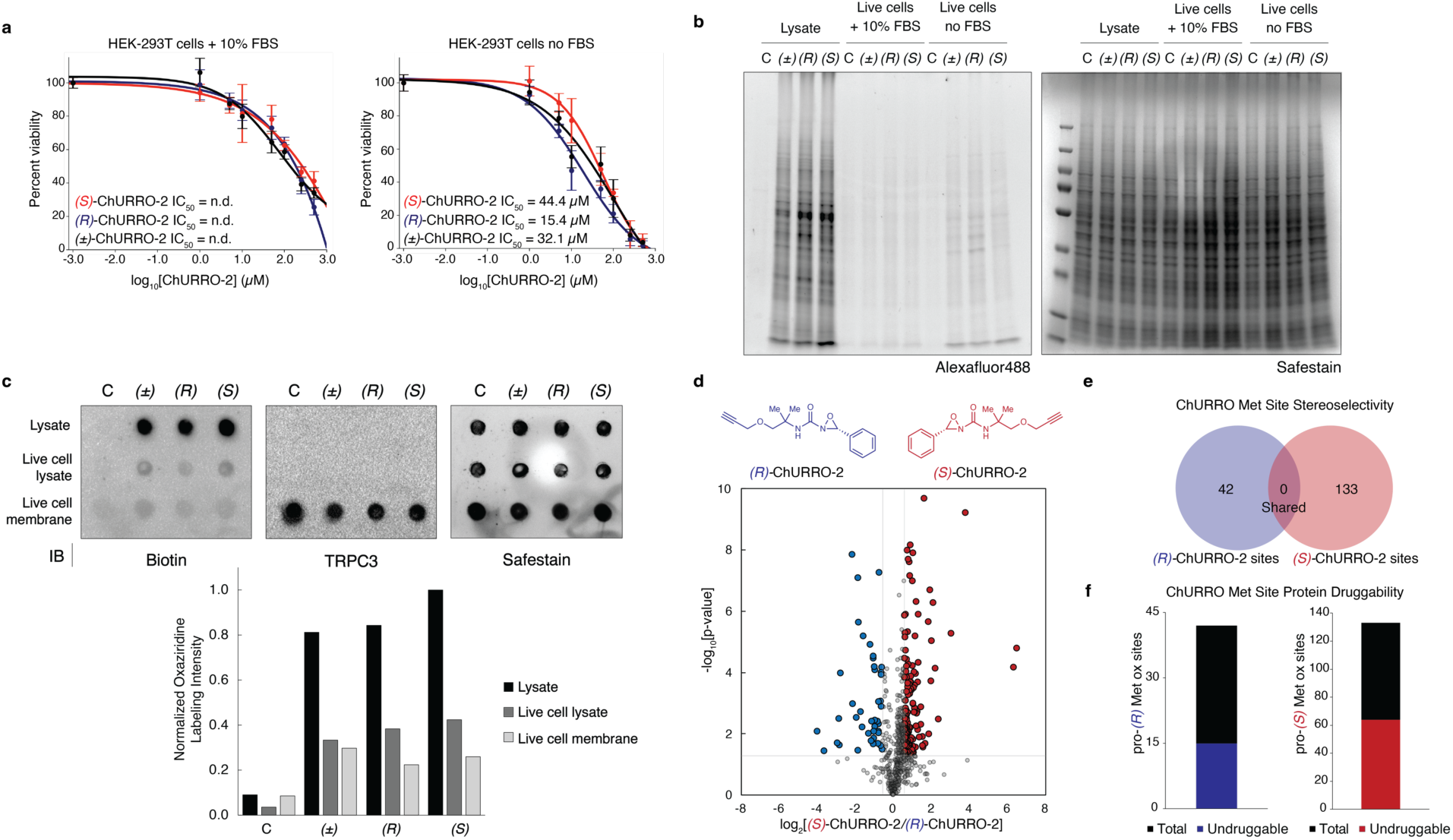
Live cell and lysate labeling experiments of ChURRO-2. **a**, Dose response curves of cell viability with or without FBS in the media (n = 3 bioreplicates, error bars represent standard deviation). **b**, In-gel fluorescence labeling comparison of equimolar doses of ChURRO-2 enantiomers (50 µM). **c**, Dot blot assessment to determine probe localization to the plasma membrane by TRPC3 staining (membrane marker). **d**, Volcano plots of quantitative proteomic experiments conducted in HEK-293T lysates with equimolar 200 µM ChURRO-2 enantiomer doses in 4 mg/mL lysate (n = 3 biological replicates, n = 3 technical MS replicates). **e**, proteomic identification of ChURRO-2 methionine sites showcases high stereoselectivity for (*R*)- and (*S*)-Met oxidation sites. **f**, ChURRO-2 identifies therapeutically relevant Met oxidation sites with high stereoselectivity in HEK-293T lysates.

**Supplementary Fig. 5:**
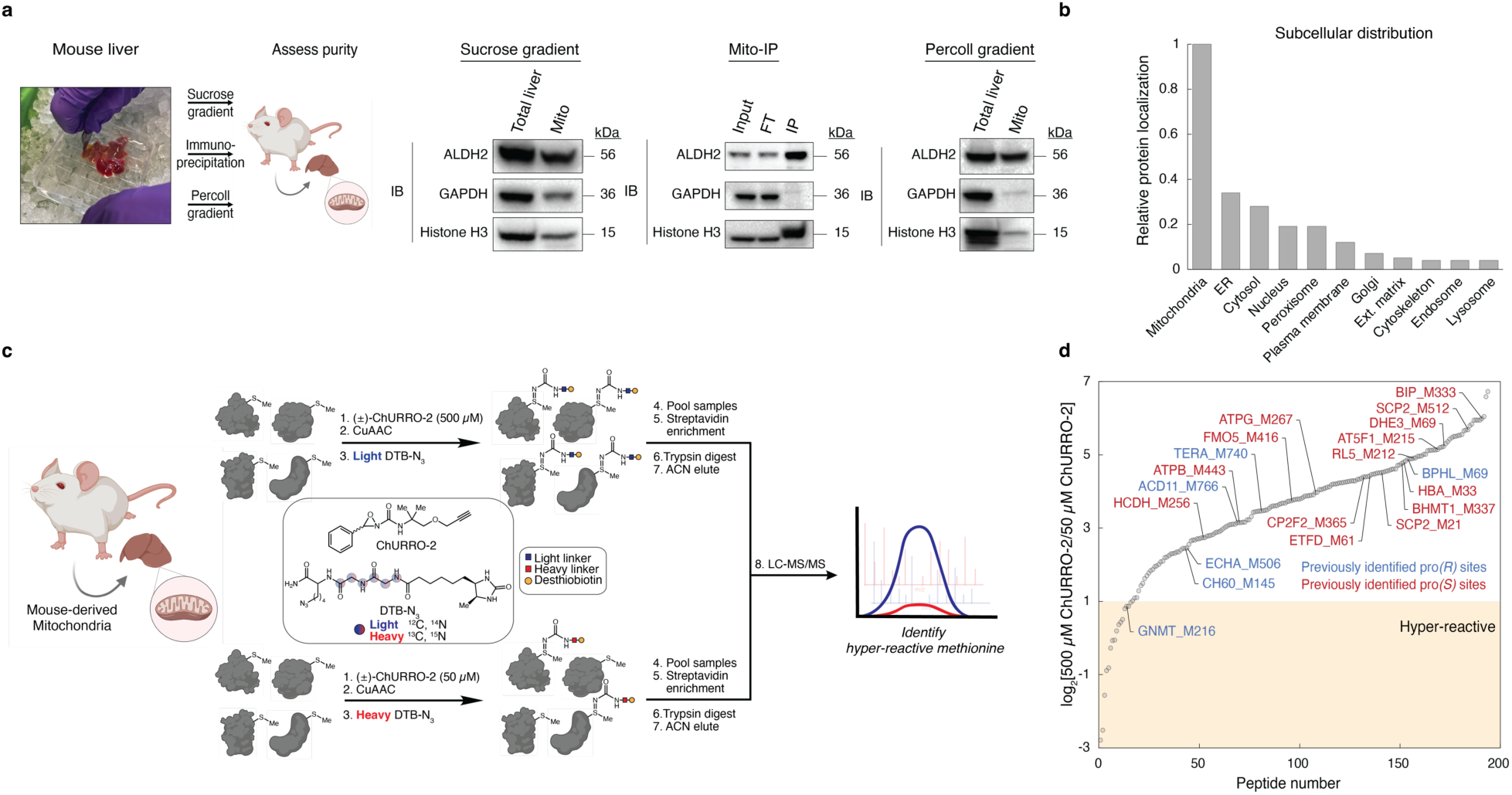
Subcellular ChURRO profiling of the mitochondria. **a**, Comparison of mitochondrial enrichment methods assessed by immunoblotting of ALDH2 (mitochondrial), GAPDH (cytosolic), or Histone H3 (nuclear). Sucrose gradient protocol adapted from Dias et. al^69^ and Mito-IP protocol adapted from Chen et. al.^49^ **b**, Normalized subcellular distribution of proteins identified from mitochondrial prochiral Met oxidation site dataset. **c**, isoTOP-ABPP workflow to identify hyper-reactive methionine sites in the mitochondria. **d**, Reactivity profiles of Met sites in mitochondria with identified prochiral Met oxidation sites labeled (n = 3 technical MS replicates).

**Supplementary Fig. 6:**
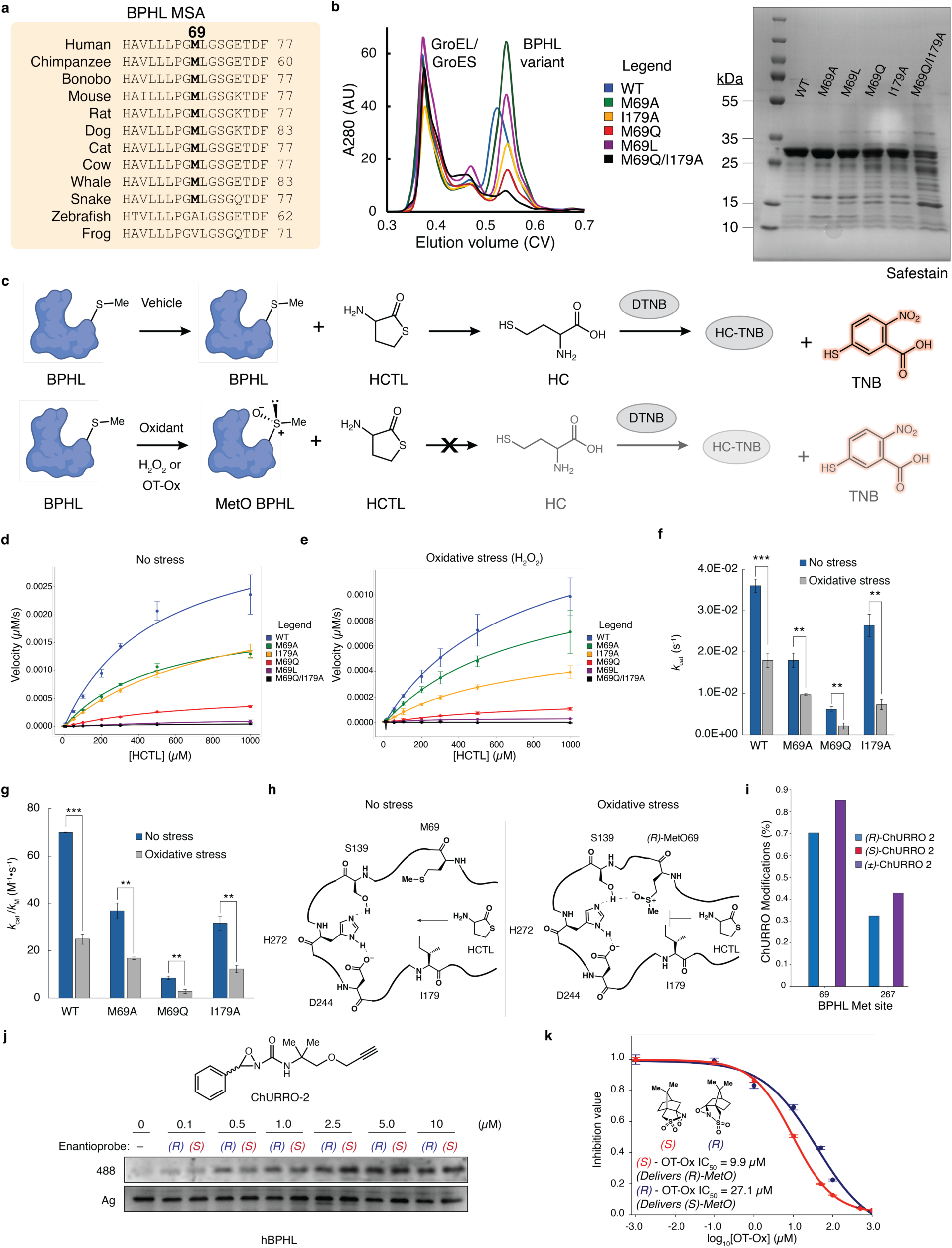
BPHL contains a regulatory M69 site that is stereospecifically oxidized and reduced. **a**, Clustal omega multiple sequence alignment of BPHL across multiple organisms. **b**, (Left) SEC chromatograms of all BPHL variants generated with aid of protein chaperones GroEL/ES. (Right) SDS-PAGE gel analysis of purified BPHL variants. **c**, Schematic of *in vitro* BPHL activity assay using native HCTL substrate and Ellman’s reagent (DTNB) to oxidize HC and form TNB as a colorimetric readout. **d-e**, HCTLase activity curves of all 6 BPHL variants under conditions of no stress (0 µM H_2_O_2_) and oxidative stress (500 µM H_2_O_2_) with determined Michaelis-Menten parameters (n = 3 technical replicates, error bars represent standard deviation). **f-g**, Additional Michaelis-Menten parameters of 4 BPHL variants under conditions of no stress and oxidative stress (n = 3 technical replicates, error bars represent standard deviation). **h**, Proposed mechanism of pro-*(R)*-MetO inhibition at 69 site. **i**, ChURRO-2 shotgun proteomic peptide spectrum matches for different BPHL methionine sites. **j**, Gel-ABPP of ChURRO-2 enantiomers on purified human BPHL. **k**, Dose-response curves of BPHL treated with camphor-derived Davis oxaziridines for asymmetric MetO formation (n = 3 technical replicates, error bars represent standard deviation).

**Supplementary Fig. 7:**
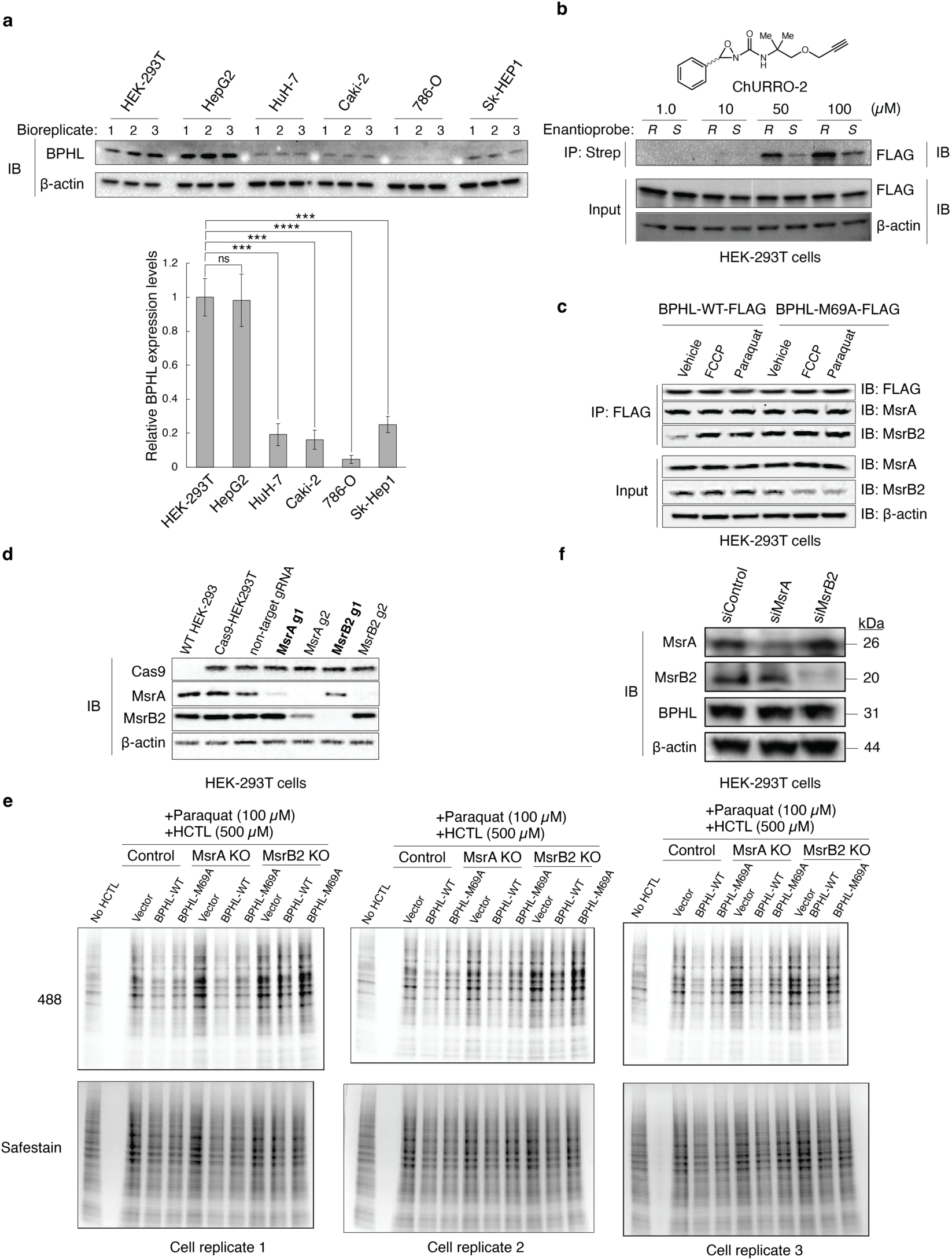
Assessing the impact of BPHL prochiral methionine oxidation on cellular *N*-homocysteinylation status. **a**, BPHL expression screen in renal and hepatic cell lines assessed by immunoblotting (n = 3 technical cell replicates, error bars represent standard deviation). **b**, Immunoprecipitation studies of FLAG-BPHL expressed in HEK-293T cells using (*R*)- and (*S*)-ChURRO-2. **c**, Representative immunoblot of BPHL Co-IP with MsrA and MsrB2 (n = 3 cell technical replicates). **d**, Immunoblot analysis of MsrA and MsrB2 KO efficiency. **e**, In-gel fluorescence data of labeled *N*-homocysteinylated proteins in HEK-293T cells with MsrA or MsrB2 KO (n = 3 technical cell replicates). **f**, Assessment of Msr knock-down efficiency in HEK-293T cells.

