## Supplementary Information for "Chiral methionine oxidation reagents reveal stereospecific proteome modifications"

**The file includes:**

- Materials and Methods
- Tables S1-3
- NMR Spectra

#### Materials and Methods

**General methods:** Reactions using moisture- or air-sensitive reagents were carried out in flame-dried glassware under an inert atmosphere of $N_2$. Solvent was passed over activated alumina and stored under argon before use when dry solvent was required. All other commercially purchased chemicals were used as received. Compound **AT-3** was prepared according to Chen et al.[49] without modification. Merck 60 F254 silica gel precoated sheets (0.25 mm thick) were used for analytical thin layer chromatography and visualized by fluorescence quenching under UV light or by staining with $KMnO_4$ or Ninhydrin. Flash chromatography was performed using a CombiFlash NextGen 100 or Biotage Selekt automated chromatography system with pre-packed silica gel columns. $^{1}$H and $^{13}$C NMR spectra were collected at 298 K in $CDCl_3$ or $CD_3OD$ (Cambridge Isotope Laboratories, Cambridge, MA) using Bruker AVQ-400, AVB-400, AV-500, AV-600, NEO-500, or NEO-501 instruments at the Pines Magnetic Resonance Center's Core NMR Facility at the University of California, Berkeley. All chemical shifts are reported in the standard notation of $\delta$ parts per million relative to the residual solvent peak at 7.26 ($CDCl_3$) or 3.31 ($CD_3OD$) for $^{1}$H and 77.16 ($CDCl_3$) or 49.00 ($CD_3OD$) for $^{13}$C as an internal reference. Splitting patterns are indicated as follows: br, broad; s, singlet; d, doublet; t, triplet; m, multiplet; dd, doublet of doublets. Enantioenrichment was determined by chiral HPLC using a Daicel CHIRALPAK® ADH column (4.6 x 250 mm) located in the Toste group at the University of California, Berkeley or by chiral SFC using a Chiral Technologies AD-H or OJ-H column (4.6 x 250 mm) located at the California Institute of Technology. Absolute configuration of oxaziridine enantiomers and sulfilimine products was determined by VCD. Experimental spectra were collected on a Chiral*IR-2X™* Dual*PEM* FT-VCD spectrometer (BioTools Inc, Jupiter, FL). Theoretical VCD spectra were calculated using Gaussian09 and compared using the CompareVOA™ program (BioTools Inc). Low-resolution electrospray mass spectral analyses were carried out using LC-MS (Agilent Technology 6130, Quadrupole LC/MS and Advion Expression-L Compact Mass Spectrometer). High-resolution mass spectral analyses (ESI-MS) were carried out using an Agilent 6230 LC-TOF (ESI) located in the Toste group at the University of California, Berkeley. All aqueous solutions for biological experiments were prepared using Milli-Q water, and all *in vitro* experiments were carried out in PBS, pH 7.4, unless otherwise noted. All biological experiments were prepared using freshly prepared aliquots.

#### Chemistry Methods

**Synthesis of ChURRO-1:**

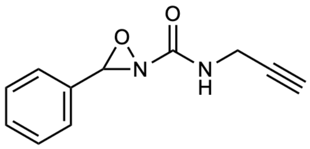


Compound **ChURRO-1** and its precursors were synthesized as a racemic mixture of enantiomers using previously reported procedures[28]. The NMR spectra of **ChURRO-1** were in accordance with previously published spectra.

**Synthesis of ChURRO-2:**

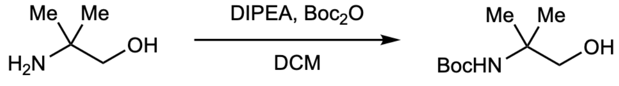


SI-1

***tert*-Butyl (1-hydroxy-2-methylpropan-2-yl)carbamate (SI-1)**

To a solution of 2-amino-2-methyl propanol (9.54 mL, 100 mmol, 1.11 equiv.) in DCM (20 mL) was added DIPEA (15.68 mL, 90 mmol, 1.00 equiv.) and Boc anhydride (19.64 g, 90 mmol, 1.00 equiv.) in DCM (23.75 mL). The reaction was allowed to stir at room temperature overnight. After 16 hours, the reaction was quenched by addition of 10% aqueous citric acid, diluted with DCM, and washed successively with water and brine. The organic layers were dried over $MgSO_4$, filtered, and concentrated under reduced pressure to give **SI-1** as a white powder (14.50 g, 76.6 mmol, 85% yield). $^1H$ NMR (600 MHz, $CDCl_3$) δ 4.65 (s, 1H), 4.00 (s, 1H), 3.65 – 3.50 (m, 2H), 1.43 (s, 9H), 1.25 (s, 6H). $^{13}C$ NMR (151 MHz, $CDCl_3$) δ 156.30, 79.94, 71.05, 54.48, 28.51, 24.88. Compound did not ionize on ESI.

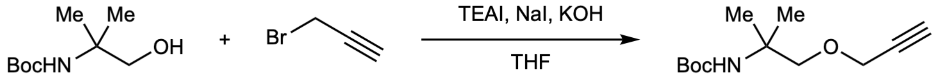


SI-2

***tert*-Butyl (2-methyl-1-(prop-2-yn-1-yloxy)propan-2-yl)carbamate (SI-2)**

A flame-dried round bottomed flask equipped with a magnetic stir bar was charged with THF (20 mL), alcohol **SI-1** (1.50 g, 7.93 mmol, 1.00 equiv.), propargyl bromide (80 wt% in toluene, 1.06 mL, 9.52 mmol, 1.20 equiv.), sodium iodide (0.204 g, 1.36 mmol, 0.17 equiv.), TEAI (0.204 g, 0.79 mmol, 0.10 equiv.), and KOH (0.89 g, 15.86 mmol, 2.00 equiv.). The resulting orange suspension was stirred vigorously overnight and quenched by addition of 20 mL $H_2O$ and washed with EtOAc (3 x 50 mL). The combined organic layers were dried over $Na_2SO_4$, filtered, and concentrated under reduced pressure. The crude yellow oil was purified using flash column chromatography (15% EtOAc/Hexanes) to give **SI-2** as a yellow oil (1.06 g, 4.67 mmol, 59% yield). $^1H$ NMR (600 MHz, $CDCl_3$) δ 4.70 (s, 1H), 4.16 (d, $J$ = 2.4 Hz, 2H), 3.47 (s, 2H), 2.42 (t, $J$ = 2.4 Hz, 1H), 1.43 (s, 9H), 1.30 (s, 6H). $^{13}C$ NMR (151 MHz, $CDCl_3$) δ 154.95, 79.81, 79.01, 76.25, 74.61, 58.67, 52.66, 28.58, 24.38. Compound did not ionize on ESI.

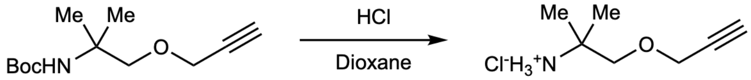


SI-3

**2-Methyl-1-(prop-2-yn-1-yloxy)propan-2-aminium chloride (SI-3)**

To a flame-dried round bottomed flask equipped with a magnetic stir bar was added a solution of carbamate **SI-2** (2.85 g, 12.52 mmol, 1.00 equiv.) and anhydrous HCl (27.54 mmol, 2.20 equiv., 4M in 1,4-dioxane) in anhydrous MeOH (25 mL). The reaction was allowed to stir overnight, and the resulting pink solution was concentrated under reduced pressure. The crude pink solid was washed with cold $Et_2O$ to afford the HCl salt of the amine as a white crystalline solid (1.50 g, 9.17 mmol, 73% yield) that was used immediately in the next step without further purification.

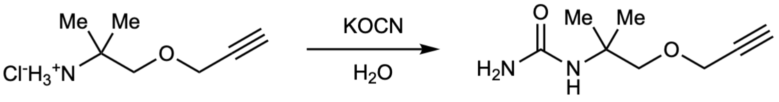


SI-4

**1-(2-Methyl-1-(prop-2-yn-1-yloxy)propan-2-yl)urea (SI-4)**

To a solution of amine **SI-3** (1.50 g, 9.17 mmol, 1.00 equiv.) in $H_2O$ (9.17 mL) was added potassium cyanate (2.98 g, 36.68 mmol, 4.00 equiv.). The reaction was heated to 60 °C and allowed to stir overnight. After 16 hours, the reaction was cooled to room temperature and extracted with EtOAc (5 x 100 mL). The combined organic layers were dried over $MgSO_4$, filtered, and concentrated under reduced pressure to afford the urea **SI-4** as a white powder (1.05 g, 6.19 mmol, 68% yield). $^1$H NMR (500 MHz, $CDCl_3$) δ 4.86 (s, 1H), 4.47 (s, 2H), 4.17 (d, $J$ = 2.4 Hz, 2H), 3.51 (s, 2H), 2.44 (t, $J$ = 2.4 Hz, 1H), 1.34 (s, 6H). $^{13}$C NMR (126 MHz, $CDCl_3$) δ 158.41, 79.64, 76.96, 74.80, 58.69, 52.99, 24.73. HRMS (ESI) calculated for $[C_8H_{14}N_2O_2+H]^+$: required *m/z* 171.1128, found *m/z* 171.1124.

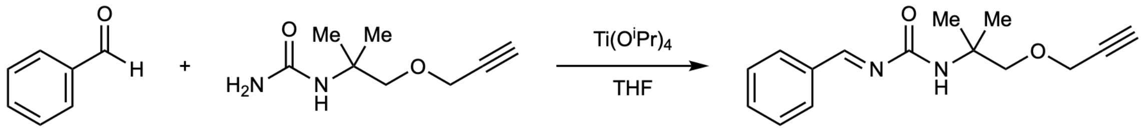


SI-5

**N-Benzylidene-3-(2-methyl-1-(prop-2-yn-1-yloxy)propan-2-yl)urea (SI-5)**

A flame-dried round bottomed flask equipped with a magnetic stir bar was charged with urea **SI-4** (1.05 g, 6.19 mmol, 1.00 equiv.) and evacuated and backfilled with $N_2$ twice. The urea was dissolved in THF (18.76 mL), to which benzaldehyde (0.76 mL, 7.43 mmol, 1.20 equiv.) and $Ti(O^iPr)_4$ (2.57 mL, 8.67 mmol, 1.40 equiv.) were added. The resulting mixture was stirred vigorously overnight. After 16 hours, the reaction was concentrated under reduced pressure and used immediately in the next step without further purification.

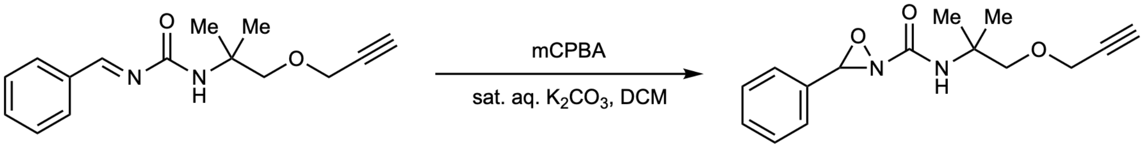


ChURRO_2

***N*-(2-Methyl-1-(prop-2-yn-1-yloxy)propan-2-yl)-3-phenyl-1,2-oxaziridine-2-carboxamide (ChURRO-2)**

To a solution of DCM (20 mL) and sat. aq. $K_2CO_3$ (20 mL) was added mCPBA (3.16 g, 18.57 mmol, 3.00 equiv.). The resulting slurry was stirred vigorously at room temperature for 10 minutes at which point a suspension of crude imine **SI-5** in DCM (15 mL) was added. The resulting yellow solution was allowed to stir at room temperature. After 8 hours, the reaction was stopped by addition of $H_2O$ (100 mL) and extracted with DCM (3 x 50 mL). The combined organic layers were dried over $MgSO_4$, filtered, and concentrated under reduced pressure. The resulting oil was purified using flash column chromatography (DCM/MeOH) to afford the racemic **ChURRO-2** as a clear oil. (0.226 g, 0.83 mmol, 13% yield over 2 steps). $^{1}$H NMR (600 MHz, $CDCl_3$) δ 7.50 – 7.38 (m, 5H), 6.20 (s, 1H), 4.98 (s, 1H), 4.24 – 4.16 (m, 2H), 3.59 (d, $J$ = 9.0 Hz, 1H), 3.50 (d, $J$ = 9.0 Hz, 1H), 2.47 (t, $J$ = 2.4 Hz, 1H), 1.41 (s, 3H), 1.39 (s, 3H). $^{13}$C NMR (151 MHz, $CDCl_3$) δ 161.12, 132.81, 131.02, 128.73, 128.12, 79.56, 79.33, 75.43, 75.02, 58.68, 53.99, 23.98, 23.73. HRMS (ESI) calculated for $[C_{15}H_{18}N_2O_3+H]^+$: required *m/z* 275.1390, found *m/z* 275.1383.

**Synthesis of ChURRO-3:**

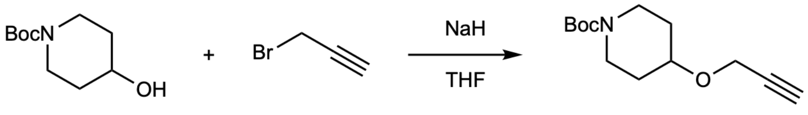


**SI-6**

***tert*-butyl 4-(prop-2-yn-1-yloxy)piperidine-1-carboxylate (SI-6)**

A flame-dried round bottomed equipped with a magnetic stir bar was charged with *tert*-butyl 4-hydroxypiperidine-1-carboxylate (5.00 g, 24.8 mmol, 1.00 equiv.) and evacuated and backfilled with $N_2$, after which anhydrous THF (80 mL) was added via syringe. NaH (60% suspension in mineral oil, 1.99 g, 49.70 mmol, 2.00 equiv.) was added in portions under an $N_2$ blanket over the course of a few minutes (bubbling observed). The cloudy grey reaction mixture was allowed to stir at ambient temperature for 45 minutes, after which propargyl bromide (80% wt in toluene, 1.39 mL, 67.10 mmol, 2.70 equiv.) was added dropwise via syringe. The brown reaction was stirred at ambient temperature for 3 hours and then cooled in an ice bath. Water (50 mL) was slowly added (vigorous bubbling) to quench the reaction and it was extracted with EtOAc (200 mL). The aqueous layer was washed with EtOAc (3 x 50 mL). The combined organic layers were dried over $MgSO_4$, filtered, and concentrated under reduced pressure. The resulting residue was purified via flash chromatography (0% to 20% EtOAc/Hexanes) to afford **SI-6** as a yellow oil that solidified upon storage at –20 °C (4.01 g, 16.62 mmol, 67% yield). $^{1}$H NMR (600 MHz, $CDCl_3$) δ 4.14 (d, J = 2.4 Hz, 2H), 3.78 – 3.61 (m, 3H), 3.12 – 2.97 (m, 2H), 2.38 (t, J = 2.4 Hz, 1H), 1.84 – 1.73 (m, 2H), 1.57 – 1.44 (m, 2H), 1.40 (s, 9H). $^{13}$C NMR (151 MHz, $CDCl_3$) δ 154.80, 80.10, 79.48, 74.16, 73.69, 55.16, 41.38, 40.91, 30.79, 28.45. HRMS (ESI) calculated for $[C_{13}H_{21}NO_3+Na]^+$: required *m/z* 262.14136, found *m/z* 262.14140.

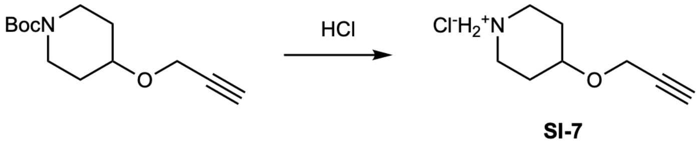


**SI-7**

**4-(prop-2-yn-1-yloxy)piperidin-1-ium chloride (SI-7)**

To a flame-dried round bottomed flask equipped with a magnetic stir bar was added a solution of carbamate **SI-6** (2.00 g, 8.36 mmol, 1.00 equiv.) and HCl (20.90 mL, 83.60 mmol, 10 equiv., 4M in 1,4-dioxane). The reaction mixture was allowed to stir for 2 hours and then concentrated under reduced pressure . The crude solid was washed with cold $Et_2O$ to afford **SI-7** as an off-white solid (1.25 g, 7.12 mmol, 85% yield) that was used immediately in the next step without further purification.

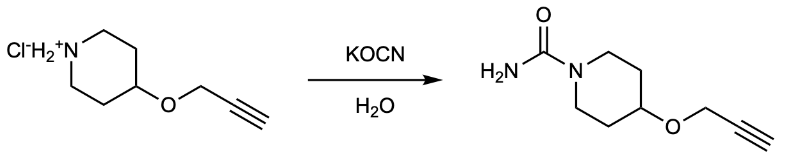


**SI-8**

**4-(prop-2-yn-1-yloxy)piperidine-1-carboxamide (SI-8)**

To a solution of amine **SI-7** (1.00 g, 5.70 mmol, 1.00 equiv.) in $H_2O$ (5.70 mL) was added potassium cyanate (0.92 g, 11.40 mmol, 2.00 equiv.). The reaction was heated to 60 °C and allowed to stir for 6 hours, after which the solvent was removed under reduced pressure. The residue was washed with 50 mL of 85/15 $CHCl_3$/IPA, and the suspension was filtered, dried over $Na_2SO_4$, and concentrated under reduced pressure to afford **SI-8** as a white solid (0.88g, 4.85 mmol, 85% yield). $^1H$ NMR (600 MHz, $CD_3OD$) δ 4.22 (d, J = 2.4 Hz, 2H), 3.81 – 3.74 (m, 1H), 3.73 – 3.65 (m, 2H), 3.23 – 3.09 (m, 2H), 2.82 (t, J = 2.4 Hz, 1H), 1.98 – 1.74 (m, 2H), 1.68 – 1.38 (m, 2H). $^{13}C$ NMR (151 MHz, $CD_3OD$) δ 161.04, 81.09, 75.42, 74.67, 56.04, 42.41, 31.81. HRMS (ESI) calculated for $[C_9H_{14}N_2O_2+H]^+$: required m/z 183.11280, found m/z 183.11271.

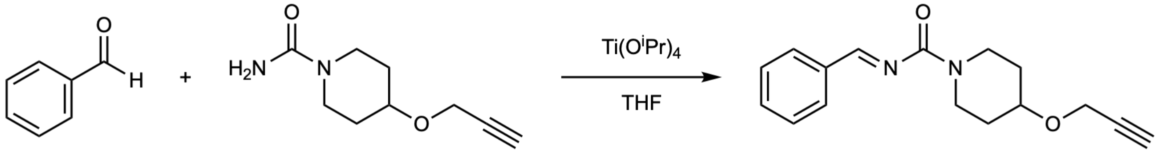


**SI-9**

**N-benzylidene-4-(prop-2-yn-1-yloxy)piperidine-1-carboxamide (SI-9)**

A flame-dried round bottomed flask equipped with a magnetic stir bar was charged with urea **SI-8** (0.324 g, 1.78 mmol, 1.00 equiv.) and evacuated and backfilled with $N_2$ twice. The urea was dissolved in THF (1 mL), to which benzaldehyde (0.22 mL, 2.14 mmol, 1.20 equiv.) and $Ti(O^iPr)_4$ (0.76 mL, 2.49 mmol, 1.40 equiv.) were added. The resulting mixture was stirred vigorously overnight. After 16 hours, the reaction was concentrated under reduced pressure and used immediately in the next step without further purification.

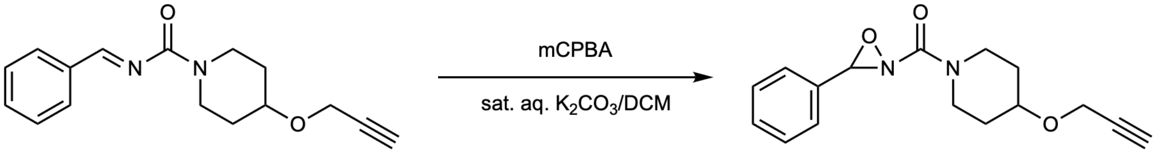


**ChURRO_3**

**(3-phenyl-1,2-oxaziridin-2-yl)(4-(prop-2-yn-1-yloxy)piperidin-1-yl)methanone (ChURRO-3)**

To a solution of DCM (10 mL) and sat. aq. $K_2CO_3$ (7.5 mL) was added mCPBA (0.92 g, 5.34 mmol, 3.00 equiv.). The resulting slurry was stirred vigorously at room temperature for 10 minutes at which point a suspension of crude imine **SI-9** in DCM (5 mL) was added. The resulting yellow solution was allowed to stir at room temperature. After 2 hours, the reaction was stopped by addition of $H_2O$ (25 mL) and extracted with DCM (3 x 50 mL). The combined organic layers were dried over $Na_2SO_4$, filtered, and concentrated under reduced pressure. The resulting oil was purified via flash chromatography (50:50:0 to 45:45:10 DCM/Hexanes/$Et_2O$) to afford racemic **ChURRO-3** as a clear oil. (0.217 g, 0.80 mmol, 45% yield over 2 steps). $^1H$ NMR (600 MHz, $CDCl_3$) δ 7.57 – 7.36 (m, 5H), 5.22 (s, 0.5H), 5.21 (s, 0.5H), 4.20 (d, J = 2.4 Hz, 1H), 4.19 (d, J = 2.4 Hz, 1H), 4.12 – 4.00 (m, 0.5H), 3.96 – 3.77 (m, 2H), 3.76 – 3.64 (m, 1H), 3.56 – 3.48 (m, 1H), 3.37 (m, 0.5H), 2.42 (t, J = 2.4 Hz, 1H), 2.04 – 1.83 (m, 2H), 1.79 – 1.61 (m, 2H). $^{13}C$ NMR (151 MHz, $CDCl_3$) δ 160.74, 160.73, 133.06, 133.02, 130.85, 128.77, 128.02, 79.97, 78.08, 78.03, 74.45, 72.82, 72.47, 55.57, 55.45, 42.20, 41.86, 41.24, 40.87, 31.11, 30.98, 30.43, 30.22. HRMS (ESI) calculated for $[C_{16}H_{18}N_2O_3+H]^+$: required m/z 287.13902, found m/z 287.13911.

**Synthesis of ChURRO-4:**

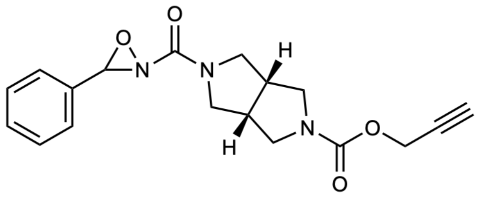


Compound **ChURRO-4** and its precursors were synthesized as a racemic mixture of enantiomers using previously reported procedures[34]. The NMR spectra of **ChURRO-4** were in accordance with previously published spectra.

**Synthesis of ChURRO-5:**

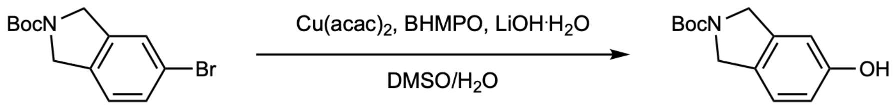


**SI-10**

***tert*-butyl 5-hydroxyisoindoline-2-carboxylate (SI-10)**

A flame-dried round bottomed flask equipped with a magnetic stir bar was charged with *tert*-butyl 5-bromoisoindoline-2-carboxylate (5.00 g, 16.80 mmol, 1.00 equiv.), copper(II) acetylacetonate (0.088 g, 0.34 mmol, 0.02 equiv.), bis(4-hydroxy-2,6-dimethylphenyl)oxalamide (0.11 g, 0.34 mmol, 0.02 equiv.), and lithium hydroxide monohydrate (1.55 g, 36.90 mmol, 2.20 equiv.). The contents of the flask were evacuated and backfilled with $N_2$ three times, after which degassed DMSO (13 mL) and degassed $H_2O$ (3.25 mL) were added via syringe. The reaction was heated to 80 ℃ and allowed to stir for 24 hours. The reaction was then allowed to cool down to ambient temperature and acidified with 2M HCl (pH <5) and extracted with EtOAc (3 x 50 mL). The combined organic layers were washed with brine, dried over $Na_2SO_4$, filtered, and concentrated under reduced pressure. The resulting residue was purified using flash chromatography (10% to 60% EtOAc/Hexanes) to afford **SI-10** as an off-white solid (2.40 g, 10.25 mmol, 61% yield). Note: compound exists as a 1:1 mixture of rotamers in $CDCl_3$ at 298 K. $^1H$ NMR (600 MHz, $CDCl_3$) δ 7.08 (d, J = 8.2 Hz, 0.5H), 7.05 (d, J = 8.2 Hz, 0.5H), 6.84 – 6.71 (m, 2H), 6.41 (s, 0.5H), 6.36 (s, 0.5H), 4.62 (d, J = 14.6 Hz, 2H), 4.58 (d, J = 14.6 Hz, 2H), 1.52 (s, 9H). $^{13}C$ NMR (151 MHz, $CDCl_3$) δ 156.05, 155.99, 155.03, 154.98,

138.82, 138.41, 128.81, 128.52, 123.75, 123.50, 115.06, 115.04, 109.79, 109.53, 80.22, 52.54, 52.20, 51.98, 51.62, 28.71. HRMS (ESI) calculated for $[C_{13}H_{17}NO_3+Na]^+$: required m/z 258.11006, found m/z 258.10990.

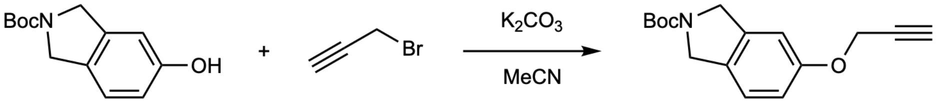


**SI-11**

**_tert_-butyl 5-(prop-2-yn-1-yloxy)isoindoline-2-carboxylate (SI-11)**

A flame-dried round bottomed flask equipped with a magnetic stir bar was charged with **SI-10** (1.50 g, 6.40 mmol, 1.00 equiv.) and potassium carbonate (1.80 g, 13.00 mmol, 2.00 equiv.). The contents of the flask were evacuated and backfilled with $N_2$ three times, after which anhydrous MeCN (32 mL) and propargyl bromide (80% wt in toluene, 0.76 mL, 7.00 mmol, 1.10 equiv.) were added via syringe. The reaction was heated to 40 °C and allowed to stir overnight. The reaction was then allowed to cool down to ambient temperature and the solids were filtered, washing with EtOAc (2 x 10 mL). The solvent was removed under reduced pressure and the resulting residue was purified via flash chromatography (0% to 30% EtOAc/Hexanes) to afford **SI-11** as an off-white solid (1.34 g, 4.93 mmol, 77% yield). Note: compound exists as a 1:1 mixture of rotamers in $CDCl_3$ at 298 K. $^1H$ NMR (600 MHz, $CDCl_3$) δ 7.18 (d, J = 8.2 Hz, 0.5H), 7.13 (d, J = 8.2 Hz, 0.5H), 6.93 – 6.77 (m, 2H), 4.68 (apparent t, J = 2.2 Hz, 2H), 4.66 – 4.57 (m, 4H), 2.51 (t, J = 2.5 Hz, 1H), 1.51 (s, 9H). $^{13}C$ NMR (151 MHz, $CDCl_3$) δ 157.43, 157.39, 154.63, 154.60, 138.94, 138.56, 130.46, 130.15, 123.68, 123.45, 114.92, 114.67, 109.18, 79.77, 78.59, 78.57, 75.74, 56.20, 52.55, 52.26, 51.89, 51.57, 28.67. HRMS (ESI) calculated for $[C_{16}H_{19}NO_3+Na]^+$: required m/z 296.12571, found m/z 296.12568.

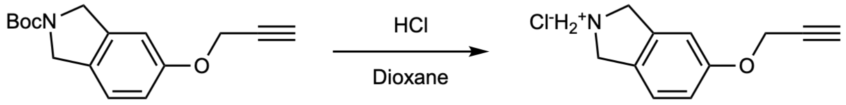


**SI-12**

**5-(prop-2-yn-1-yloxy)isoindolin-2-ium chloride (SI-12)**

To a flame-dried vial equipped with a magnetic stir bar was added a solution of carbamate **SI-11** (0.95 g, 3.50 mmol, 1.00 equiv.) and HCl (8.70 mL, 35.00 mmol, 10.00 equiv., 4M in 1,4-dioxane). The orange reaction mixture was allowed to stir for 2 hours, after which a pinkish-white precipitate crashed out. The crude solid was washed with $Et_2O$ to afford **SI-12** as a white solid (0.73 g, 3.50 mmol, quantitative yield) that was used immediately in the next step without further purification.

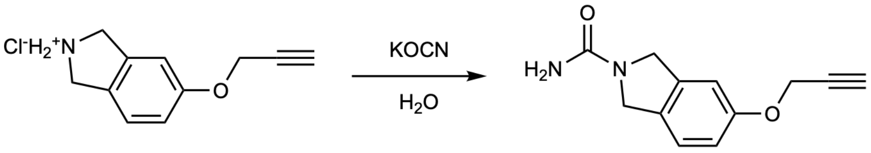


**SI-13**

**5-(prop-2-yn-1-yloxy)isoindoline-2-carboxamide (SI-13)**

To a solution of amine **SI-12** (0.73 g, 3.50 mmol, 1.00 equiv.) in $H_2O$ (3.5 mL) was added potassium cyanate (0.85 g, 10.00 mmol, 3.00 equiv.). The reaction was heated to 60 °C and allowed to stir overnight. After 16 hours, the reaction was cooled to room temperature and extracted with EtOAc (5 x 50 mL). The combined organic layers were dried over MgSO4, filtered, and concentrated under reduced pressure to afford the urea **SI-13** as an off-white powder (0.545 g, 2.52 mmol, 72% yield). $^1$H NMR (600 MHz, $CD_3OD$) δ 7.20 (d, J = 8.2 Hz, 1H), 6.97 – 6.88 (m, 2H), 4.71 (d, J = 2.4 Hz, 2H), 4.63 (s, 2H), 4.59 (s, 2H), 2.93 (t, J = 2.4 Hz, 1H). $^{13}$C NMR (151 MHz, $CD_3OD$) δ 160.51, 159.12, 139.36, 130.69, 124.52, 116.15, 110.00, 79.74, 76.79, 56.89, 53.22, 52.56. HRMS (ESI) calculated for $[C_{12}H_{12}N_2O_2+H]^+$: required m/z 217.09715, found m/z 217.09785.

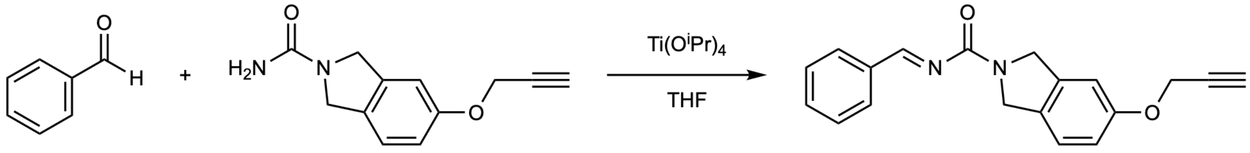


**SI-14**

**N-benzylidene-5-(prop-2-yn-1-yloxy)isoindoline-2-carboxamide (SI-14)**

A flame-dried round bottomed flask equipped with a magnetic stir bar was charged with urea **SI-13** (0.231 g, 1.07 mmol, 1.00 equiv.) and evacuated and backfilled with $N_2$ twice. The urea was dissolved in THF (11 mL), to which benzaldehyde (0.13 mL, 1.28 mmol, 1.20 equiv.) and Ti(O$^i$Pr)$_4$ (0.45 mL, 1.50 mmol, 1.40 equiv.) were added. The resulting mixture was stirred vigorously overnight. After 16 hours, the reaction was concentrated under reduced pressure and used immediately in the next step without further purification.

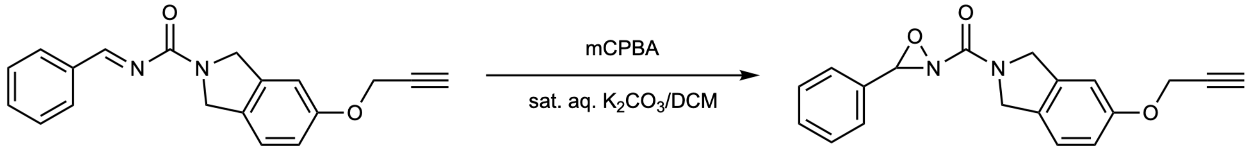


**ChURRO_5**

**(3-phenyl-1,2-oxaziridin-2-yl)(5-(prop-2-yn-1-yloxy)isoindolin-2-yl)methanone (ChURRO-5)**

To a solution of DCM (7.5 mL) and sat. aq. $K_2CO_3$ (5 mL) was added mCPBA (0.553 g, 3.20 mmol, 3.00 equiv.). The resulting slurry was stirred vigorously at room temperature for 10 minutes at which point a suspension of crude imine **SI-14** in DCM (2.5 mL) was added. The resulting yellow solution was allowed to stir at room temperature. After 2 hours, the reaction was stopped by addition of $H_2O$ (50 mL) and extracted with DCM (3 x 50 mL). The combined organic layers were dried over $Na_2SO4$, filtered, and concentrated under reduced pressure. The resulting residue was purified using flash column chromatography (50:50:0 to 45:45:10 DCM/Hexanes/$Et_2O$) to afford the racemic **ChURRO-5** as an off-white solid. (0.133 g, 0.42 mmol, 39% yield over 2 steps). Note: compound exists as a 1:1 mixture of rotamers in $CDCl_3$ at 298 K. $^1$H NMR (600 MHz, $CDCl_3$) δ 7.57 – 7.52 (m, 2H), 7.50 – 7.39 (m, 3H), 7.22 (d, J = 8.4 Hz, 0.5H), 7.16 (d, J = 8.4 Hz, 0.5H), 7.00 – 6.84 (m, 2H), 5.30 (s, 0.5H), 5.29 (s, 0.5H), 5.13 (d, J = 14.5 Hz, 0.5H), 5.08 (d, J = 14.5 Hz, 0.5H), 4.90 – 4.77 (m, 3H), 4.70 (d, J = 2.4 Hz, 1H), 4.69 (d,

J = 2.4 Hz, 1H), 2.53 (t, J = 2.4 Hz, 0.5H), 2.52 (t, J = 2.4 Hz, 0.5H). $^{13}C$ NMR (151 MHz, $CDCl_3$) δ 160.57, 160.52, 157.80, 157.75, 137.80, 137.05, 133.07, 130.97, 129.24, 128.83, 128.53, 128.10, 123.83, 123.68, 115.41, 115.37, 109.28, 109.20, 78.42, 78.04, 78.03, 75.91, 56.26, 56.23, 53.41, 52.80, 52.72, 52.12. HRMS (ESI) calculated for $[C_{19}H_{16}N_2O_3+H]^+$: required m/z 321.12337, found m/z 321.12337.

**Preparative SFC separation:** Preparative oxaziridine separation was conducted at the California Institute of Technology by Dr. Scott Virgil via chiral supercritical fluid chromatography (SFC). The enantiomers were purified on a JASCO System 2000 preparative SFC system with a Chiral Technologies 21.2 x 250 mm AD-H column, a Chiral Technologies 20 x 250 mm OZ-H column, or a Chiral Technologies 10 x 250 mm OJ-H column. The fractions were analyzed on a Thar SFC system with a 4.6 x 250 mm Chiral Technologies AD-3, AD-H or OJ-H column.

**ChURRO-1:** Enantiomers were purified via preparatory SFC (OJ-H column, 30% $IPA/CO_2$, 18 mL/min, 210 nm).
**(*R*)-ChURRO-1** is the first eluting enantiomer on OJ-H column. Enantiopurity was determined by analytical chiral SFC (OJ-H column, 30% $IPA/CO_2$, 2.5 mL/min, 210 nm) retention time: 2.2 min (99%)
**(*S*)-ChURRO-1** is the second eluting enantiomer on OJ-H column. Enantiopurity was determined by analytical chiral SFC (OJ-H column, 30% $IPA/CO_2$, 2.5 mL/min, 210 nm) retention time: 2.8 min (99%)

**ChURRO-2:** Enantiomers were purified via preparatory SFC (OZ-H column, 25% $IPA/CO_2$, 25 mL/min, 210 nm).
**(*R*)-ChURRO-2** is the first eluting enantiomer on AD-H column and the second eluting enantiomer on OZ-H column. Enantiopurity was determined by analytical chiral SFC (AD-H column, 10% $IPA/CO_2$, 2.5 mL/min, 210 nm) retention time: 4.4 min (99%). $^1H$ NMR (500 MHz, $CDCl_3$) δ 7.50 – 7.37 (m, 5H), 6.20 (s, 1H), 4.98 (s, 1H), 4.20 (t, *J* = 2.3 Hz, 2H), 3.59 (d, *J* = 9.0 Hz, 1H), 3.49 (d, *J* = 9.0 Hz, 1H), 2.47 (t, *J* = 2.4 Hz, 1H), 1.40 (d, *J* = 5.8 Hz, 6H). $^{13}C$ NMR (126 MHz, $CDCl_3$) δ 161.12, 132.78, 131.03, 128.73, 128.12, 79.55, 79.32, 75.41, 75.03, 58.67, 53.98, 23.96, 23.72. HRMS (ESI) calculated for $[C_{15}H_{18}N_2O_3+Na]^+$: required *m/z* 297.1210, found *m/z* 297.1211.
**(*S*)-ChURRO-2** is the second eluting enantiomer on AD-H column and the first eluting enantiomer on OZ-H column. Enantiopurity was determined by analytical chiral SFC (AD-H column, 10% $IPA/CO_2$, 2.5 mL/min, 210 nm) retention time: 4.8 min (99%). $^1H$ NMR (500 MHz, $CDCl_3$) δ 7.50 – 7.37 (m, 5H), 6.20 (s, 1H), 4.98 (s, 1H), 4.20 (t, *J* = 2.3 Hz, 2H), 3.59 (d, *J* = 9.1 Hz, 1H), 3.49 (d, *J* = 9.1 Hz, 1H), 2.47 (t, *J* = 2.4 Hz, 1H), 1.40 (d, *J* = 5.8 Hz, 6H). $^{13}C$ NMR (126 MHz, $CDCl_3$) δ 161.11, 132.78, 131.02, 128.73, 128.11, 79.55, 79.32, 75.40, 75.03, 58.66, 53.98, 23.96, 23.71. HRMS (ESI) calculated for $[C_{15}H_{18}N_2O_3+Na]^+$: required *m/z* 297.1210, found *m/z* 297.1210.

**ChURRO-3:** Enantiomers were purified via preparatory SFC (AD-H column, 35% $EtOH/CO_2$, 35 mL/min, 210 nm).

**(*R*)-ChURRO-3** is the first eluting enantiomer on AD-H column. Enantiopurity was determined by analytical chiral SFC (AD-H column, 35% EtOH/$CO_2$, 2.5 mL/min, 210 nm) retention time: 3.2 min (99%).
**(*S*)-ChURRO-3** is the second eluting enantiomer on AD-H column. Enantiopurity was determined by analytical chiral SFC (AD-H column, 35% EtOH/$CO_2$, 2.5 mL/min, 210 nm) retention time: 4.3 min (99%).

**ChURRO-4:** Enantiomers were purified via preparatory SFC (OJ-H column, 35% EtOH/$CO_2$, 18 mL/min, 210 nm).
**(*R*)-ChURRO-4** is the first eluting enantiomer on OJ-H column. Enantiopurity was determined by analytical chiral SFC (AD-H column, 35% EtOH/$CO_2$, 2.5 mL/min, 210 nm) retention time: 3.3 min (99%).
**(*S*)-ChURRO-4** is the second eluting enantiomer on OJ-H column. Enantiopurity was determined by analytical chiral SFC (AD-H column, 35% EtOH/$CO_2$, 2.5 mL/min, 210 nm) retention time: 4.0 min (99%).

**ChURRO-5:** Enantiomers were purified via preparatory SFC (AD-H column, 40% IPA/$CO_2$, 35 mL/min, 210 nm).
**(*R*)-ChURRO-5** is the first eluting enantiomer on AD-H column. Enantiopurity was determined by analytical chiral SFC (AD-3 column, 40% IPA/$CO_2$, 2.5 mL/min, 210 nm) retention time: 3.8 min (99%).
**(*S*)-ChURRO-5** is the second eluting enantiomer on AD-H column. Enantiopurity was determined by analytical Chiral SFC (AD-3 column, 40% IPA/$CO_2$, 2.5 mL/min, 210 nm) retention time: 5.0 min (99%).

**VCD analysis**: Vibrational circular dichroism (VCD) measurements were conducted at BioTools (Jupiter, FL) by Jordan Nafie as follows:
Probe measurements: Dissolved E1 (first eluting enantiomer of ChURRO-probe) or E2 (second eluting enantiomer of ChURRO-probe) (7-10 mg) in 120-650 $\mu$L $CDCl_3$ (previously run through a plug of activated basic alumina) and transferred to a 100 $\mu$m $BaF_2$ IR cell. Simultaneous measurement of IR and VCD were taken for ~18 hours in one-hour blocks on a BioTools (Jupiter, FL) DualPEM FT-VCD spectrometer at 4 $cm^{-1}$ resolution and PEM maximum frequencies both set to 1400 $cm^{-1}$. The first block of IR was solvent and water vapor corrected, and offset to zero at 2000 $cm^{-1}$. VCD was block averaged and corrected by half difference ((E1 – E2) / 2). Noise spectra were block averaged and used as is.

To analyze the stability of the enantiomeric pairs of ChURRO-2, the IR and VCD were measured simultaneously of each enantiomer in $CD_3OD$ for 24 hours in 30 min blocks as described above. Various numbers of blocks were averaged to determine the optimal number of blocks required for reasonably good signal to noise ratio VCD data, arriving at 8 hours (16 blocks), resulting in 33 VCD spectra of 8 hour avg each. The raw VCD data was then baseline corrected using the half difference method as described above.

Calculations: The (*R*) enantiomer of each ChURRO-probe was constructed using BioTools (Jupiter, FL) ComputeVOA software. A thorough conformational search was performed at the molecular mechanics level (GMMX) using the MMF94 force field in a 7 kcal / mol energy window. The resulting 121 conformations were optimized at the DFT (Gaussian 09, Wallingford, CT) level (B3LYP / 6-31G(d) with CPCM - Chloroform) and the IR and VCD frequencies were also calculated at the same level. The conformations were separated into cis and trans oxaziridine isomers, and the 20 lowest energy conformers from each group (cis and trans) were re-optimized and frequencies calculated at the B3PW91 / cc-pVTZ level. The spectra from the resulting conformers were Boltzmann averaged using the electronic energy values and plotted using a line width of 5 $cm^{-1}$ HWHH, and an x-axis scaling factor of 0.976 for comparison with the experimental IR and VCD data. Of the geometric isomers, it was determined that the trans was a good match with the data and the cis a poor match.

Methionine adduct:
Measurements: Two samples were received which were derived from (*R*)- and (*S*)-ChURRO-2 starting material. Each sample (~10 mg) was separately dissolved in 125 – 150 $\mu$L $CDCl_3$ (previously run through a plug of activated basic alumina) and transferred to a 100 $\mu$m $BaF_2$ IR cell. Simultaneous measurement of IR and VCD were taken for ~18 hours in one-hour blocks on a BioTools (Jupiter, FL) DualPEM FT-VCD spectrometer at 4 $cm^{-1}$ resolution and PEM maximum frequencies both set to 1400 $cm^{-1}$. The first block of IR was solvent and water vapor corrected, and offset to zero at 2000 $cm^{-1}$. VCD was block averaged and corrected by subtraction of a solvent baseline which was previously acquired (18 hours averaged). Noise spectra were block averaged and used as is. The two samples had identical IR and nearly identical VCD – based on this, and the fact that the calculated (s*R*) and (s*S*) diastereomers had very different VCD spectra, the samples were determined to be identical.

Calculations: The (s*R*, *S*) and (s*S*, *S*) diastereomers were separately constructed using BioTools (Jupiter, FL) ComputeVOA software. A thorough conformational search was performed on each at the molecular mechanics level (GMMX) using the MMF94 force field in a 7 kcal / mol energy window. The resulting ~150 conformations for each diastereomer were optimized at the DFT (Gaussian 09, Wallingford, CT) level (B3LYP / 6-31G(d) with CPCM - Chloroform) and the IR and VCD frequencies were also calculated at the same level. The 20 lowest energy conformers from each group (s*R* or s*S*) were re-optimized and frequencies calculated at the B3LYP / 6-311G(3df,2pd) level. The spectra from the resulting conformers were Boltzmann averaged using the electronic energy values and plotted using a line width of 5 $cm^{-1}$ HWHH, and an x-axis scaling factor of 0.982 for comparison with the experimental IR and VCD data. The (s*S*, *S*) DFT spectra were a much better match to the experimental than the (s*R*, *S*).

**Diastereoselectivity studies**:

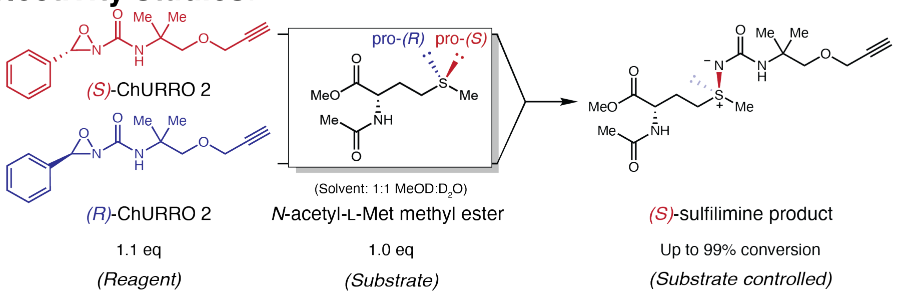


To an NMR tube was added (*R*)- or (*S*)-ChURRO-2 (9.1 mg, 0.033 mmol, 1.1 equiv.) and *N*-acetyl-L-methionine methyl ester (6.2 mg, 0.03 mmol, 1.0 equiv.) in 0.6 mL 1:1 $CD_3OD$:$D_2O$. The reaction was incubated and monitored by $^1H$ NMR. After 30 minutes the reaction was concentrated and purified using flash column chromatography (10% MeOH/DCM eluent) to give sulfilimine product which was analyzed by $^1H$ NMR, HRMS, VCD, and IR to confirm absolute configuration at sulfur.

**(*R*)-ChURRO-2 product:** 11.1 mg isolated. $^1H$ NMR (600 MHz, $CD_3OD$) δ 4.58 – 4.52 (m, 1H), 4.16 (d, *J* = 2.3 Hz, 2H), 3.74 (d, *J* = 1.4 Hz, 3H), 3.56 – 3.47 (m, 2H), 3.06 – 2.87 (m, 2H), 2.82 (t, *J* = 2.4 Hz, 1H), 2.63 (s, 3H), 2.30 – 2.23 (m, 1H), 2.10 – 2.02 (m, 1H), 2.01 (d, *J* = 3.0 Hz, 3H), 1.29 (d, *J* = 2.0 Hz, 6H, *overlap with grease*). HRMS (ESI): calculated for $[C_{16}H_{27}N_3O_5S+H]^+$: required *m/z* 374.1744, found *m/z* 374.1754.

**(*S*)-ChURRO-2 product:** 8.7 mg isolated. $^1H$ NMR (600 MHz, $CD_3OD$) δ 4.60 – 4.51 (m, 1H *overlap with impurity*), 4.16 (d, *J* = 2.4 Hz, 2H), 3.74 (d, *J* = 1.4 Hz, 3H), 3.56 – 3.47 (m, 2H), 3.07 – 2.87 (m, 2H), 2.82 (t, *J* = 2.4 Hz, 1H), 2.63 (s, 3H), 2.30 – 2.23 (m, 1H), 2.10 – 2.02 (m, 1H), 2.01 (d, *J* = 2.9 Hz, 3H), 1.29 (d, *J* = 2.0 Hz, 6H, *overlap with grease*). HRMS (ESI): calculated for $[C_{16}H_{27}N_3O_5S+H]^+$: required *m/z* 374.1744, found *m/z* 374.1754.

**Energies of Optimized Geometries**

| structure | energy in kcal | Delta G (R) | Delta G (S) |
|---|---|---|---|
| methionine | -622532.5111 | | |
| (*R*)-ChURRO-2 | -575332.7136 | | |
| (*S*)-ChURRO-2 | -575332.711 | | |
| *R* SMs | -1197865.225 | 0.00 | |
| *S* SMs | -1197865.222 | | 0.00 |
| *R*_s*S* TS1 | -1197841.608 | 23.62 | |
| *R*_s*R* TS1 | -1197839.958 | 25.27 | |
| *S*_s*R* TS1 | -1197836.37 | | 28.85 |
| *S*_s*S* TS1 | -1197841.263 | | 23.96 |

| | | | |
|---|---|---|---|
| *R*_s*R* int1 | -1197870.053 | -4.83 | |
| *S*_s*R* int1 | -1197876.253 | | -11.03 |
| *R*_s*S* int1 | -1197871.261 | -6.04 | |
| *S*_s*S*_int1 | -1197875.102 | | -9.88 |
| benzaldehyde | -216710.2069 | | |
| s*R* sulfilimine | -981174.3494 | | |
| s*S* sulfilimine | -981178.2042 | | |
| s*R* pdts | -1197884.556 | -19.33 | |
| s*S* pdts | -1197888.411 | | -23.19 |

**<u>XYZ Coordinates of Optimized Geometries</u>**

***N*-acetyl-L-methionine methyl ester**

| | | | |
|---|---|---|---|
| N | 0.65411900 | 1.28373700 | -0.63063100 |
| H | 0.67394800 | 1.50072700 | -1.62215700 |
| C | 0.20752600 | -0.03821200 | -0.25997900 |
| H | -0.22647500 | -0.01087900 | 0.74624800 |
| C | -0.86693200 | -0.52788700 | -1.23471700 |
| C | 1.37210700 | -1.02085700 | -0.20030600 |
| H | -0.44932300 | -0.54164600 | -2.24861000 |
| H | -1.12348200 | -1.55819400 | -0.97942900 |
| C | -2.12242000 | 0.33996300 | -1.23939900 |
| O | 2.52750200 | -0.74691000 | -0.44520300 |
| H | -1.90358500 | 1.34430500 | -1.61079500 |
| H | -2.86359700 | -0.10785400 | -1.90675500 |
| S | -2.88192900 | 0.58957700 | 0.39883800 |
| C | -3.08367700 | -1.13970500 | 0.91043700 |
| H | -2.11972500 | -1.61318600 | 1.11260800 |
| H | -3.67292400 | -1.13710400 | 1.82885200 |
| H | -3.62096700 | -1.70526300 | 0.14567900 |
| C | 1.39008400 | 2.11387000 | 0.15180500 |
| O | 1.90607900 | 3.12942500 | -0.32064700 |
| C | 1.50547500 | 1.76242900 | 1.61243800 |
| H | 0.51396600 | 1.67082700 | 2.06495300 |
| H | 2.02875800 | 0.81108300 | 1.74297300 |
| H | 2.06385000 | 2.55160300 | 2.11320800 |
| O | 0.96357100 | -2.22908100 | 0.18730600 |
| C | 1.99157200 | -3.22648400 | 0.30399000 |
| H | 1.48659400 | -4.13253000 | 0.63139400 |
| H | 2.47201100 | -3.38397500 | -0.66284300 |
| H | 2.73335600 | -2.91347100 | 1.04043000 |

**(*R*)-ChURRO-2**

| | | | |
|---|---|---|---|
| O | -0.75286600 | -2.01503000 | -1.28781200 |
| C | -0.73823000 | -1.62652100 | -0.12372400 |
| N | -1.49884200 | -0.67429500 | 0.41967100 |
| C | -2.54546200 | 0.07552400 | -0.30826300 |
| C | -3.47507900 | -0.88285900 | -1.05975100 |
| H | -4.35977100 | -0.33097900 | -1.38772200 |
| H | -2.99422800 | -1.32019100 | -1.93542800 |
| H | -3.79658200 | -1.68899300 | -0.39502900 |
| C | -3.34987100 | 0.84248800 | 0.73808100 |
| H | -3.87264800 | 0.14228700 | 1.39580600 |
| H | -4.09182300 | 1.46895400 | 0.23660600 |
| H | -2.70751300 | 1.48497300 | 1.34422400 |
| C | -1.91757400 | 1.03797400 | -1.32715400 |
| O | -1.33396900 | 2.20605600 | -0.77154300 |
| C | -0.04264300 | 2.04045400 | -0.20766500 |
| C | -0.08010900 | 1.93413100 | 1.26165500 |
| C | -0.12016800 | 1.85755000 | 2.46610100 |
| H | -0.15078100 | 1.78926800 | 3.53395200 |
| H | 0.54989300 | 2.92037600 | -0.47967100 |
| H | 0.45425600 | 1.15664300 | -0.62800000 |
| H | -1.18036600 | 0.49685600 | -1.93604000 |
| H | -2.71414200 | 1.39861400 | -1.98479100 |
| H | -1.28815600 | -0.39588600 | 1.37193500 |
| N | 0.07591300 | -2.38132600 | 0.80051000 |
| O | 0.50089800 | -1.59383500 | 1.95780400 |
| C | 1.43868600 | -1.96146700 | 0.97130200 |
| C | 2.06291400 | -0.85994800 | 0.18326000 |
| C | 2.59131100 | 0.23478800 | 0.86888600 |
| C | 3.21671400 | 1.25913800 | 0.16276500 |
| C | 3.31959000 | 1.18632000 | -1.22581900 |
| C | 2.80694800 | 0.08378700 | -1.90794000 |
| C | 2.18390000 | -0.94560900 | -1.20558300 |
| H | 1.78240400 | -1.80612900 | -1.73050600 |
| H | 2.89178300 | 0.02362100 | -2.98787700 |
| H | 3.80212100 | 1.98756600 | -1.77612200 |
| H | 3.61819100 | 2.11493300 | 0.69569800 |
| H | 2.49824600 | 0.28743900 | 1.94934100 |
| H | 2.10155500 | -2.77202700 | 1.26786000 |

**(*S*)-ChURRO-2**

| | | | |
|---|---|---|---|
| O | 0.75286000 | -2.01424800 | -1.28854500 |
| C | 0.73805400 | -1.62625400 | -0.12427900 |

| | | | |
|---|---|---|---|
| N | 1.49877100 | -0.67441400 | 0.41970000 |
| C | 2.54560300 | 0.07554400 | -0.30781400 |
| C | 3.34968600 | 0.84243500 | 0.73880000 |
| H | 4.09191500 | 1.46876500 | 0.23757200 |
| H | 2.70722300 | 1.48502000 | 1.34471400 |
| H | 3.87211900 | 0.14215800 | 1.39670900 |
| C | 3.47552200 | -0.88278600 | -1.05903200 |
| H | 2.99499600 | -1.32011700 | -1.93488500 |
| H | 3.79681600 | -1.68892600 | -0.39422100 |
| H | 4.36029700 | -0.33086300 | -1.38669100 |
| C | 1.91814400 | 1.03800900 | -1.32691200 |
| O | 1.33401800 | 2.20602000 | -0.77168000 |
| C | 0.04275500 | 2.04003400 | -0.20777100 |
| C | 0.08027600 | 1.93346000 | 1.26151000 |
| C | 0.12037100 | 1.85704500 | 2.46595900 |
| H | 0.15097500 | 1.78868800 | 3.53381100 |
| H | -0.45407400 | 1.15629400 | -0.62836800 |
| H | -0.54992600 | 2.91995600 | -0.47948100 |
| H | 2.71505300 | 1.39873300 | -1.98406600 |
| H | 1.18138800 | 0.49680500 | -1.93626900 |
| H | 1.28783000 | -0.39620300 | 1.37198700 |
| N | -0.07620800 | -2.38157800 | 0.79933800 |
| O | -0.50143400 | -1.59499400 | 1.95707200 |
| C | -1.43905800 | -1.96217900 | 0.97020300 |
| C | -2.06323600 | -0.86014400 | 0.18290100 |
| C | -2.18462700 | -0.94498100 | -1.20592500 |
| C | -2.80743500 | 0.08505400 | -1.90755600 |
| C | -3.31941200 | 1.18742600 | -1.22469100 |
| C | -3.21605500 | 1.25945900 | 0.16389800 |
| C | -2.59093100 | 0.23445300 | 0.86927800 |
| H | -2.49741700 | 0.28652700 | 1.94973000 |
| H | -3.61701500 | 2.11513600 | 0.69740200 |
| H | -3.80177200 | 1.98920400 | -1.77436100 |
| H | -2.89251500 | 0.02548200 | -2.98749800 |
| H | -1.78350700 | -1.80529000 | -1.73148500 |
| H | -2.10182000 | -2.77309500 | 1.26593900 |

**R_sR_TS1**

| | | | |
|---|---|---|---|
| O | -1.70246400 | -1.34129800 | -1.84486900 |
| C | -1.89360900 | -1.63528100 | -0.66984000 |
| N | -3.03841200 | -1.47421900 | 0.00094000 |
| C | -4.27074100 | -0.87188400 | -0.54215000 |
| C | -4.80240600 | -1.70201400 | -1.71137300 |
| H | -5.73470600 | -1.26428700 | -2.07853100 |

| | | | | |
|---|---|---|---|---|
| 604 | H | -4.08399100 | -1.73588300 | -2.53254200 |
| 605 | H | -5.00659900 | -2.72252800 | -1.37784200 |
| 606 | C | -5.28185300 | -0.85772300 | 0.60173000 |
| 607 | H | -5.49261200 | -1.87873500 | 0.93245800 |
| 608 | H | -6.21497200 | -0.40444200 | 0.25911300 |
| 609 | H | -4.90477500 | -0.28217200 | 1.45129000 |
| 610 | C | -4.00501600 | 0.55495400 | -1.02986600 |
| 611 | O | -3.49652300 | 1.33206400 | 0.04112800 |
| 612 | C | -3.22192700 | 2.66307700 | -0.36039700 |
| 613 | C | -2.50902100 | 3.35391500 | 0.71931800 |
| 614 | C | -1.88337900 | 3.91326000 | 1.58648300 |
| 615 | H | -1.34032300 | 4.42176000 | 2.35685600 |
| 616 | H | -4.15358900 | 3.19375500 | -0.59885800 |
| 617 | H | -2.59442600 | 2.66755100 | -1.26393200 |
| 618 | H | -3.28836700 | 0.55010500 | -1.86186600 |
| 619 | H | -4.95292700 | 0.97863900 | -1.39075700 |
| 620 | H | -3.04373300 | -1.76543300 | 0.97115700 |
| 621 | N | -0.84912900 | -2.19047900 | 0.20721400 |
| 622 | O | -0.75097000 | -3.84696000 | -0.71014700 |
| 623 | C | 0.17593900 | -2.84956300 | -0.58988100 |
| 624 | C | 1.43062200 | -3.13191100 | 0.19571800 |
| 625 | C | 2.60174100 | -2.43606300 | -0.10385900 |
| 626 | C | 3.75876800 | -2.65209500 | 0.64220100 |
| 627 | C | 3.74837300 | -3.57892600 | 1.68305400 |
| 628 | C | 2.58133800 | -4.28638400 | 1.97562500 |
| 629 | C | 1.42407300 | -4.06229700 | 1.23481000 |
| 630 | H | 0.51085300 | -4.60904900 | 1.45033900 |
| 631 | H | 2.57604000 | -5.01223700 | 2.78274100 |
| 632 | H | 4.64826300 | -3.75401200 | 2.26414000 |
| 633 | H | 4.66439100 | -2.09952000 | 0.40924200 |
| 634 | H | 2.60673500 | -1.72191600 | -0.92428700 |
| 635 | H | 0.41735500 | -2.31268900 | -1.51907900 |
| 636 | S | -0.03728600 | -0.25535500 | 0.96641300 |
| 637 | C | -0.26408200 | 1.03672900 | -0.29837000 |
| 638 | C | -1.31197000 | 0.17231200 | 2.17178600 |
| 639 | C | 0.97750600 | 1.11844100 | -1.17422100 |
| 640 | H | -0.45581500 | 1.97976100 | 0.22312900 |
| 641 | H | -1.14584900 | 0.78830500 | -0.89270900 |
| 642 | H | -2.24640700 | 0.40408800 | 1.65968400 |
| 643 | H | -1.43052900 | -0.69139800 | 2.82836400 |
| 644 | H | -0.97717200 | 1.03239900 | 2.75569500 |
| 645 | C | 2.14189400 | 1.83723300 | -0.46442100 |
| 646 | H | 0.73033500 | 1.68140400 | -2.08041200 |
| 647 | H | 1.29097100 | 0.11712100 | -1.48966700 |

| | | | |
|---|---|---|---|
| N | 1.85849400 | 3.24208100 | -0.27942400 |
| H | 2.33344900 | 1.36865700 | 0.50492000 |
| C | 3.40289100 | 1.66954000 | -1.29664600 |
| H | 1.77886000 | 3.79944700 | -1.12435800 |
| C | 1.52341100 | 3.87926700 | 0.86957800 |
| O | 3.90858100 | 2.53871900 | -1.97203900 |
| O | 3.85767900 | 0.42146800 | -1.21971400 |
| O | 1.17939900 | 5.06421400 | 0.84712400 |
| C | 1.61171100 | 3.10622400 | 2.16168200 |
| C | 5.00192400 | 0.11478100 | -2.03296800 |
| H | 1.36887900 | 3.78253100 | 2.97972600 |
| H | 0.90798800 | 2.26828100 | 2.17061300 |
| H | 2.61756000 | 2.70305900 | 2.30610200 |
| H | 5.19444800 | -0.94545600 | -1.88112400 |
| H | 4.78007900 | 0.31737900 | -3.08177700 |
| H | 5.85774100 | 0.71031700 | -1.71136800 |

**R_sS_TS1**

| | | | |
|---|---|---|---|
| O | -0.00549200 | -2.75687300 | 1.30268000 |
| C | 0.01748300 | -1.53644600 | 1.22309200 |
| N | -0.95069700 | -0.69154300 | 1.59284200 |
| C | -2.29850900 | -1.07080500 | 2.06364600 |
| C | -2.24428400 | -2.14741800 | 3.14729700 |
| C | -2.93477100 | 0.19571600 | 2.63215600 |
| C | -3.11111700 | -1.58501700 | 0.87367300 |
| O | -3.12306500 | -0.58367300 | -0.13401400 |
| C | -4.10440700 | -0.78937300 | -1.13352600 |
| C | -3.92449800 | -2.05059300 | -1.87300100 |
| C | -3.76742200 | -3.08855200 | -2.46803800 |
| N | 1.16853400 | -0.81553200 | 0.63308500 |
| O | 2.02458200 | -0.47652000 | 2.27484300 |
| C | 2.41040200 | -1.27490200 | 1.22906300 |
| C | 3.62786800 | -0.87590100 | 0.43336900 |
| C | 3.94999800 | 0.47094600 | 0.25832500 |
| C | 5.05620600 | 0.83305600 | -0.50469100 |
| C | 5.85119000 | -0.15138600 | -1.09258000 |
| C | 5.53719400 | -1.49701400 | -0.91479700 |
| C | 4.42748900 | -1.85723300 | -0.15223200 |
| S | 1.08786100 | -1.64285900 | -1.45269400 |
| C | 0.84886000 | 0.03054300 | -2.12625000 |
| C | -0.52233100 | -2.39791000 | -1.78092200 |
| C | -0.45756400 | 0.66999200 | -1.64483800 |
| C | -0.31085300 | 2.10623300 | -1.12522100 |
| N | 0.69276400 | 2.18412900 | -0.09023900 |

| | | | |
|---|---|---|---|
| C | -1.67055700 | 2.51195300 | -0.55523300 |
| C | 0.99462800 | 3.37492100 | 0.47587400 |
| O | -1.89890100 | 2.70795200 | 0.61930400 |
| O | -2.59903300 | 2.57654300 | -1.50734600 |
| O | 0.54311000 | 4.43296200 | 0.02721900 |
| C | 1.92972400 | 3.33820000 | 1.65623900 |
| C | -3.93096700 | 2.86969600 | -1.05210000 |
| H | -0.78818400 | 0.29537600 | 1.42077400 |
| H | -3.24518500 | -2.27075700 | 3.56930900 |
| H | -1.90753100 | -3.10783800 | 2.75776500 |
| H | -1.56926300 | -1.83457000 | 3.94833600 |
| H | -3.95258800 | -0.02837900 | 2.96051700 |
| H | -2.36191300 | 0.55295500 | 3.49249200 |
| H | -2.97573100 | 0.98767000 | 1.88120900 |
| H | -2.66415900 | -2.51001700 | 0.48210700 |
| H | -4.13594000 | -1.80338900 | 1.20517100 |
| H | -5.10837800 | -0.77346600 | -0.68838200 |
| H | -4.02121600 | 0.05472000 | -1.82437500 |
| H | -3.63227000 | -4.00718900 | -3.00090400 |
| H | 2.41667700 | -2.35503900 | 1.43591900 |
| H | 3.33572100 | 1.23149600 | 0.72893000 |
| H | 5.30212900 | 1.88194900 | -0.63732600 |
| H | 6.71564700 | 0.13100300 | -1.68496200 |
| H | 6.15600200 | -2.26539700 | -1.36689100 |
| H | 4.17674300 | -2.90547600 | -0.00986800 |
| H | 1.73499300 | 0.57574300 | -1.78784800 |
| H | 0.88991000 | -0.04049100 | -3.21483000 |
| H | -0.70368700 | -2.38468300 | -2.85837900 |
| H | -0.47510100 | -3.42688600 | -1.42317500 |
| H | -1.32040200 | -1.86336700 | -1.26413300 |
| H | -1.19269300 | 0.66905200 | -2.45214700 |
| H | -0.89473100 | 0.08808600 | -0.83099900 |
| H | -0.06482300 | 2.79559300 | -1.93905400 |
| H | 1.00257200 | 1.31906200 | 0.35108700 |
| H | 1.59185400 | 4.07127600 | 2.39008200 |
| H | 1.97802300 | 2.34879600 | 2.11524700 |
| H | 2.93116100 | 3.62751800 | 1.32364200 |
| H | -4.55725300 | 2.84962100 | -1.94134300 |
| H | -4.25528900 | 2.11185700 | -0.33539900 |
| H | -3.95725500 | 3.85617700 | -0.58658400 |

**S_sS_TS1**

| | | | |
|---|---|---|---|
| O | -1.38036100 | -0.93107900 | 0.10181300 |
| C | -1.79542800 | 0.15945600 | 0.48749400 |

| | | | | |
|---|---|---|---|---|
| 736 | N | -3.01279100 | 0.39437700 | 0.97815500 |
| 737 | C | -4.13783900 | -0.55292100 | 0.86457800 |
| 738 | C | -3.93580700 | -1.73413400 | 1.81140500 |
| 739 | C | -5.39687100 | 0.22751600 | 1.23375800 |
| 740 | C | -4.23896000 | -1.06520000 | -0.57881800 |
| 741 | O | -4.18014400 | 0.04632300 | -1.45868800 |
| 742 | C | -3.92918200 | -0.32058400 | -2.80220800 |
| 743 | C | -2.54380900 | -0.78071100 | -3.01613000 |
| 744 | C | -1.40839900 | -1.15401100 | -3.18866900 |
| 745 | N | -0.96248600 | 1.36751300 | 0.50527400 |
| 746 | O | -1.50880700 | 1.87292300 | -1.23957300 |
| 747 | C | -0.27634700 | 1.50769100 | -0.77533900 |
| 748 | C | 0.79778400 | 2.56483700 | -0.76243800 |
| 749 | C | 2.06828500 | 2.28416000 | -1.26410000 |
| 750 | C | 3.05692300 | 3.26851500 | -1.26356800 |
| 751 | C | 2.77270200 | 4.53695600 | -0.76565500 |
| 752 | C | 1.49712700 | 4.82489100 | -0.27523700 |
| 753 | C | 0.51163500 | 3.84416600 | -0.27895800 |
| 754 | S | 0.36641600 | 0.85241300 | 2.20526900 |
| 755 | C | 0.81353500 | -0.90793900 | 2.44582800 |
| 756 | C | -0.93687800 | 0.98745200 | 3.44935900 |
| 757 | C | 2.19778500 | -1.21672400 | 1.88840500 |
| 758 | C | 2.27945300 | -0.97452400 | 0.38114400 |
| 759 | N | 1.33972400 | -1.78464100 | -0.36925400 |
| 760 | C | 3.66718800 | -1.13640300 | -0.22781100 |
| 761 | C | 1.57762200 | -3.09844000 | -0.57199000 |
| 762 | O | 3.87769600 | -1.04485200 | -1.41984600 |
| 763 | O | 4.62121700 | -1.32006500 | 0.67804400 |
| 764 | O | 2.66180200 | -3.61340500 | -0.27375500 |
| 765 | C | 0.46918500 | -3.87560000 | -1.23120100 |
| 766 | C | 5.95271500 | -1.44886000 | 0.15449000 |
| 767 | H | -3.23483400 | 1.36793200 | 1.14909700 |
| 768 | H | -4.74474200 | -2.45818300 | 1.67699400 |
| 769 | H | -2.98433200 | -2.23429000 | 1.61602500 |
| 770 | H | -3.94996900 | -1.38842700 | 2.84852900 |
| 771 | H | -6.26020700 | -0.44146200 | 1.21149500 |
| 772 | H | -5.30564100 | 0.64379400 | 2.24109200 |
| 773 | H | -5.56795700 | 1.04438900 | 0.52717700 |
| 774 | H | -3.42182500 | -1.76161700 | -0.79470300 |
| 775 | H | -5.19259800 | -1.59829700 | -0.69767700 |
| 776 | H | -4.62638800 | -1.10254800 | -3.13038700 |
| 777 | H | -4.11108300 | 0.57054200 | -3.40818700 |
| 778 | H | -0.40057300 | -1.47967700 | -3.34758500 |
| 779 | H | 0.11316500 | 0.56346700 | -1.18673700 |

| | | | |
|---|---|---|---|
| H | 2.28461400 | 1.29412200 | -1.66134900 |
| H | 4.04418600 | 3.04349400 | -1.65409900 |
| H | 3.54039200 | 5.30422200 | -0.76291000 |
| H | 1.27385500 | 5.81641900 | 0.10566900 |
| H | -0.48650900 | 4.06266200 | 0.08902000 |
| H | 0.79307700 | -1.07337100 | 3.52623600 |
| H | 0.03200700 | -1.51842200 | 1.98289900 |
| H | -0.47932700 | 0.98970100 | 4.43951600 |
| H | -1.45764200 | 1.93004100 | 3.27731500 |
| H | -1.63215900 | 0.15075500 | 3.35437500 |
| H | 2.94765500 | -0.60353300 | 2.39448800 |
| H | 2.42447000 | -2.26633900 | 2.09944000 |
| H | 2.02928100 | 0.07447500 | 0.17694000 |
| H | 0.38630100 | -1.43693900 | -0.45521800 |
| H | -0.48996900 | -3.35678300 | -1.17744100 |
| H | 0.39173000 | -4.85309000 | -0.75306300 |
| H | 0.73082300 | -4.03085400 | -2.28181700 |
| H | 6.00711300 | -2.31436600 | -0.50808600 |
| H | 6.23073000 | -0.54540600 | -0.39033300 |
| H | 6.59683400 | -1.58805100 | 1.01974700 |

***S*_s*R*_TS1**

| | | | |
|---|---|---|---|
| O | 1.25219900 | -1.84412600 | -1.23755600 |
| C | 1.94150300 | -1.66875100 | -0.23523600 |
| N | 3.17798300 | -1.16452300 | -0.24505900 |
| C | 3.83689300 | -0.59737500 | -1.43333000 |
| C | 5.17202300 | -0.04084700 | -0.94190300 |
| C | 4.07186800 | -1.67001000 | -2.49773700 |
| C | 2.99347500 | 0.52816900 | -2.03582900 |
| O | 2.78295900 | 1.53128700 | -1.05458700 |
| C | 2.17333200 | 2.69068700 | -1.58510600 |
| C | 0.75745900 | 2.48012900 | -1.93308100 |
| C | -0.40109100 | 2.31033900 | -2.22744100 |
| N | 1.42620800 | -2.12986900 | 1.06459400 |
| O | 2.87937300 | -1.85947700 | 2.25524500 |
| C | 1.65969000 | -1.25468100 | 2.21801800 |
| C | 1.66984200 | 0.25010000 | 2.03564300 |
| C | 2.84993200 | 0.95903800 | 2.26281500 |
| C | 2.86990100 | 2.34779000 | 2.14804300 |
| C | 1.70976500 | 3.03573400 | 1.79749900 |
| C | 0.52766700 | 2.33152000 | 1.56922700 |
| C | 0.50615300 | 0.94520100 | 1.70349600 |
| S | -0.81439000 | -2.40840000 | 0.79408100 |
| C | -1.68580400 | -1.87101100 | -0.71624200 |

| | | | | |
|---|---|---|---|---|
| 824 | C | -0.51950600 | -4.12414900 | 0.32718500 |
| 825 | C | -2.05385100 | -0.39769400 | -0.67347500 |
| 826 | C | -3.07257200 | -0.06004500 | 0.42790300 |
| 827 | N | -4.26910700 | -0.87167700 | 0.36222200 |
| 828 | C | -3.45601300 | 1.41845700 | 0.45400900 |
| 829 | C | -5.15497200 | -0.73168800 | -0.64730900 |
| 830 | O | -4.49990700 | 1.84945800 | 0.89314000 |
| 831 | O | -2.47734600 | 2.19021000 | -0.00753300 |
| 832 | O | -4.99571400 | 0.11660000 | -1.53185700 |
| 833 | C | -6.34498600 | -1.65292300 | -0.63394100 |
| 834 | C | -2.72293400 | 3.60467300 | 0.02053900 |
| 835 | H | 3.64474500 | -1.10940400 | 0.65442500 |
| 836 | H | 5.70848300 | 0.41965700 | -1.77515300 |
| 837 | H | 5.01607200 | 0.71339800 | -0.16572900 |
| 838 | H | 5.78968900 | -0.84585400 | -0.53334200 |
| 839 | H | 4.60966800 | -1.23626500 | -3.34548000 |
| 840 | H | 4.67872000 | -2.47796400 | -2.08089000 |
| 841 | H | 3.12940100 | -2.08667700 | -2.85708600 |
| 842 | H | 3.54293800 | 0.94993800 | -2.88980900 |
| 843 | H | 2.03297800 | 0.14275900 | -2.39735900 |
| 844 | H | 2.24634200 | 3.46492500 | -0.81678300 |
| 845 | H | 2.71618600 | 3.03809900 | -2.47472600 |
| 846 | H | -1.42987300 | 2.16813700 | -2.48732600 |
| 847 | H | 0.95593000 | -1.54151400 | 3.01463000 |
| 848 | H | 3.75042700 | 0.41639500 | 2.53347000 |
| 849 | H | 3.79142000 | 2.89133200 | 2.33130800 |
| 850 | H | 1.72355200 | 4.11733100 | 1.70558000 |
| 851 | H | -0.38118400 | 2.86203100 | 1.30394400 |
| 852 | H | -0.42712200 | 0.40620500 | 1.56900700 |
| 853 | H | -1.02077600 | -2.07499200 | -1.55510000 |
| 854 | H | -2.57491400 | -2.50461400 | -0.79805600 |
| 855 | H | -1.47475100 | -4.61406800 | 0.13186600 |
| 856 | H | 0.11518500 | -4.14454500 | -0.56060500 |
| 857 | H | -0.01789300 | -4.61401400 | 1.16091700 |
| 858 | H | -1.15152800 | 0.20676300 | -0.54182500 |
| 859 | H | -2.48217200 | -0.13088100 | -1.64369800 |
| 860 | H | -2.62796200 | -0.24710100 | 1.41326900 |
| 861 | H | -4.42549100 | -1.57549200 | 1.07013400 |
| 862 | H | -6.33748100 | -2.23955600 | -1.55538200 |
| 863 | H | -6.35389100 | -2.32465200 | 0.22475300 |
| 864 | H | -7.25201100 | -1.04454300 | -0.63178600 |
| 865 | H | -1.83086800 | 4.06473400 | -0.40130200 |
| 866 | H | -3.59922500 | 3.84206000 | -0.58487300 |
| 867 | H | -2.87908600 | 3.93766300 | 1.04809400 |

***R*_s*R*_int1**

| | | | |
|---|---|---|---|
| O | 2.06926000 | 0.23751300 | 1.64108500 |
| C | 1.97518900 | -0.55538100 | 0.70616000 |
| N | 2.95900900 | -1.25204100 | 0.13612100 |
| C | 4.37455200 | -1.22545400 | 0.53389100 |
| C | 4.59918200 | -2.13411400 | 1.74266300 |
| H | 5.63879600 | -2.06827400 | 2.07740300 |
| H | 3.94638300 | -1.83561700 | 2.56814800 |
| H | 4.38364600 | -3.17292900 | 1.47965400 |
| C | 5.16501100 | -1.71652500 | -0.67768100 |
| H | 4.81508900 | -2.70615300 | -0.98683400 |
| H | 6.22498900 | -1.78847000 | -0.42246200 |
| H | 5.05070200 | -1.02516400 | -1.51761500 |
| C | 4.81855300 | 0.19372900 | 0.91106300 |
| O | 4.34697200 | 1.11577100 | -0.05742300 |
| C | 4.31073300 | 2.43874200 | 0.44958500 |
| C | 3.64322600 | 3.30485200 | -0.52661700 |
| C | 3.06898200 | 4.00747800 | -1.32107200 |
| H | 2.56882200 | 4.63501800 | -2.02974600 |
| H | 5.32560700 | 2.80889900 | 0.64834500 |
| H | 3.75182500 | 2.46432600 | 1.39567400 |
| H | 4.43145600 | 0.46093600 | 1.89860300 |
| H | 5.91674700 | 0.21092800 | 0.95199700 |
| H | 2.67179700 | -1.81289000 | -0.67873800 |
| N | 0.67267800 | -0.79406100 | 0.10641100 |
| O | 1.12596800 | -2.40526300 | -1.61464900 |
| C | 0.45395700 | -2.22519600 | -0.48387000 |
| C | -1.06542400 | -2.40726300 | -0.55911300 |
| C | -1.86112300 | -2.23496800 | 0.57862700 |
| C | -3.23748700 | -2.43625000 | 0.51793700 |
| C | -3.83119100 | -2.83079700 | -0.68317900 |
| C | -3.04321800 | -3.00791500 | -1.81802800 |
| C | -1.66524900 | -2.79322100 | -1.75548000 |
| H | -1.03820900 | -2.93239300 | -2.62990300 |
| H | -3.50086400 | -3.31264100 | -2.75424900 |
| H | -4.90319200 | -2.99485300 | -0.73228200 |
| H | -3.84534800 | -2.29084700 | 1.40719600 |
| H | -1.39737900 | -1.93137400 | 1.51437800 |
| H | 0.80541000 | -2.82446200 | 0.38463700 |
| S | 0.23694900 | 0.35775100 | -1.03805300 |
| C | 0.07796000 | 1.87614600 | -0.05002200 |
| C | 1.67975900 | 0.75380600 | -2.04037200 |

| | | | |
|---|---|---|---|
| C | -0.96335700 | 1.69896900 | 1.04562300 |
| H | -0.20088800 | 2.64291500 | -0.77875100 |
| H | 1.06190800 | 2.10971500 | 0.36268100 |
| H | 2.53137300 | 0.99820200 | -1.40241800 |
| H | 1.87363500 | -0.12766600 | -2.65075400 |
| H | 1.39841300 | 1.60702400 | -2.66291000 |
| C | -2.32809800 | 1.25665400 | 0.50614400 |
| H | -1.06658200 | 2.66130800 | 1.55774700 |
| H | -0.60754800 | 0.96814100 | 1.77359500 |
| N | -2.82644600 | 2.20231800 | -0.46212700 |
| H | -2.22017900 | 0.27490900 | 0.02726900 |
| C | -3.33958900 | 1.01371300 | 1.62485100 |
| H | -2.70972000 | 3.18661900 | -0.23997100 |
| C | -3.79011000 | 1.93717800 | -1.38756100 |
| O | -4.47422200 | 1.43949300 | 1.62440700 |
| O | -2.84085800 | 0.22615500 | 2.57282000 |
| O | -4.32916100 | 2.85447100 | -2.00721500 |
| C | -4.12281200 | 0.48987300 | -1.63822200 |
| C | -3.75344200 | -0.14399200 | 3.62061200 |
| H | -4.56821800 | 0.04007400 | -0.74572900 |
| H | -4.83190300 | 0.43449600 | -2.46280500 |
| H | -3.22322600 | -0.08158800 | -1.88915000 |
| H | -3.20084500 | -0.82510100 | 4.26379300 |
| H | -4.06209100 | 0.74314900 | 4.17571800 |
| H | -4.62798100 | -0.63928100 | 3.19565800 |

***S***_s***R***_int1

| | | | |
|---|---|---|---|
| O | 0.94485200 | -1.48844500 | -1.60894300 |
| C | 1.32904600 | -1.39063900 | -0.43992800 |
| N | 2.59595000 | -1.36779100 | -0.01783700 |
| C | 3.76559700 | -1.13362100 | -0.87598000 |
| C | 4.94164600 | -0.88167300 | 0.06528700 |
| C | 4.03787100 | -2.34558000 | -1.76774600 |
| C | 3.53241200 | 0.09023200 | -1.76212100 |
| O | 3.20563900 | 1.20888600 | -0.94998500 |
| C | 2.72405300 | 2.29412200 | -1.72107600 |
| C | 1.34575600 | 2.06226000 | -2.18981400 |
| C | 0.21622900 | 1.84897200 | -2.56041900 |
| N | 0.37889000 | -1.29282300 | 0.62143400 |
| O | 1.72287300 | -1.69768700 | 2.58267500 |
| C | 0.85274200 | -0.85319100 | 2.06178200 |
| C | 1.33726300 | 0.59590100 | 1.88557400 |
| C | 2.53058900 | 0.98748500 | 2.48886400 |
| C | 2.97894500 | 2.30412000 | 2.38285200 |

| | | | | |
|---|---|---|---|---|
| 956 | C | 2.22691400 | 3.24559600 | 1.68282600 |
| 957 | C | 1.02841400 | 2.85987300 | 1.08070300 |
| 958 | C | 0.59124500 | 1.54195700 | 1.17969900 |
| 959 | S | -0.85193700 | -2.40535100 | 0.63460100 |
| 960 | C | -1.90468700 | -2.03411000 | -0.81043600 |
| 961 | C | -0.13160200 | -3.98110900 | 0.15354500 |
| 962 | C | -2.16356800 | -0.54347200 | -0.92830800 |
| 963 | C | -2.88548500 | 0.05861400 | 0.28661100 |
| 964 | N | -4.10407000 | -0.63427900 | 0.63864400 |
| 965 | C | -3.15904400 | 1.55435800 | 0.11815500 |
| 966 | C | -5.19184100 | -0.57502100 | -0.16275500 |
| 967 | O | -4.06022400 | 2.14994200 | 0.66552500 |
| 968 | O | -2.24348100 | 2.14028200 | -0.64774100 |
| 969 | O | -5.16841900 | 0.04166300 | -1.23232800 |
| 970 | C | -6.42466700 | -1.28817700 | 0.32160300 |
| 971 | C | -2.34560300 | 3.56767300 | -0.77304400 |
| 972 | H | 2.71644700 | -1.44843700 | 0.99981000 |
| 973 | H | 5.85274200 | -0.72092700 | -0.51642400 |
| 974 | H | 4.75810600 | 0.00073100 | 0.68490000 |
| 975 | H | 5.09579300 | -1.74507800 | 0.71957300 |
| 976 | H | 4.90559400 | -2.15466500 | -2.40587900 |
| 977 | H | 4.24906700 | -3.22224000 | -1.14976300 |
| 978 | H | 3.17609100 | -2.56246700 | -2.40317100 |
| 979 | H | 4.45120200 | 0.29382800 | -2.33019600 |
| 980 | H | 2.72215000 | -0.10748000 | -2.47306800 |
| 981 | H | 2.74723500 | 3.17779700 | -1.07873700 |
| 982 | H | 3.37833800 | 2.47992900 | -2.58303200 |
| 983 | H | -0.78502600 | 1.67055000 | -2.89570400 |
| 984 | H | -0.13169500 | -0.80386400 | 2.57808400 |
| 985 | H | 3.10532200 | 0.24716200 | 3.03568200 |
| 986 | H | 3.91356400 | 2.59574900 | 2.85244100 |
| 987 | H | 2.57291500 | 4.27144800 | 1.60219200 |
| 988 | H | 0.44057600 | 3.58122100 | 0.51860700 |
| 989 | H | -0.33020700 | 1.24580400 | 0.68872400 |
| 990 | H | -1.38934100 | -2.41534000 | -1.69211400 |
| 991 | H | -2.81820800 | -2.60921200 | -0.62991700 |
| 992 | H | -0.94717900 | -4.70604400 | 0.10724000 |
| 993 | H | 0.35665600 | -3.87855300 | -0.81661500 |
| 994 | H | 0.57771200 | -4.25056800 | 0.93721600 |
| 995 | H | -1.21138000 | -0.02838600 | -1.07507800 |
| 996 | H | -2.77012100 | -0.39292800 | -1.82474300 |
| 997 | H | -2.22632500 | 0.00353400 | 1.16444700 |
| 998 | H | -4.17161300 | -1.09148900 | 1.53727600 |
| 999 | H | -7.21071600 | -0.54561600 | 0.47870900 |

| | | | |
|---|---|---|---|
| H | -6.76046700 | -1.97167400 | -0.46077700 |
| H | -6.25777900 | -1.84072800 | 1.24627700 |
| H | -1.51084300 | 3.86571200 | -1.40488900 |
| H | -3.29532100 | 3.83350900 | -1.23994800 |
| H | -2.27093900 | 4.03404800 | 0.21071600 |

***R*_s*S*_int1**

| | | | |
|---|---|---|---|
| O | 0.37677200 | -2.71268700 | -0.43469100 |
| C | 0.70061100 | -1.74166500 | 0.25063600 |
| N | 0.09065800 | -1.29178100 | 1.35183600 |
| C | -0.96108100 | -1.99208000 | 2.10654000 |
| C | -0.32274600 | -3.03159700 | 3.03030300 |
| H | -1.09513500 | -3.59879900 | 3.55837300 |
| H | 0.28914200 | -3.72844600 | 2.44967200 |
| H | 0.31269000 | -2.53751900 | 3.77018900 |
| C | -1.70625000 | -0.93130700 | 2.91735200 |
| H | -1.00512500 | -0.34175100 | 3.51742500 |
| H | -2.41161200 | -1.42176500 | 3.59315900 |
| H | -2.26249800 | -0.25366500 | 2.26182800 |
| C | -1.94020500 | -2.71723800 | 1.17916100 |
| O | -2.34685900 | -1.85031600 | 0.13274800 |
| C | -2.96150000 | -2.56965100 | -0.92327300 |
| C | -3.21577200 | -1.65946800 | -2.04218400 |
| C | -3.41371600 | -0.91454700 | -2.97012500 |
| H | -3.59911300 | -0.24397600 | -3.78387500 |
| H | -3.90392600 | -3.02286300 | -0.58687800 |
| H | -2.29930600 | -3.38011100 | -1.25888800 |
| H | -1.47270200 | -3.61144200 | 0.75707300 |
| H | -2.81058500 | -3.02712900 | 1.77492400 |
| H | 0.48553700 | -0.41675000 | 1.73467200 |
| N | 1.83458600 | -0.94533700 | -0.13446300 |
| O | 1.86629700 | 0.78163300 | 1.58400400 |
| C | 2.60088000 | -0.15415400 | 1.00856100 |
| C | 3.90396400 | 0.31229900 | 0.36176600 |
| C | 4.81311800 | -0.60815300 | -0.16827300 |
| C | 5.99406500 | -0.16947400 | -0.75980000 |
| C | 6.27761100 | 1.19583000 | -0.82329500 |
| C | 5.37741200 | 2.11506100 | -0.28999500 |
| C | 4.19443700 | 1.67239400 | 0.30128900 |
| H | 3.48280500 | 2.37344900 | 0.72470200 |
| H | 5.59603900 | 3.17755600 | -0.33388000 |
| H | 7.19873100 | 1.53879100 | -1.28370600 |

| | | | |
|---|---|---|---|
| H | 6.69526300 | -0.88993400 | -1.16919500 |
| H | 4.59187700 | -1.67164200 | -0.11547800 |
| H | 2.83670700 | -1.02734100 | 1.65307900 |
| S | 1.73600500 | -0.22443900 | -1.63164400 |
| C | 0.00099900 | 0.30494000 | -1.79433200 |
| C | 1.83417700 | -1.55315400 | -2.84112400 |
| C | -0.23405500 | 1.45150800 | -0.81051200 |
| H | -0.62329000 | -0.57233500 | -1.58944500 |
| H | -0.12540700 | 0.58887200 | -2.84273800 |
| H | 1.85084100 | -1.07615700 | -3.82324700 |
| H | 2.77671100 | -2.07142800 | -2.65774300 |
| H | 0.98215800 | -2.22131700 | -2.72902600 |
| C | -1.67245200 | 1.46216300 | -0.29115100 |
| H | -0.00122400 | 2.41057900 | -1.28382600 |
| H | 0.42908200 | 1.33984100 | 0.05538200 |
| N | -2.59392700 | 1.85227200 | -1.32947100 |
| H | -1.90550500 | 0.44540400 | 0.05410400 |
| C | -1.79562100 | 2.35666900 | 0.94166700 |
| H | -2.23619900 | 2.41309100 | -2.09494400 |
| C | -3.93236100 | 1.64000000 | -1.33210200 |
| O | -2.60786700 | 3.24469500 | 1.08633400 |
| O | -0.91270800 | 1.99879600 | 1.86800800 |
| O | -4.62927300 | 2.05893000 | -2.26163000 |
| C | -4.52592200 | 0.89265200 | -0.16585000 |
| C | -0.98675500 | 2.68394600 | 3.12587600 |
| H | -5.55905800 | 0.64721000 | -0.40845200 |
| H | -3.97138100 | -0.02364300 | 0.05265700 |
| H | -4.50566400 | 1.52611800 | 0.72609100 |
| H | -0.16398600 | 2.29065100 | 3.71978000 |
| H | -0.87091100 | 3.75848800 | 2.97683400 |
| H | -1.94417700 | 2.47466400 | 3.60734900 |

***S*_s*S*_int1**

| | | | |
|---|---|---|---|
| O | 1.48000700 | 1.38719000 | 0.84636900 |
| C | 1.76368400 | 0.20597500 | 0.62647100 |
| N | 2.97967800 | -0.29138100 | 0.39009300 |
| C | 4.16014500 | 0.50324000 | 0.01226600 |
| C | 4.55440500 | 1.47874200 | 1.12047700 |
| C | 5.28929700 | -0.49354800 | -0.23923500 |
| C | 3.83435700 | 1.27734200 | -1.26537400 |
| O | 3.47830100 | 0.34860300 | -2.27927300 |
| C | 2.69945000 | 0.92588900 | -3.30917900 |
| C | 1.32762500 | 1.23116200 | -2.86454900 |
| C | 0.20357100 | 1.48470300 | -2.50112700 |

| | | | |
|---|---|---|---|
| N | 0.73285000 | -0.79344100 | 0.62899900 |
| O | 1.88438500 | -2.88276800 | 0.35032700 |
| C | 1.00273300 | -2.08754800 | -0.22937200 |
| C | -0.39135800 | -2.65854300 | -0.48828700 |
| C | -1.26918300 | -2.00157100 | -1.35596900 |
| C | -2.56053200 | -2.48336600 | -1.55244000 |
| C | -2.98183700 | -3.63508900 | -0.88488400 |
| C | -2.10303600 | -4.30533500 | -0.03631300 |
| C | -0.81015100 | -3.81907100 | 0.15790800 |
| S | 0.11100800 | -1.17092500 | 2.13651000 |
| C | -1.00165700 | 0.19982000 | 2.55641500 |
| C | 1.44516500 | -0.91973500 | 3.31685600 |
| C | -2.18791800 | 0.26897900 | 1.59989300 |
| C | -1.84606000 | 0.96318400 | 0.27676400 |
| N | -1.32580700 | 2.30173000 | 0.45868000 |
| C | -3.00080100 | 0.99925400 | -0.71639400 |
| C | -2.14327700 | 3.31724200 | 0.80497000 |
| O | -2.96215000 | 1.62912000 | -1.75542800 |
| O | -4.01851400 | 0.22224400 | -0.36873000 |
| O | -3.36693000 | 3.15322400 | 0.88996400 |
| C | -1.49544500 | 4.64979000 | 1.06329800 |
| C | -5.11042100 | 0.16978200 | -1.30088700 |
| H | 3.01877400 | -1.31624300 | 0.32018500 |
| H | 5.47303000 | 2.00038800 | 0.83783700 |
| H | 3.77219400 | 2.21784100 | 1.30003800 |
| H | 4.74037300 | 0.92985200 | 2.04775600 |
| H | 6.18609400 | 0.04151700 | -0.56095100 |
| H | 5.52144500 | -1.04088200 | 0.67884300 |
| H | 5.01150200 | -1.21090100 | -1.01566700 |
| H | 3.00305800 | 1.96728300 | -1.07345300 |
| H | 4.70995900 | 1.86296300 | -1.57712200 |
| H | 3.17026600 | 1.84098500 | -3.69132000 |
| H | 2.66438400 | 0.19838100 | -4.12394500 |
| H | -0.80689600 | 1.72717700 | -2.22708900 |
| H | 1.33453000 | -1.58717000 | -1.16726000 |
| H | -0.93155800 | -1.11039100 | -1.88472800 |
| H | -3.23490100 | -1.96839900 | -2.23065400 |
| H | -3.98847600 | -4.01239100 | -1.03504500 |
| H | -2.42467500 | -5.20731900 | 0.47513200 |
| H | -0.11126400 | -4.32965000 | 0.81347000 |
| H | -1.33231300 | -0.04514500 | 3.57032600 |
| H | -0.41177900 | 1.11947000 | 2.58278500 |
| H | 1.02787000 | -1.09516400 | 4.31053100 |
| H | 2.20713500 | -1.66517400 | 3.08701400 |

| | | | |
|---|---|---|---|
| H | 1.83447600 | 0.09734100 | 3.23587700 |
| H | -2.57520400 | -0.73507700 | 1.39690000 |
| H | -2.98156100 | 0.82869200 | 2.10208900 |
| H | -1.05916600 | 0.39625500 | -0.22892000 |
| H | -0.31631200 | 2.40857600 | 0.46623600 |
| H | -0.41608300 | 4.62864500 | 0.91145600 |
| H | -1.71434600 | 4.94748900 | 2.09136200 |
| H | -1.94528800 | 5.38931200 | 0.39728300 |
| H | -5.58400800 | 1.15076800 | -1.36817800 |
| H | -4.75281400 | -0.13704200 | -2.28479100 |
| H | -5.80494400 | -0.56423300 | -0.89823300 |

**s*R*_sulfilimine**

| | | | |
|---|---|---|---|
| O | -3.81013000 | 0.41887100 | 0.83918100 |
| C | -2.81168400 | 0.15727500 | 0.12299800 |
| N | -2.87272400 | -0.93261700 | -0.70422800 |
| C | -1.77637600 | -1.63198900 | -1.38794800 |
| C | -2.35953600 | -2.95321600 | -1.89179300 |
| H | -1.59964100 | -3.50193300 | -2.45438900 |
| H | -2.69566800 | -3.57648100 | -1.05943700 |
| H | -3.20842400 | -2.76125300 | -2.55417700 |
| C | -1.27756400 | -0.81499400 | -2.57948000 |
| H | -1.04770700 | 0.21073700 | -2.29310800 |
| H | -2.05207900 | -0.78470600 | -3.35137400 |
| H | -0.37833000 | -1.26666400 | -3.01000600 |
| C | -0.60726400 | -1.92913300 | -0.44247100 |
| O | -1.08719900 | -2.68674800 | 0.65904400 |
| C | -0.05138100 | -3.25925600 | 1.43067800 |
| C | 0.66151700 | -2.27883600 | 2.26878500 |
| C | 1.25135300 | -1.50165600 | 2.98017300 |
| H | 1.77362300 | -0.81371400 | 3.61282400 |
| H | 0.67372800 | -3.77390100 | 0.78424200 |
| H | -0.51723400 | -4.00693400 | 2.07829100 |
| H | 0.14828600 | -2.50975500 | -0.99310400 |
| H | -0.14047600 | -1.00225700 | -0.08895500 |
| H | -3.69696400 | -1.48906400 | -0.51917000 |
| N | -1.68083900 | 0.91982600 | 0.07200400 |
| S | -1.74611100 | 2.05211200 | 1.28583100 |
| C | -0.04089000 | 2.66384400 | 1.23386000 |
| C | -2.59558000 | 3.48075100 | 0.59219000 |
| C | 0.93906900 | 1.50903800 | 1.47281300 |
| H | 0.01625300 | 3.42259200 | 2.01714300 |

| | | | |
|---|---|---|---|
| H | 0.10255800 | 3.14787000 | 0.26468900 |
| H | -2.18106200 | 3.68855000 | -0.39579000 |
| H | -3.65069100 | 3.21468400 | 0.52541200 |
| H | -2.45900200 | 4.32916900 | 1.26607800 |
| C | 1.92905200 | 1.33489500 | 0.30816700 |
| H | 0.39904500 | 0.56558800 | 1.60468100 |
| H | 1.51599000 | 1.68023600 | 2.38282000 |
| N | 1.22296000 | 1.25802700 | -0.94954800 |
| H | 2.61253800 | 2.19050900 | 0.30296000 |
| C | 2.79531200 | 0.11527000 | 0.61722000 |
| H | 0.22707900 | 1.03049500 | -0.90893500 |
| C | 1.78437400 | 1.43245200 | -2.16961700 |
| O | 3.59896500 | 0.10988400 | 1.52791800 |
| O | 2.57491900 | -0.91831500 | -0.18514000 |
| O | 1.11013600 | 1.35245900 | -3.20107200 |
| C | 3.25868600 | 1.74754400 | -2.21384900 |
| C | 3.36615000 | -2.09033600 | 0.07272300 |
| H | 3.84805500 | 0.97733200 | -1.70817500 |
| H | 3.56487500 | 1.80852600 | -3.25684000 |
| H | 3.45841600 | 2.70392800 | -1.72185500 |
| H | 3.03872800 | -2.83038100 | -0.65468500 |
| H | 4.42394300 | -1.85968600 | -0.06633200 |
| H | 3.19119700 | -2.44306500 | 1.09015800 |

**s*S*_sulfilimine**

| | | | |
|---|---|---|---|
| O | 1.75718200 | 1.92189900 | 1.26556400 |
| C | 1.44110200 | 1.67706000 | 0.08423000 |
| N | 2.22861600 | 0.93186200 | -0.74803100 |
| C | 3.43851000 | 0.21413300 | -0.32452600 |
| C | 3.99910600 | -0.46290500 | -1.57578400 |
| H | 4.86358900 | -1.07728400 | -1.31039700 |
| H | 3.24777900 | -1.10360700 | -2.04466000 |
| H | 4.31744600 | 0.29239300 | -2.29987400 |
| C | 4.48465300 | 1.18007700 | 0.23734700 |
| H | 4.16838600 | 1.60961100 | 1.18842900 |
| H | 4.65407400 | 1.99231100 | -0.47470200 |
| H | 5.42987300 | 0.65064900 | 0.38732300 |
| C | 3.12222100 | -0.83661900 | 0.74184400 |
| O | 2.32630900 | -1.86798100 | 0.17323500 |
| C | 1.97023600 | -2.85445200 | 1.12012900 |
| C | 0.94429500 | -2.37753200 | 2.06491500 |
| C | 0.10488700 | -1.97661900 | 2.83496900 |
| H | -0.63599300 | -1.62823400 | 3.52536100 |
| H | 2.85285100 | -3.18864500 | 1.68215300 |

| | | | |
|---|---|---|---|
| H | 1.58239500 | -3.71007800 | 0.56071400 |
| H | 4.06163100 | -1.26773000 | 1.11837700 |
| H | 2.59222700 | -0.36718700 | 1.57984500 |
| H | 1.76589100 | 0.56549400 | -1.57466800 |
| N | 0.29886700 | 2.11201600 | -0.55876700 |
| S | -0.71583000 | 2.93247000 | 0.44515700 |
| C | -2.14214300 | 1.80749300 | 0.64118300 |
| C | -1.45824400 | 4.10891200 | -0.68822800 |
| C | -1.71614100 | 0.46513900 | 1.25945800 |
| H | -2.58509200 | 1.69512300 | -0.35499300 |
| H | -2.85291000 | 2.34093400 | 1.27691600 |
| H | -2.32837300 | 4.55074600 | -0.19920800 |
| H | -0.71417700 | 4.87739800 | -0.89825100 |
| H | -1.74290700 | 3.58407800 | -1.60133300 |
| C | -1.76977600 | -0.70883800 | 0.26925600 |
| H | -2.37490500 | 0.23578200 | 2.09769900 |
| H | -0.69568100 | 0.50987300 | 1.65796700 |
| N | -0.76465000 | -0.55288300 | -0.75167400 |
| H | -1.59260100 | -1.62568400 | 0.84212600 |
| C | -3.16079800 | -0.85121000 | -0.34605100 |
| H | -0.45104900 | 0.40596300 | -0.94901400 |
| C | -0.21789500 | -1.51877800 | -1.51509800 |
| O | -3.38831600 | -0.94807900 | -1.53268500 |
| O | -4.10575600 | -0.87439700 | 0.59287500 |
| O | 0.60838000 | -1.22700900 | -2.39286900 |
| C | -0.65241900 | -2.94097400 | -1.27586100 |
| C | -5.45219500 | -1.03260300 | 0.11530500 |
| H | -0.00606600 | -3.60496100 | -1.84827200 |
| H | -0.60412400 | -3.20303100 | -0.21546000 |
| H | -1.68812200 | -3.06898400 | -1.60512400 |
| H | -6.08094000 | -1.03396500 | 1.00282500 |
| H | -5.71568700 | -0.20083800 | -0.53982100 |
| H | -5.54885200 | -1.97577100 | -0.42464900 |

**Benzaldehyde**

| | | | |
|---|---|---|---|
| C | 2.20777400 | -0.24645000 | 0.00000200 |
| C | 1.72658000 | 1.06171400 | 0.00000200 |
| C | 0.35390400 | 1.29068900 | -0.00000100 |
| C | -0.53047800 | 0.20962300 | -0.00000300 |
| C | -0.04595800 | -1.10329700 | -0.00000300 |
| C | 1.32366900 | -1.32888600 | 0.00000000 |
| H | 3.27808200 | -0.42668100 | 0.00000400 |
| H | 2.41862600 | 1.89699900 | 0.00000400 |
| H | -0.03922800 | 2.30397500 | -0.00000100 |

| | | | |
|---|---|---|---|
| H | -0.75056700 | -1.92919300 | -0.00000500 |
| H | 1.70853500 | -2.34316800 | 0.00000000 |
| C | -1.98149200 | 0.47348100 | -0.00000600 |
| O | -2.83386900 | -0.39800400 | 0.00000600 |
| H | -2.26850100 | 1.54086200 | 0.00000100 |

**Table S1.**

Proteomic data containing shotgun proteomics data of proteins with evidence for stereospecific MetO reduction treated with (*R*)- or (*S*)-ChURRO-2, identified prochiral Met oxidation sites in oxidatively-stressed HEK-293T cells with MsrA or MsrB2 knocked down, quantitative isoDTB proteomic data in HEK-293T cell lysates for (*R*)- or (*S*)-ChURRO-2, identified prochiral Met oxidation sites in purified mitochondria derived from mouse liver, isoTOP-ABPP hyper-reactivity data of Met sites in purified mitochondria derived from mouse liver, BPHL shotgun proteomics with ChURRO-2, and $^{18}O$-$H_2O_2$ proteomics shotgun proteomics with BPHL, and isoTOP-ABPP hyper-reactivity of *N*-Hcylated sites in HEK-293T cells.

**Table S2.**

Quantification data containing protein labeling data for all 5 ChURRO-probes determined by densitomtry, cell viability data for live cell labeling of ChURRO-2 in HEK-293T cells and densitometry quantification of dot blot to assess probe internalization, Michaelis-Menten kinetic data for all purified hBPHL variants, relative activity of WT hBPHL oxidative stress treated with (*R*)- or (*S*)-ChURRO-2 and (*R*)- or (*S*)-OT-Ox used to compute dose-response curves and $IC_{50}$ values, peptide spectrum matches for shotgun proteomics analysis of WT hBPHL treated with racemic (*R*)- or (*S*)-ChURRO-2, densitometry quantification of BPHL expression in a panel of renal and hepatic cell lines, SOD1 cellular activity assays, densitometry quantification of protein *N*-homocysteinylation with MsrA or MsrB2 KO, and densitometry quantification of BPHL Co-IP with MsrA or MsrB2.

**Table S3.**

Bioinformatic data containing druggability analysis identified stereoselective Met oxidation sites for all ChURRO-2 probe pairs according to ChEMBL database, extracted phi and psi angles and Ramachandran analysis of identified Met oxidation sites for ChURRO-2 probe pairs, crowdedness analysis of identified Met oxidation sites for ChURRO-2 probe pairs by predicted partial sphere exposure analysis, subcellular distribution of identified prochiral Met oxidation sites in purified mitochondria derived from mouse liver, AlphaMissense analysis of identified prochiral Met oxidation sites in purified mitochondria derived from mouse liver, and gene ontology analysis of identified prochiral Met oxidation sites in purified mitochondria derived from mouse liver.

1308

### NMR Spectra

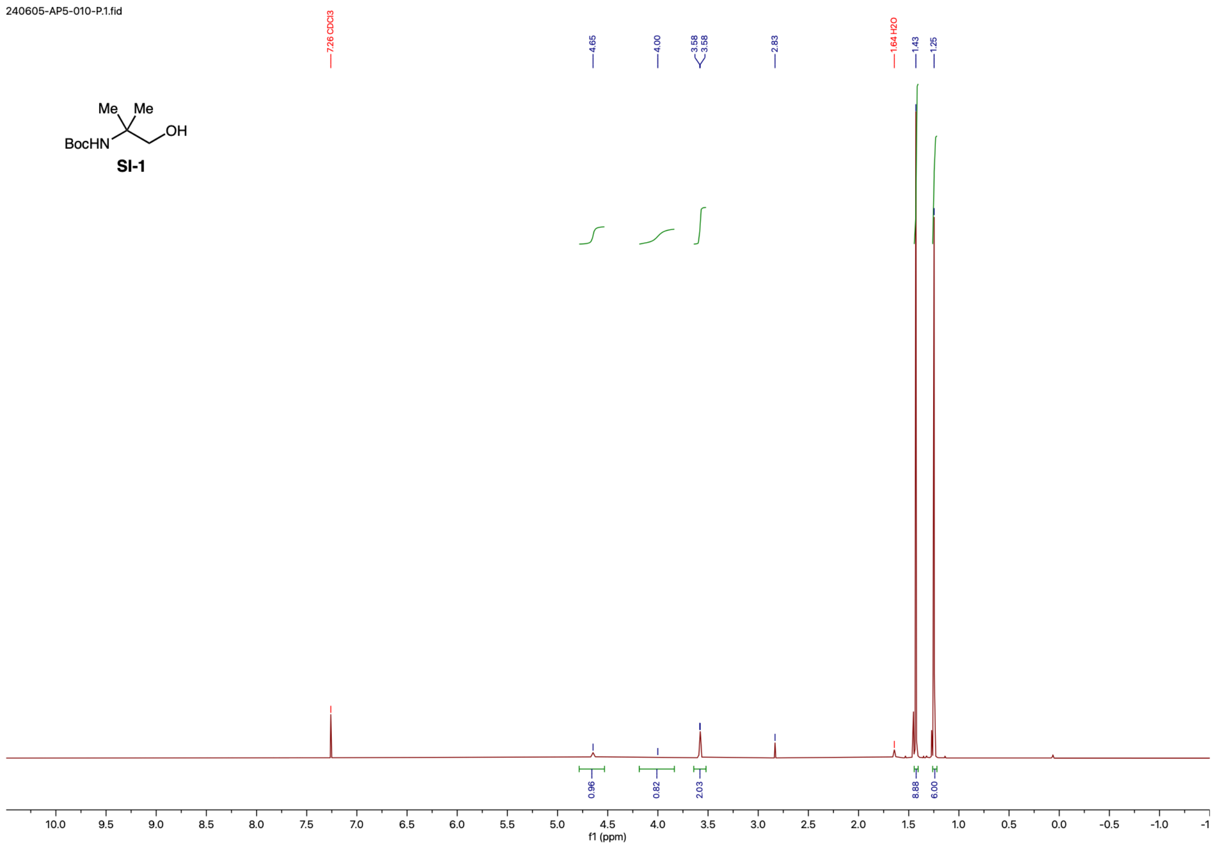

240605-AP5-010-P.1.fid
7.26 CDCl3
4.65
4.00
3.58
3.58
2.83
1.64 H2O
1.43
1.25
Me Me
BocHN
OH
SI-1
0.96
0.82
2.03
8.88
6.00
f1 (ppm)

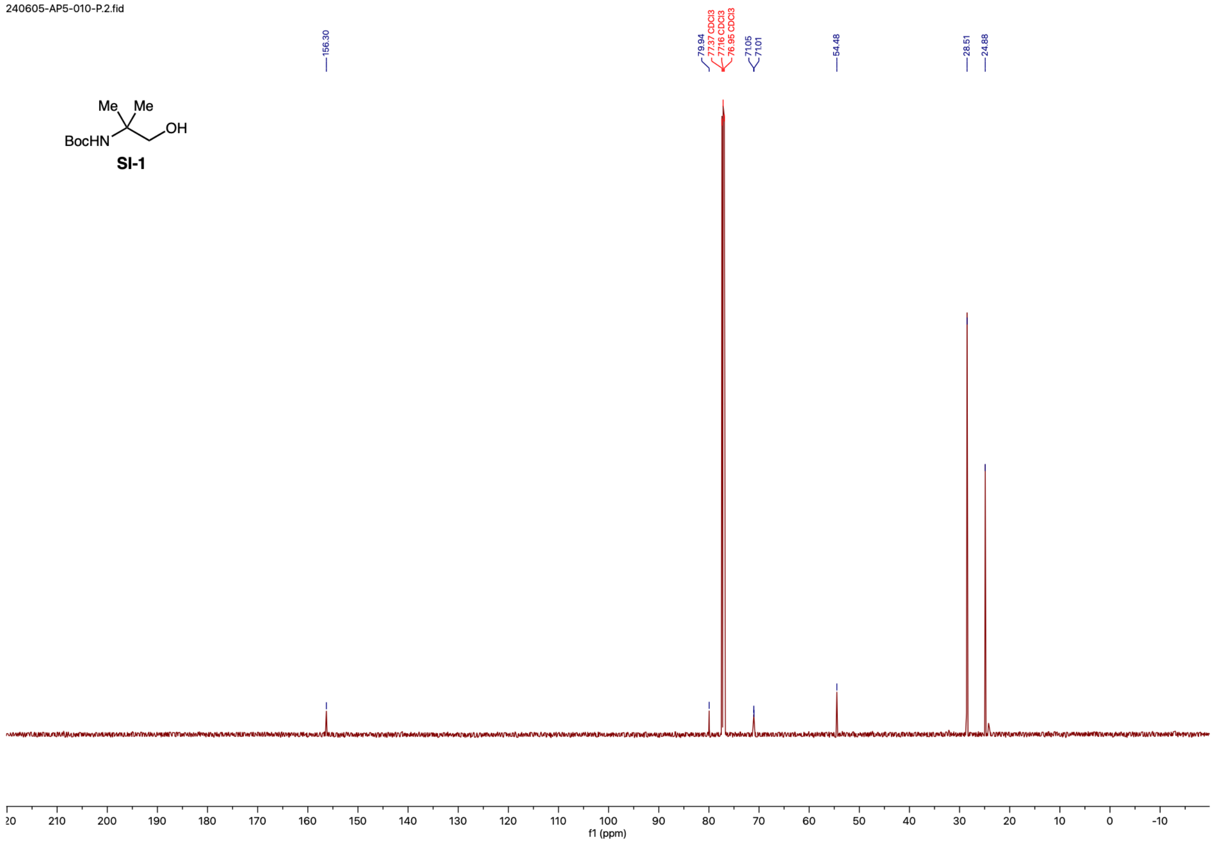

240605-AP5-010-P.2.fid
156.30
79.94
77.37 CDCl3
77.16 CDCl3
76.95 CDCl3
71.05
71.01
54.48
28.51
24.88
Me
Me
BocHN
OH
SI-1
f1 (ppm)


1312

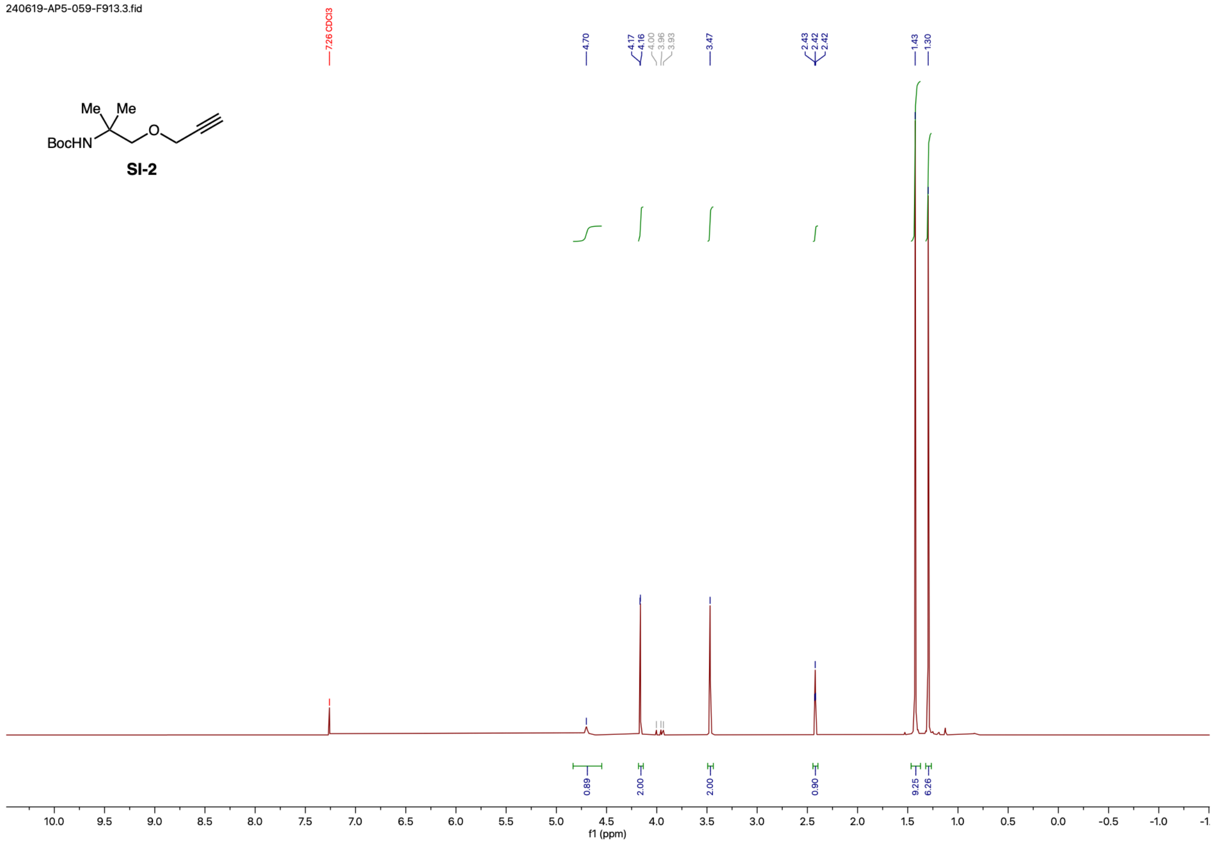
240619-AP5-059-F913.3.fid
7.26 CDCl3
4.70
4.17
4.16
4.00
3.96
3.93
3.47
2.43
2.42
2.42
1.43
1.30
Me Me
BocHN
O
SI-2
0.89
2.00
2.00
0.90
9.25
6.26
10.0 9.5 9.0 8.5 8.0 7.5 7.0 6.5 6.0 5.5 5.0 4.5 4.0 3.5 3.0 2.5 2.0 1.5 1.0 0.5 0.0 -0.5 -1.0 -1
f1 (ppm)

1313

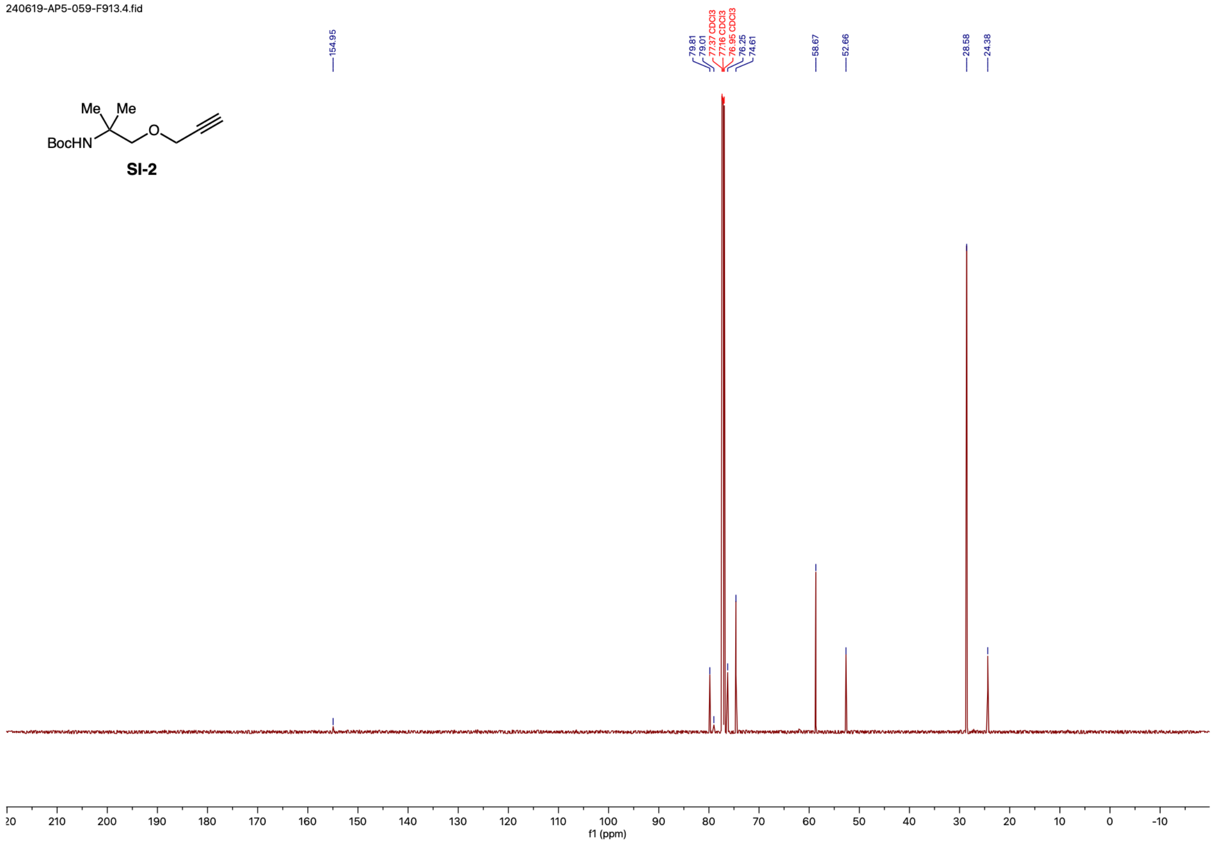
240619-AP5-059-F913.4.fid
154.95
79.81
79.01
77.37 CDCl3
77.16 CDCl3
76.95 CDCl3
76.25
74.61
58.67
52.66
28.58
24.38
Me Me
BocHN
O
SI-2
f1 (ppm)

1314

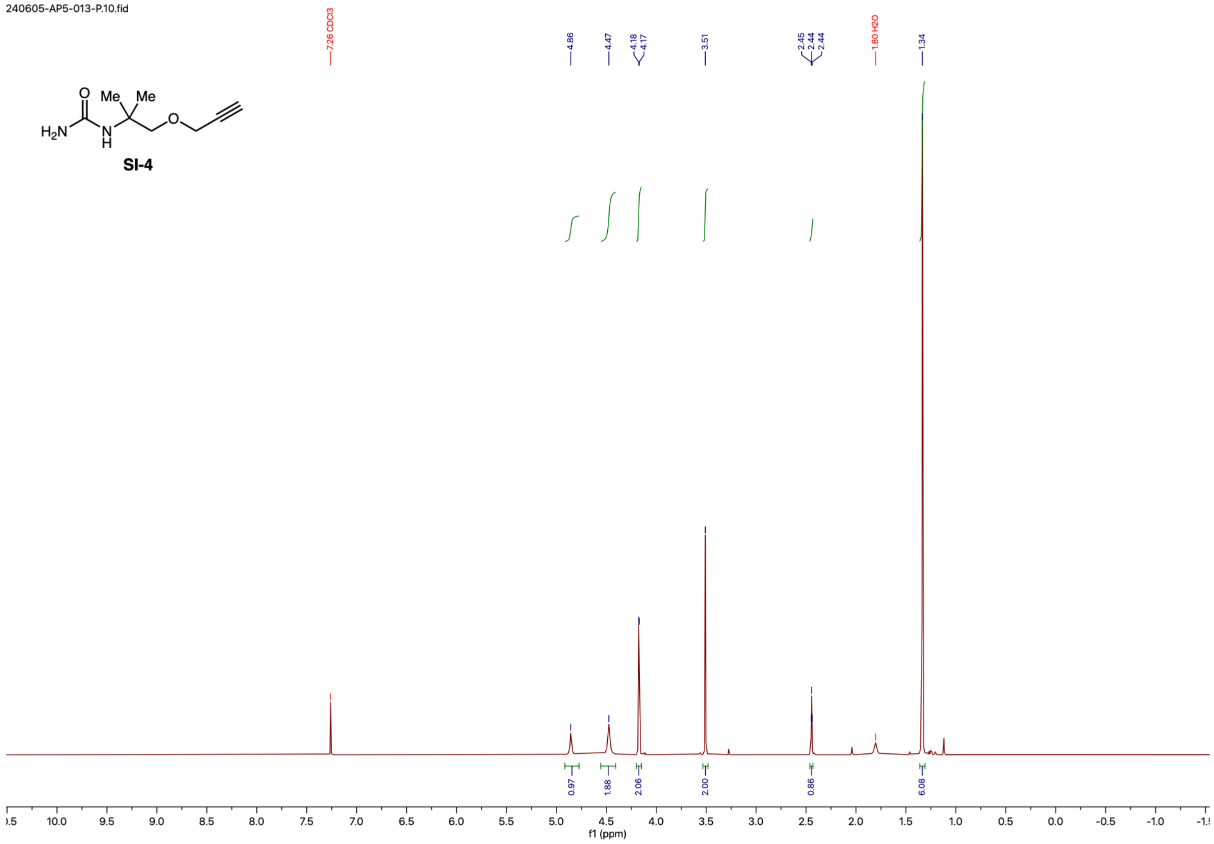
240605-AP5-013-P.10.fid
SI-4
Me
Me
$H_2N$
O
N
H
7.26 CDCl3
4.86
4.47
4.18
4.17
3.51
2.45
2.44
2.44
1.80 H2O
1.34
0.97
1.88
2.06
2.00
0.86
6.08
f1 (ppm)

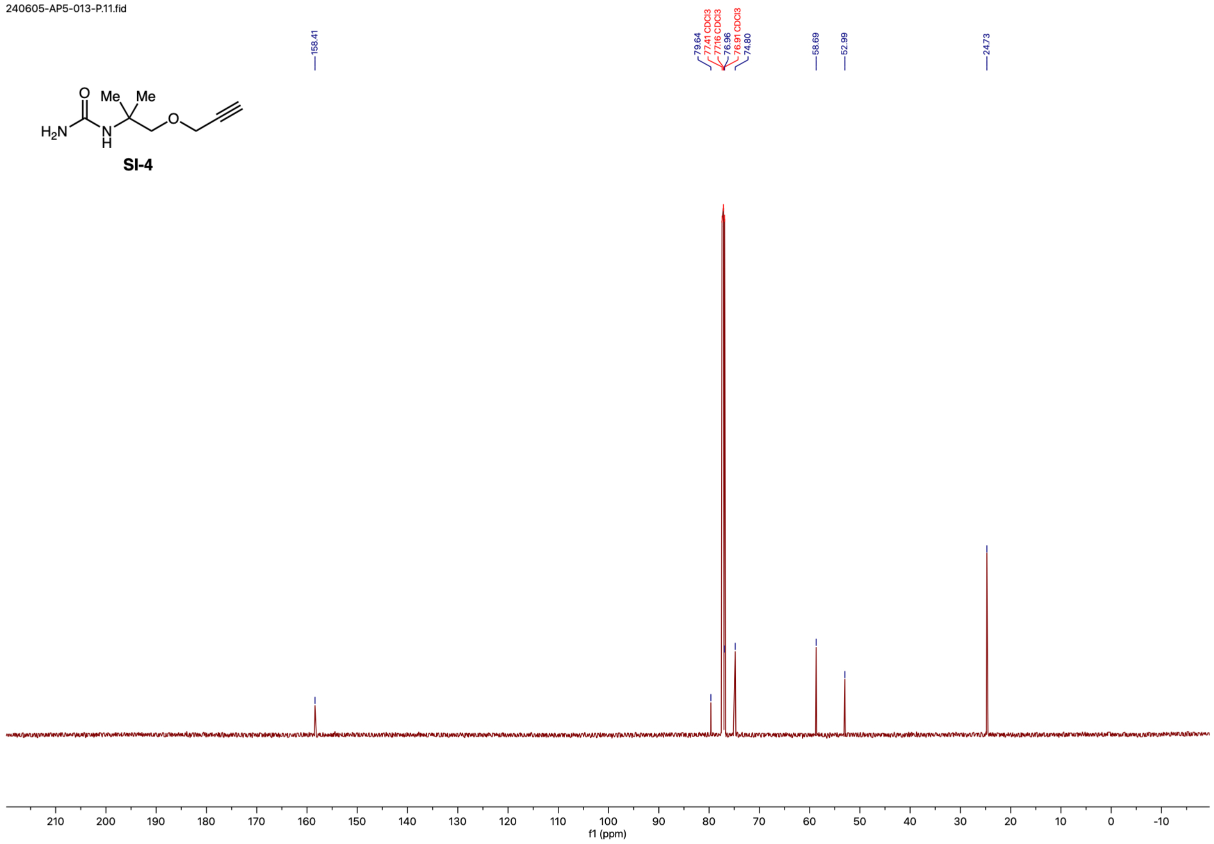

240605-AP5-013-P.11.fid
158.41
79.64
77.41 CDCl3
77.16 CDCl3
76.96
76.91 CDCl3
74.80
58.69
52.99
24.73
Me
Me
O
$H_2N$
N
H
O
SI-4
210 200 190 180 170 160 150 140 130 120 110 100 90 80 70 60 50 40 30 20 10 0 -10
f1 (ppm)


1316

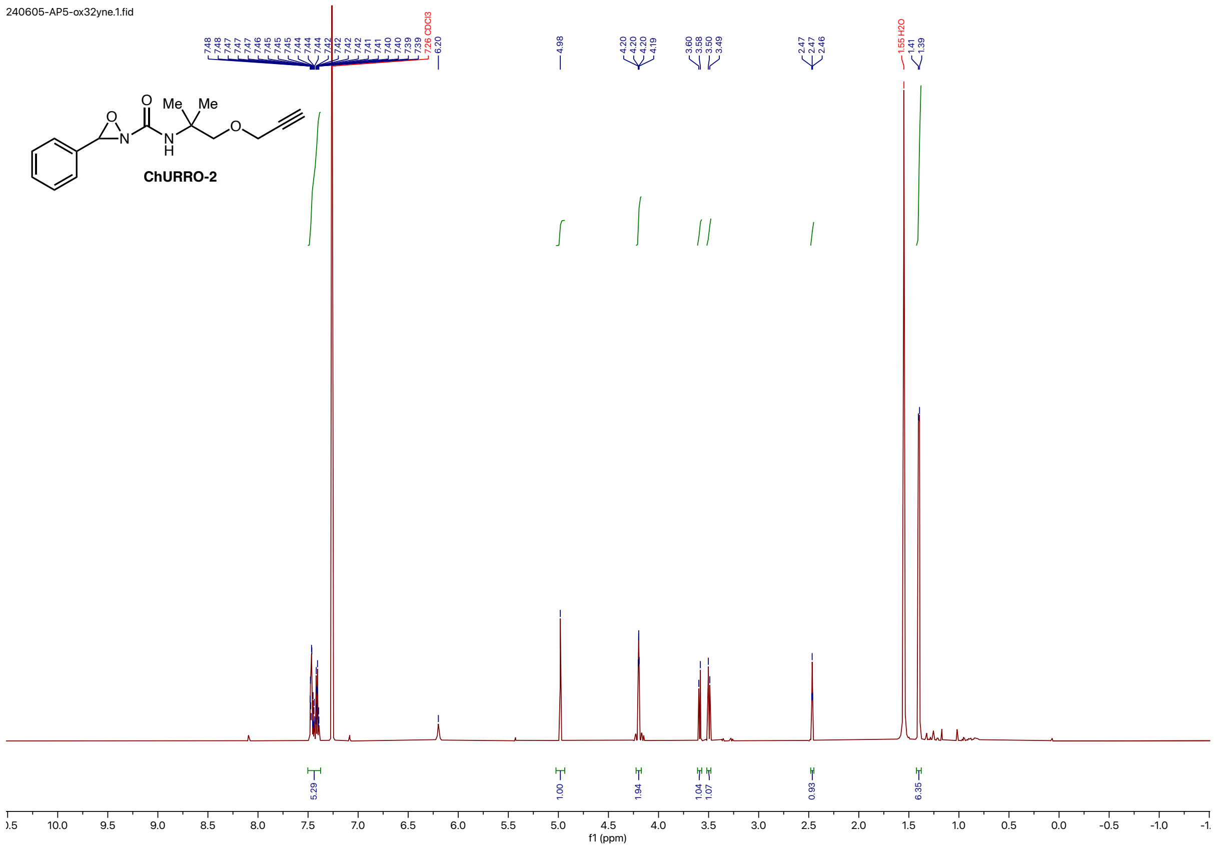


1317

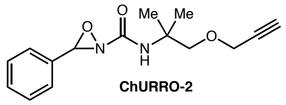

**ChURRO-2**

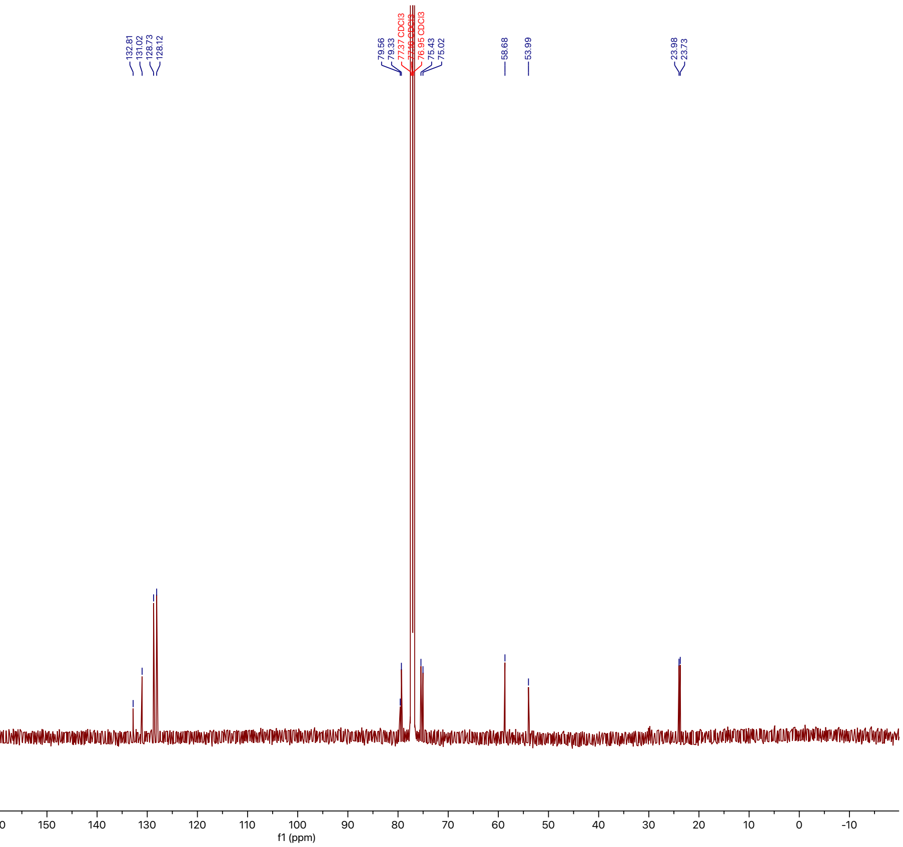


1318

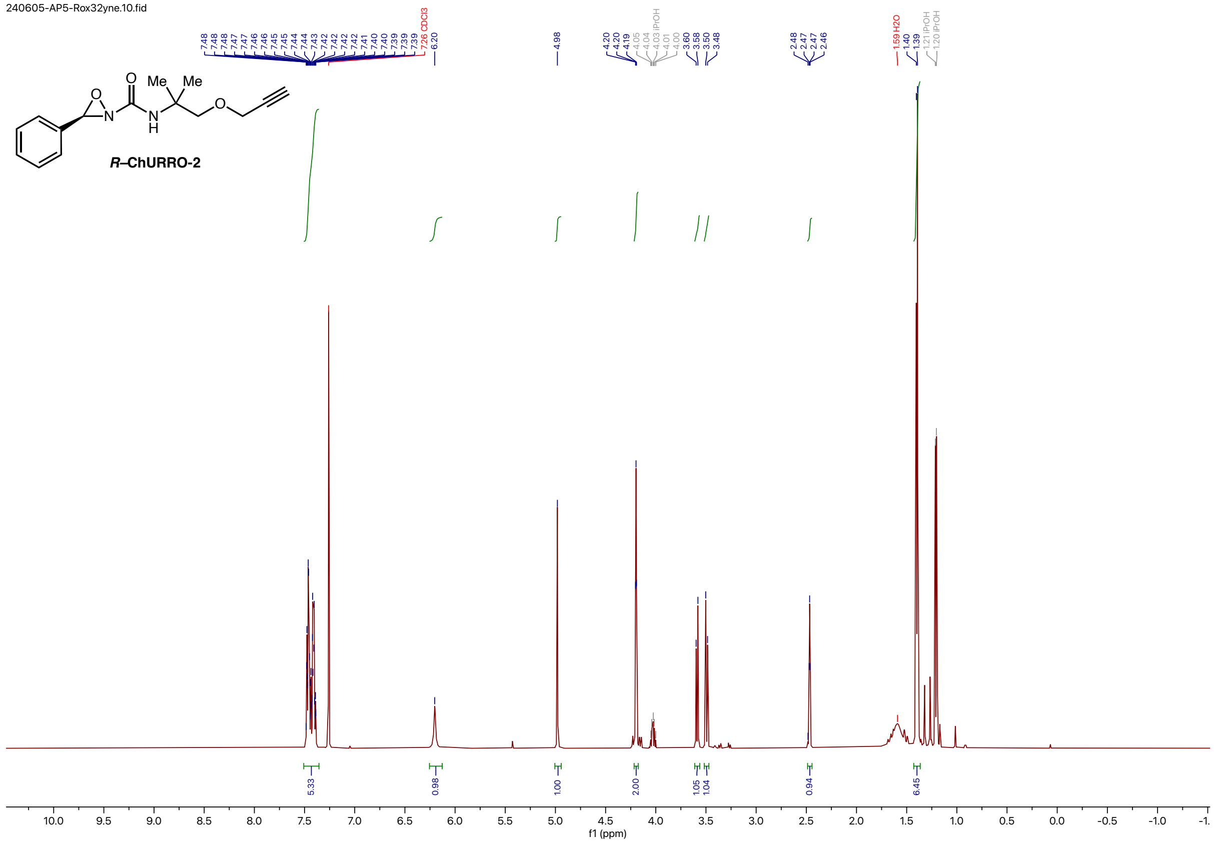


1319

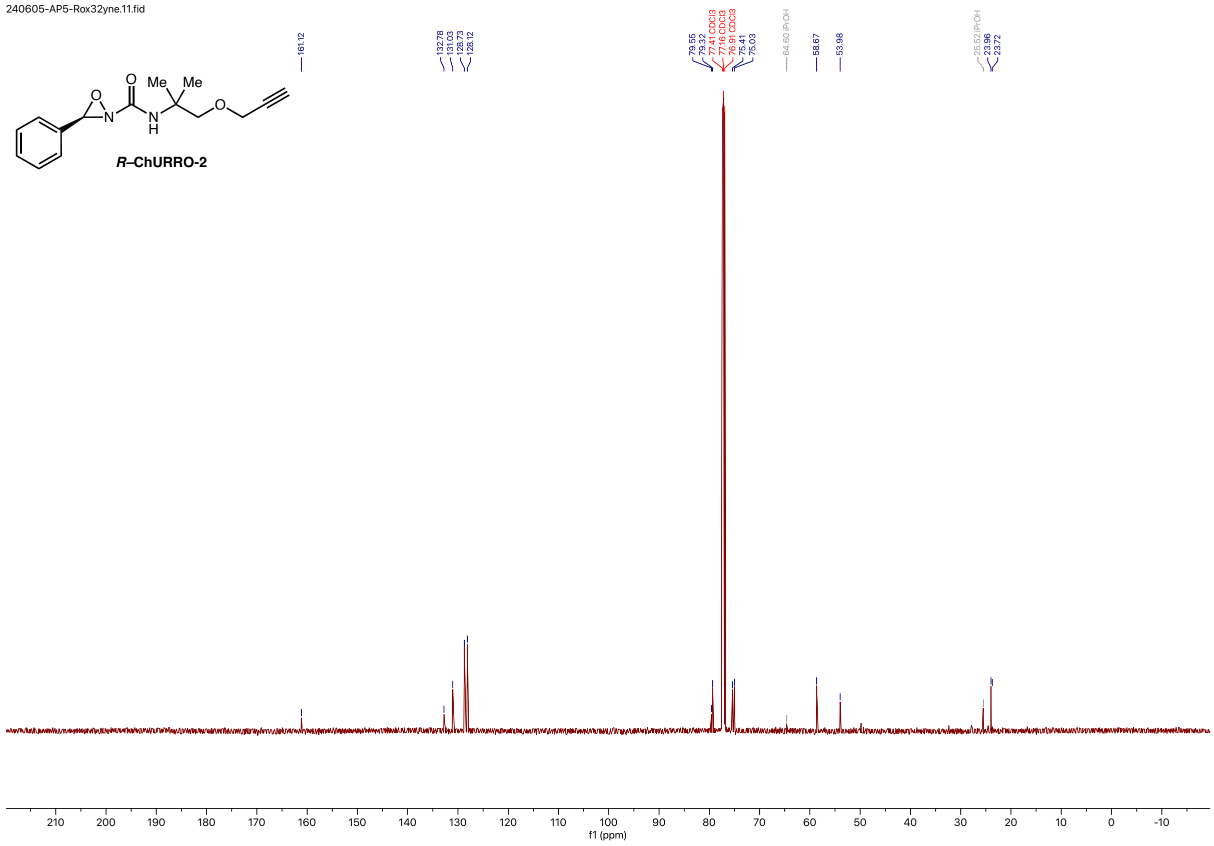
240605-AP5-Rox32yne.11.fid
161.12
132.78
131.03
128.73
128.12
79.55
79.32
77.41 CDCl3
77.16 CDCl3
76.91 CDCl3
75.41
75.03
64.60 iPrOH
58.67
53.98
25.52 iPrOH
23.96
23.72
R–ChURRO-2
f1 (ppm)

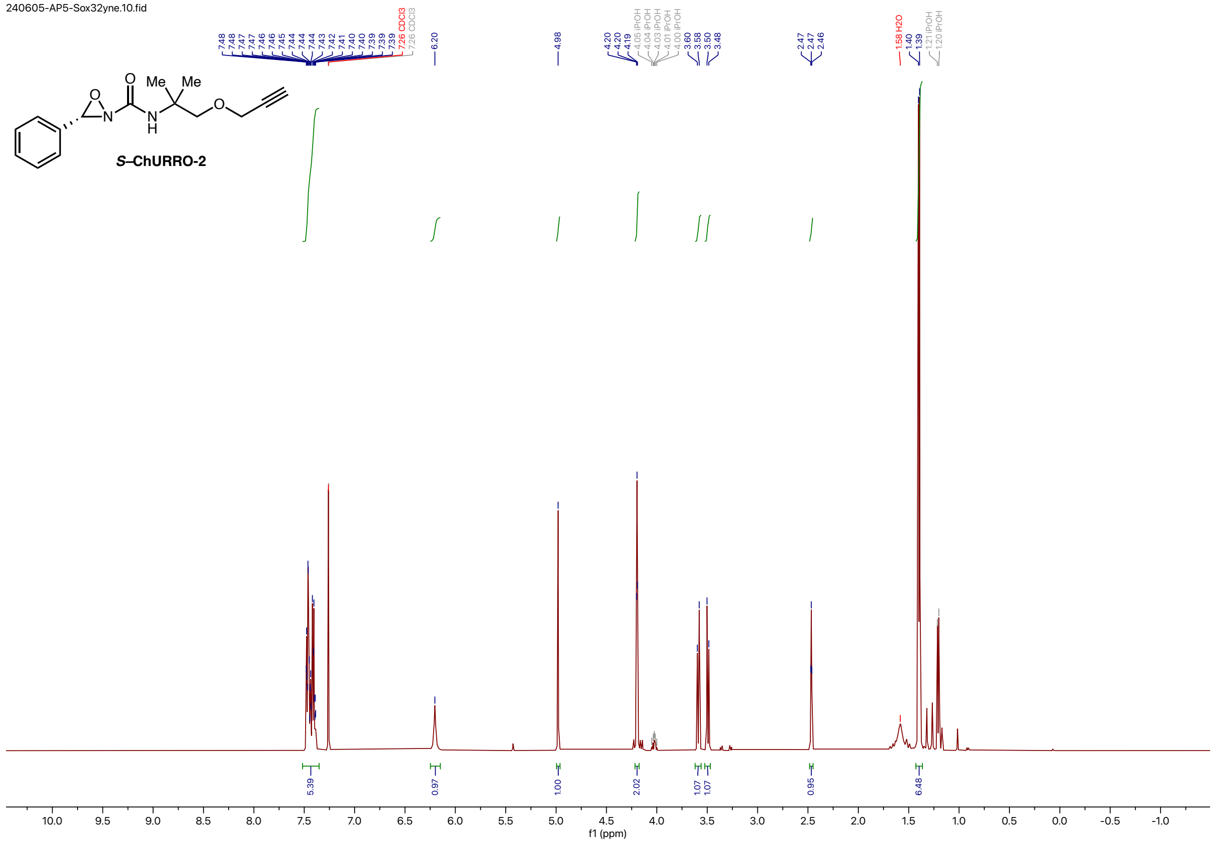
240605-AP5-Sox32yne.10.fid
Me
Me
O
N
H
S–ChURRO-2
5.39
0.97
1.00
2.02
1.07
1.07
0.95
6.48
f1 (ppm)

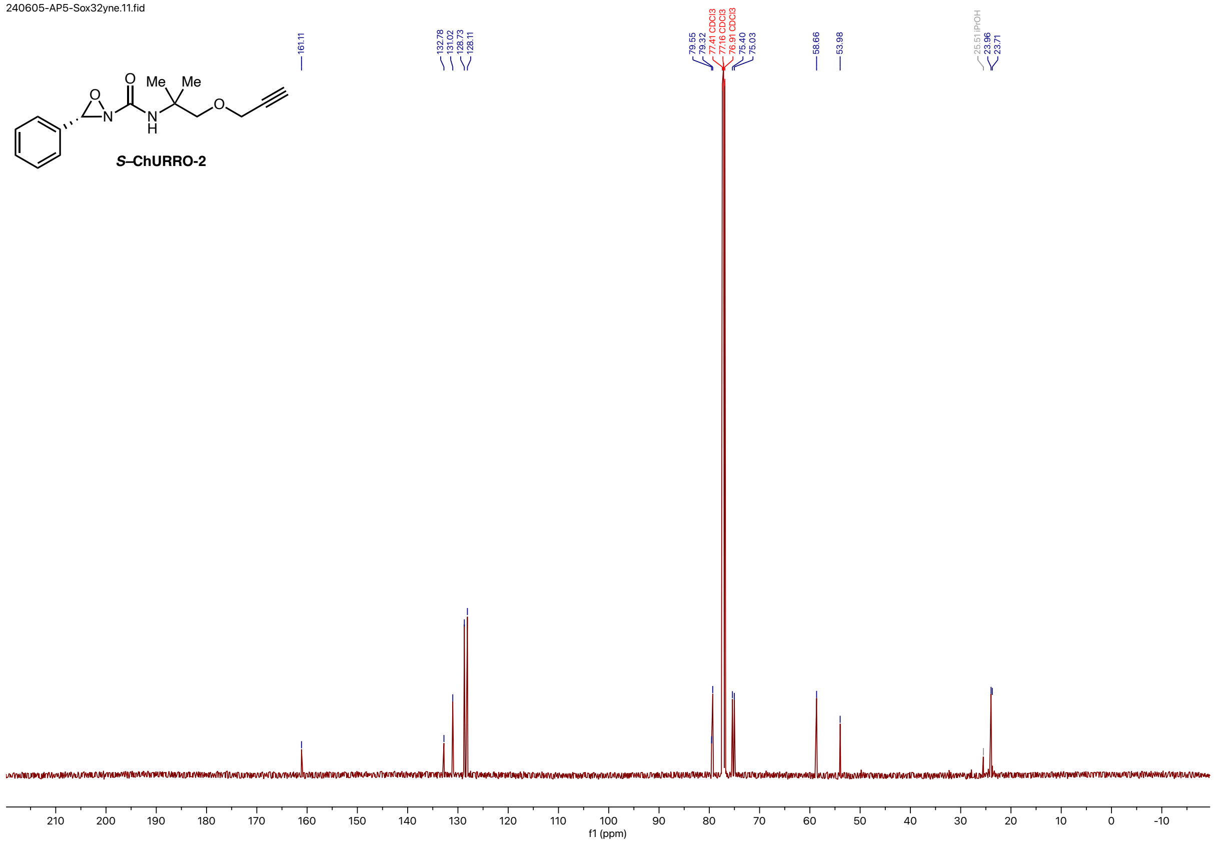
240605-AP5-Sox32yne.11.fid
161.11
132.78
131.02
128.73
128.11
79.55
79.32
77.41 CDCl3
77.16 CDCl3
76.91 CDCl3
75.40
75.03
58.66
53.98
25.51 iPrOH
23.96
23.71
Me
Me
O
N
O
N
H
O
S–ChURRO-2
210
200
190
180
170
160
150
140
130
120
110
100
90
80
70
60
50
40
30
20
10
0
-10
f1 (ppm)

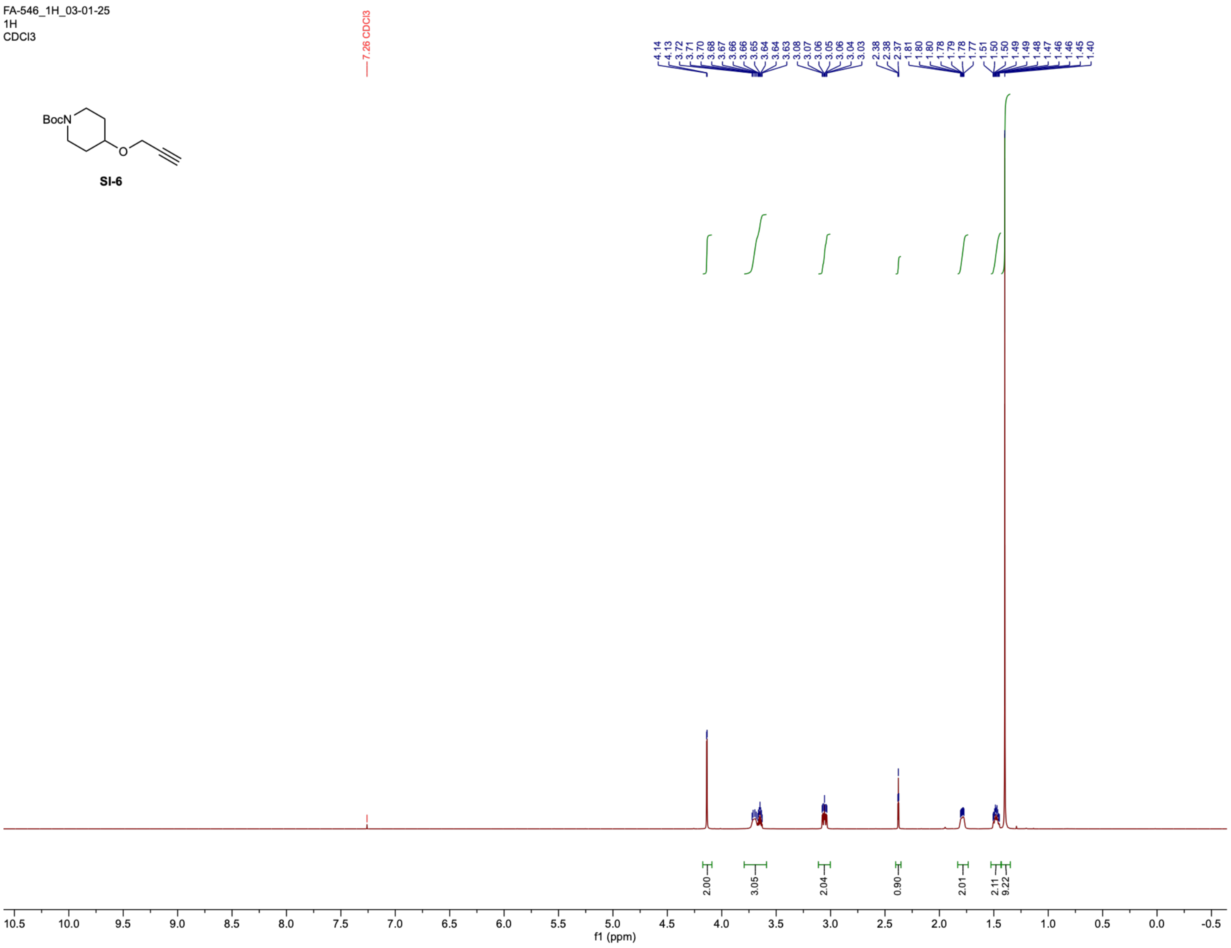

FA-546_1H_03-01-25
1H
CDCl3
7.26 CDCl3
4.14 4.13 3.72 3.71 3.70 3.68 3.67 3.66 3.66 3.65 3.64 3.64 3.63 3.08 3.07 3.06 3.05 3.06 3.04 3.03 2.38 2.38 2.37 1.81 1.80 1.80 1.78 1.79 1.78 1.77 1.51 1.50 1.50 1.49 1.48 1.47 1.46 1.46 1.45 1.40
BocN
O
SI-6
2.00 3.05 2.04 0.90 2.01 2.11 9.22
10.5 10.0 9.5 9.0 8.5 8.0 7.5 7.0 6.5 6.0 5.5 5.0 4.5 4.0 3.5 3.0 2.5 2.0 1.5 1.0 0.5 0.0 -0.5
f1 (ppm)

1328

FA-546_13C_03-01-25
13C
CDCl3

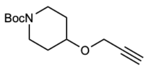


**SI-6**

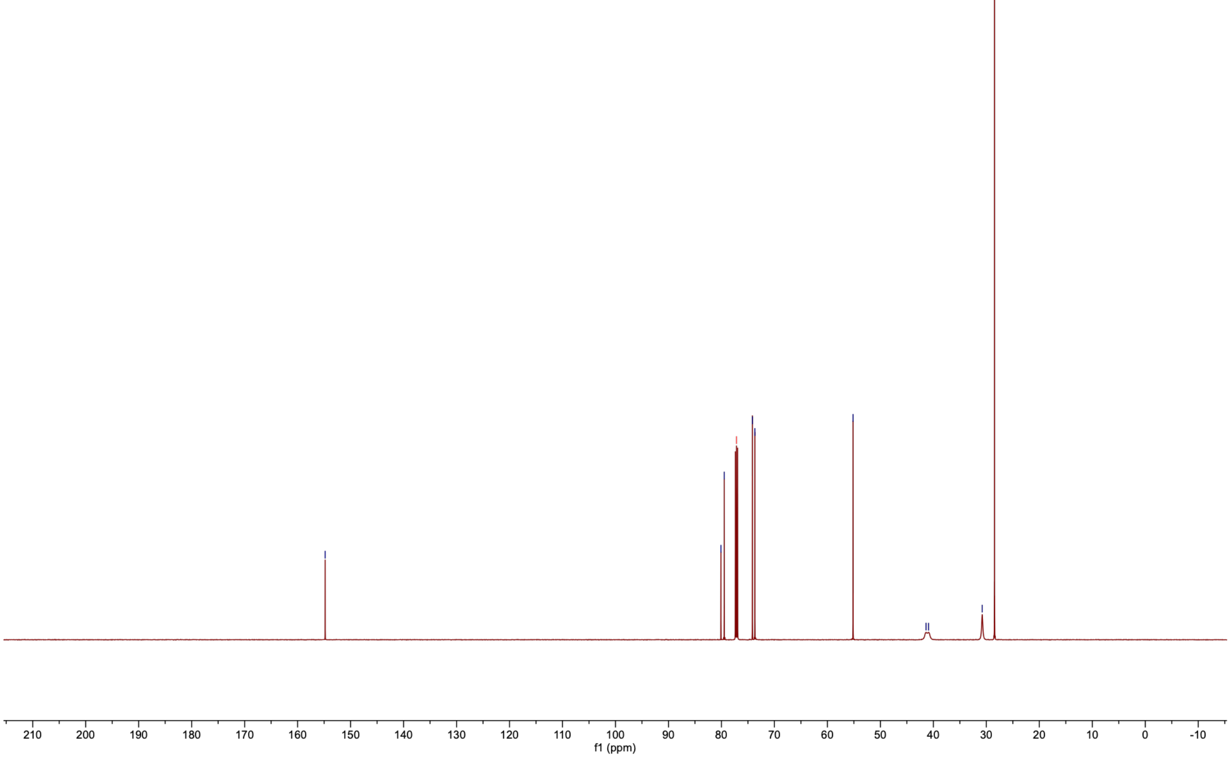

1329

FA-547_E446_03-01-25
1H
MeOD

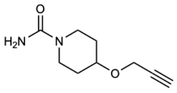


**SI-8**

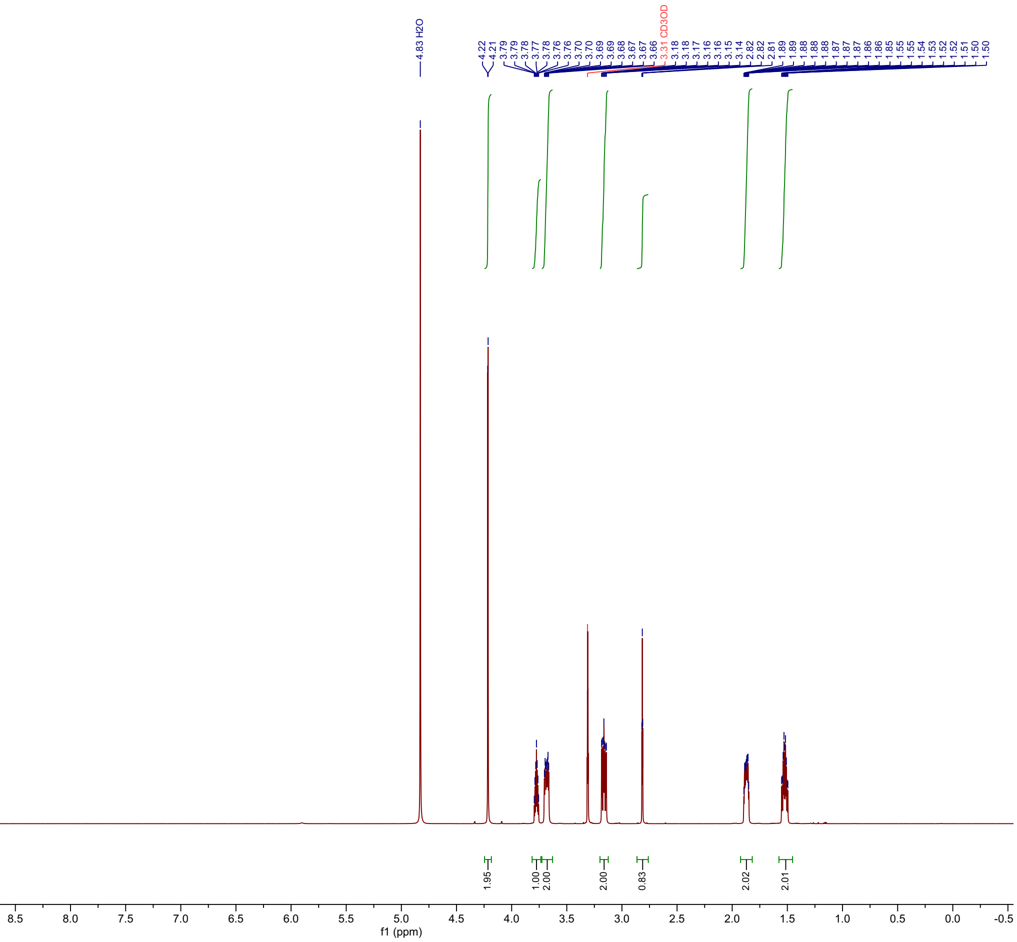

1330 FA-547_E446_13C_03-01-25
13C
MeOD

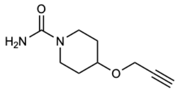


**SI-8**

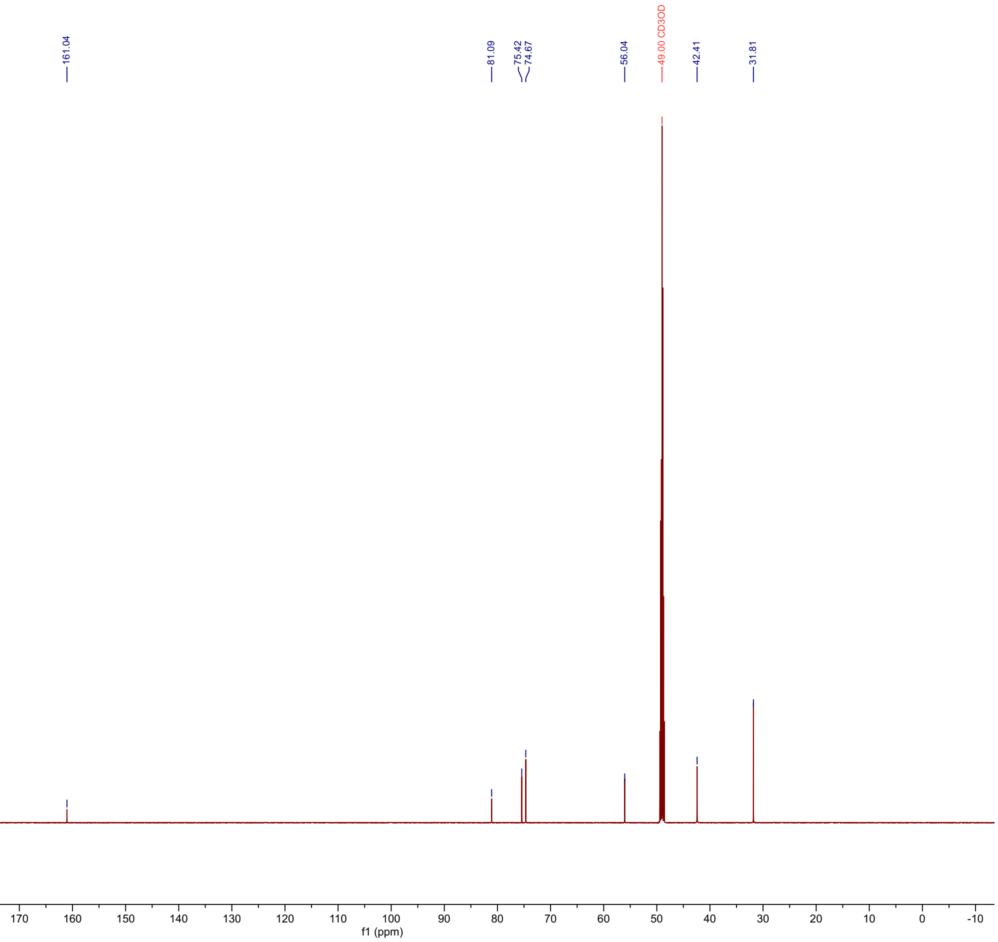

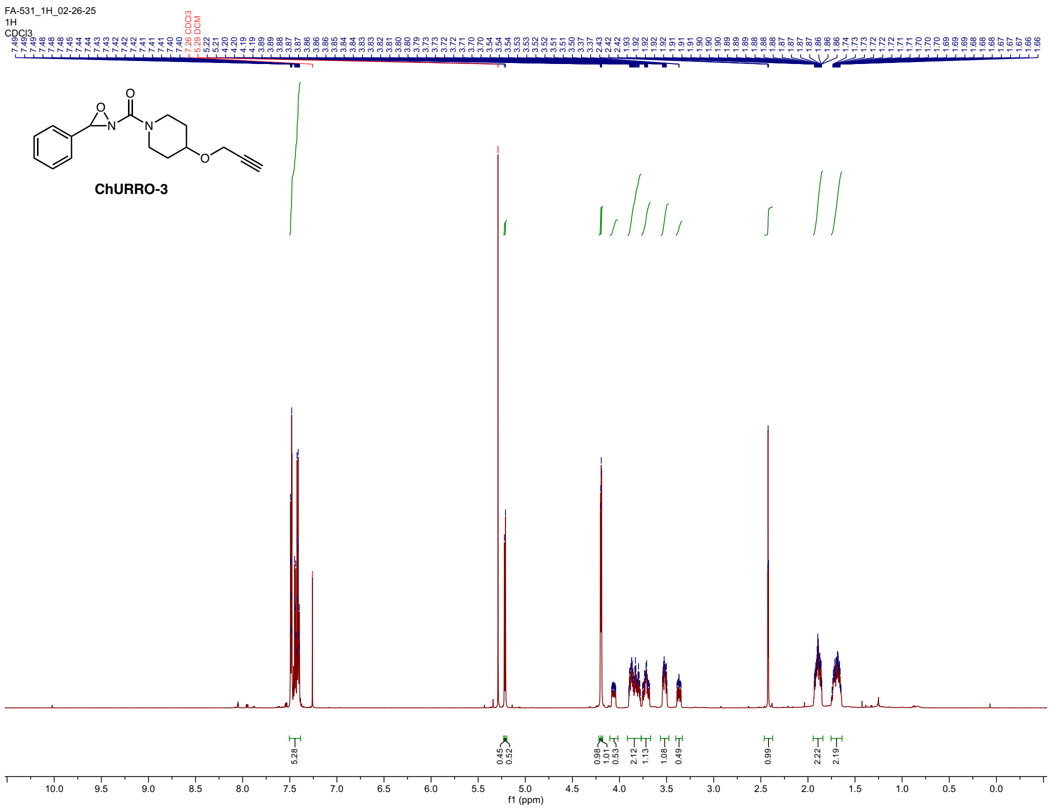


1331

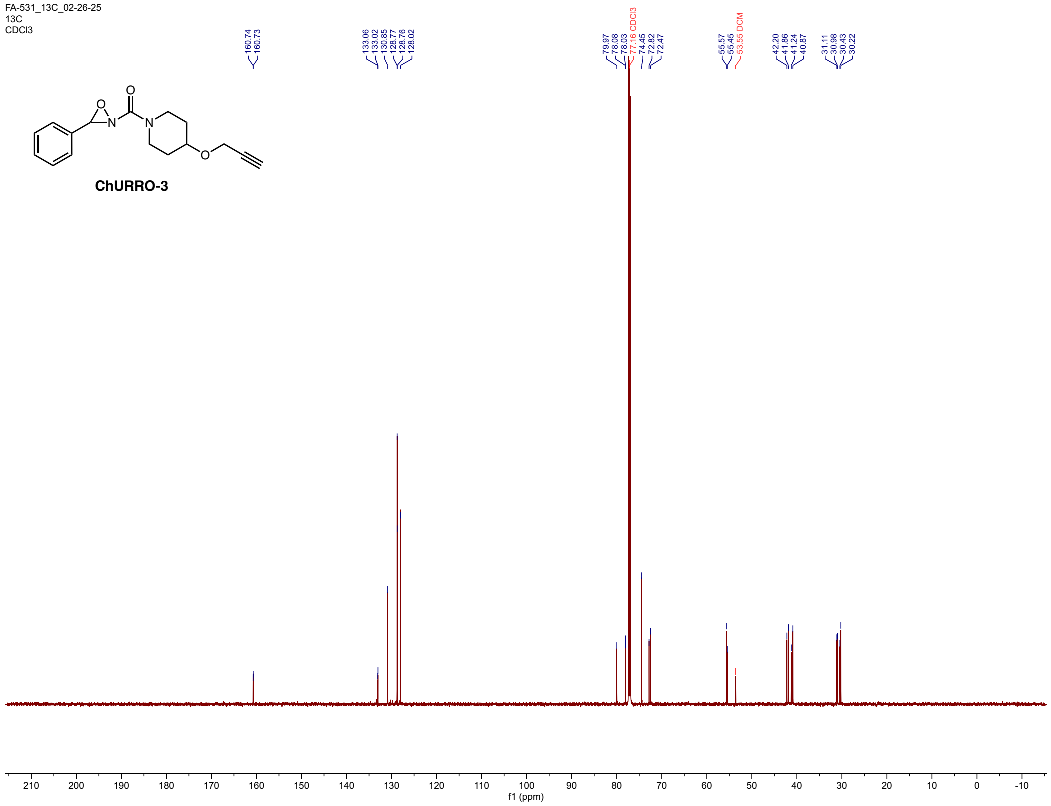
FA-531_13C_02-26-25
13C
CDCl3
ChURRO-3
160.74
160.73
133.06
133.02
130.85
128.77
128.76
128.02
79.97
78.08
78.03
77.16 CDCl3
74.45
72.82
72.47
55.57
55.45
53.35 DCM
42.20
41.86
41.24
40.87
31.11
30.98
30.43
30.22
210 200 190 180 170 160 150 140 130 120 110 100 90 80 70 60 50 40 30 20 10 0 -10
f1 (ppm)

1332

1333

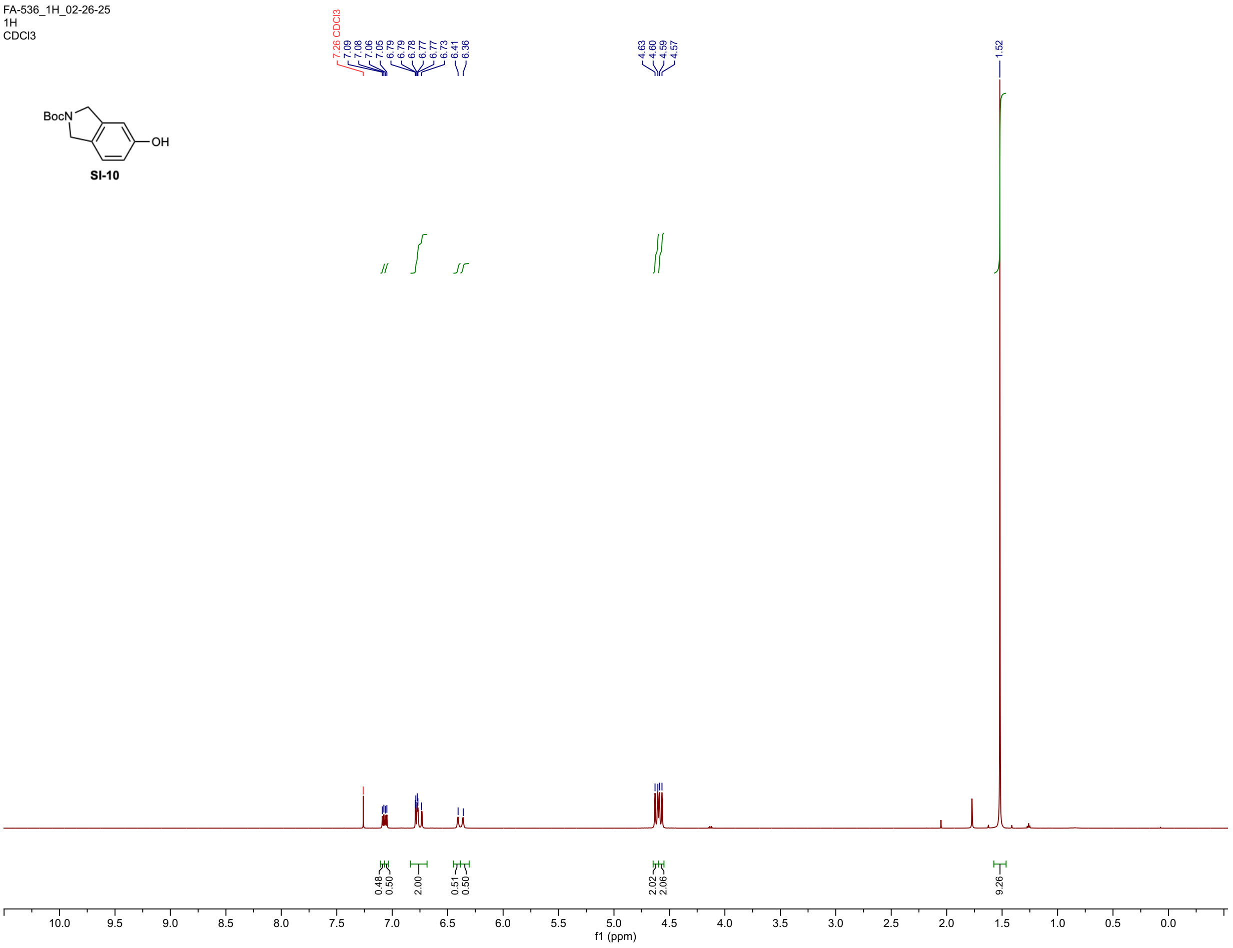
FA-536_1H_02-26-25
1H
CDCl3
BocN
OH
SI-10
7.26 CDCl3
7.09
7.08
7.06
7.05
6.79
6.79
6.78
6.77
6.77
6.73
6.41
6.36
4.63
4.60
4.59
4.57
1.52
0.48
0.50
2.00
0.51
0.50
2.02
2.06
9.26
10.0
9.5
9.0
8.5
8.0
7.5
7.0
6.5
6.0
5.5
5.0
4.5
4.0
3.5
3.0
2.5
2.0
1.5
1.0
0.5
0.0
f1 (ppm)

1334

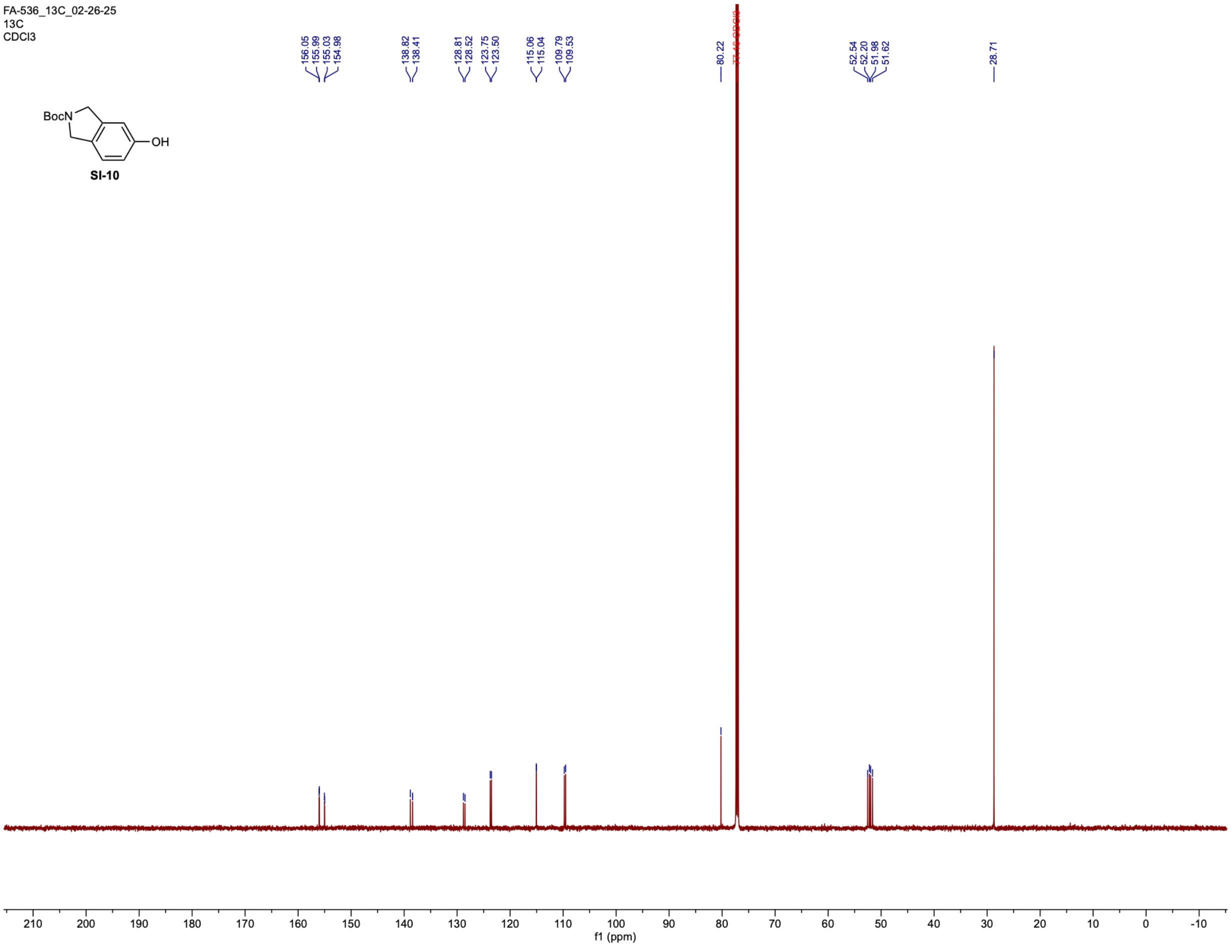

FA-536_13C_02-26-25
13C
CDCl3
156.05
155.99
155.03
154.98
138.82
138.41
128.81
128.52
123.75
123.50
115.06
115.04
109.79
109.53
80.22
52.54
52.20
51.98
51.62
28.71
210 200 190 180 170 160 150 140 130 120 110 100 90 80 70 60 50 40 30 20 10 0 -10
f1 (ppm)
BocN
OH


**SI-10**

1335

FA-537_1H_03-01-25
1H
CDCl3

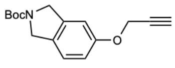
SI-11

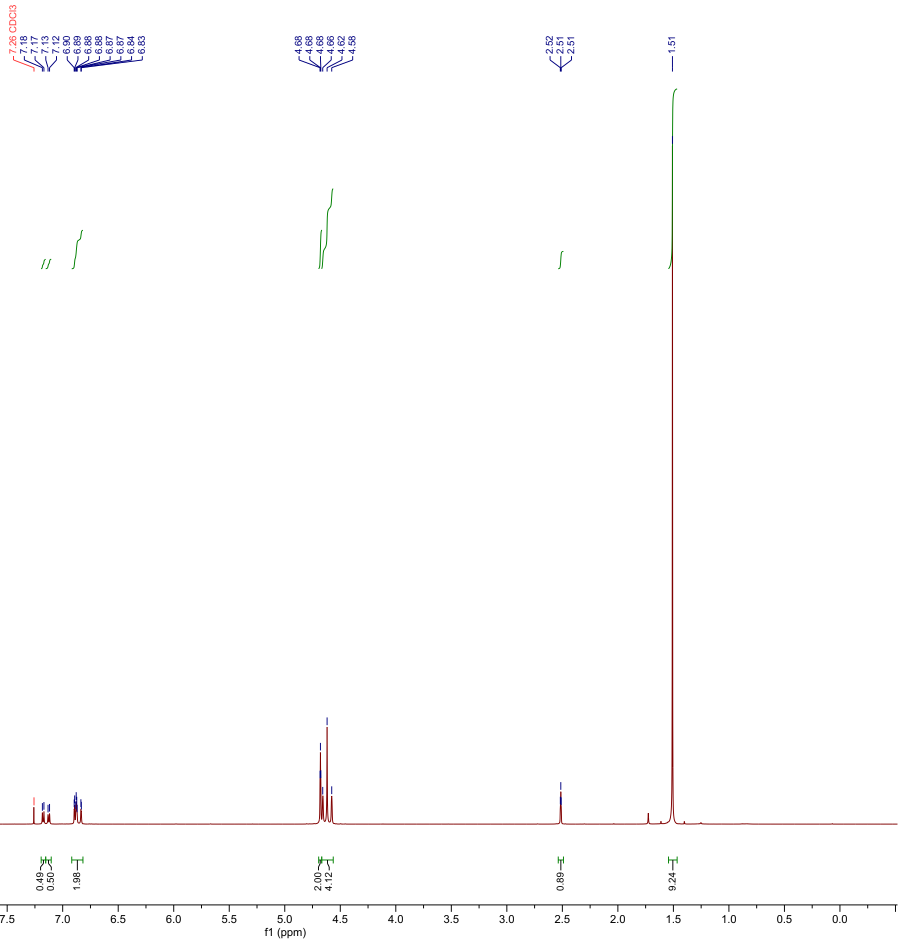

1336

FA-537_13C_03-01-25
13C
CDCl3

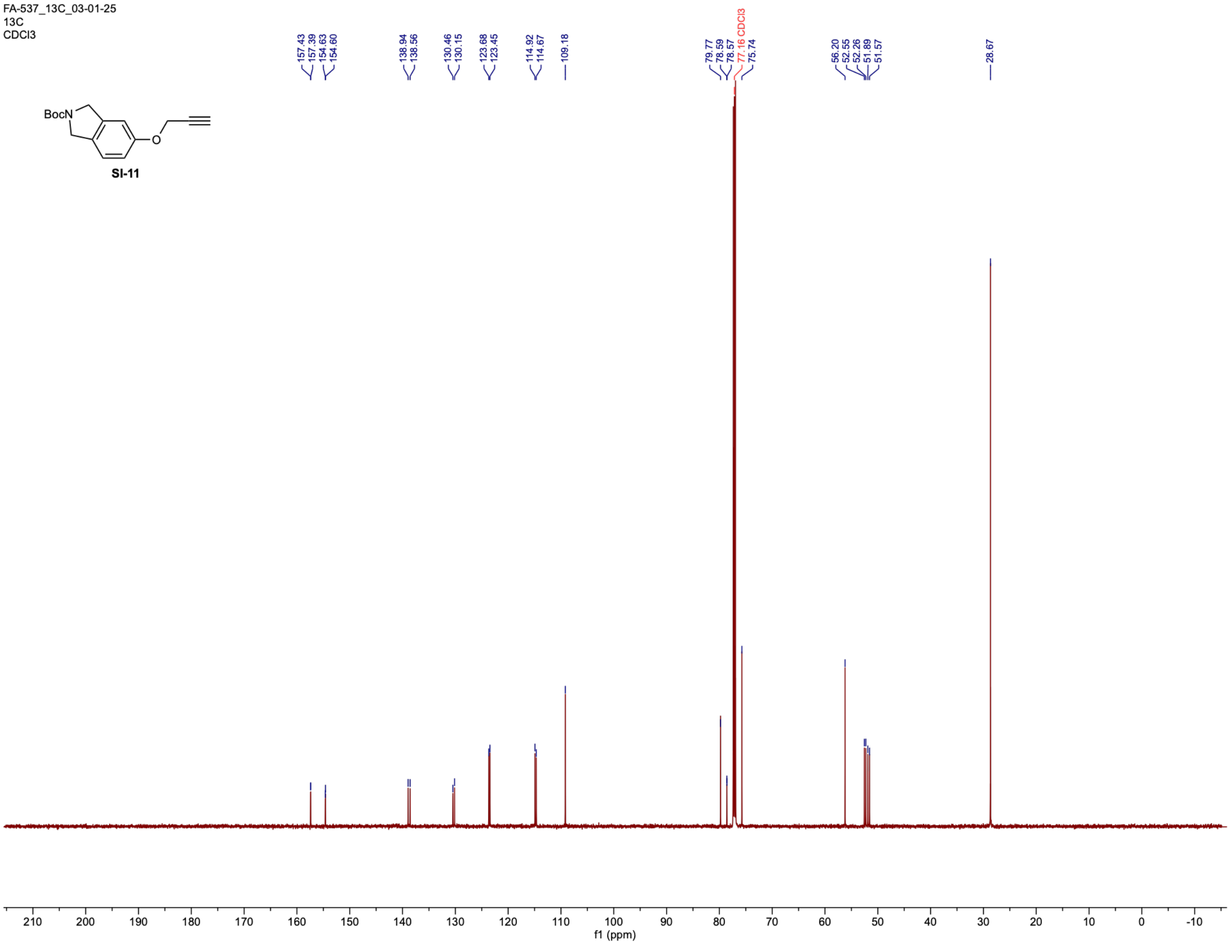

1337

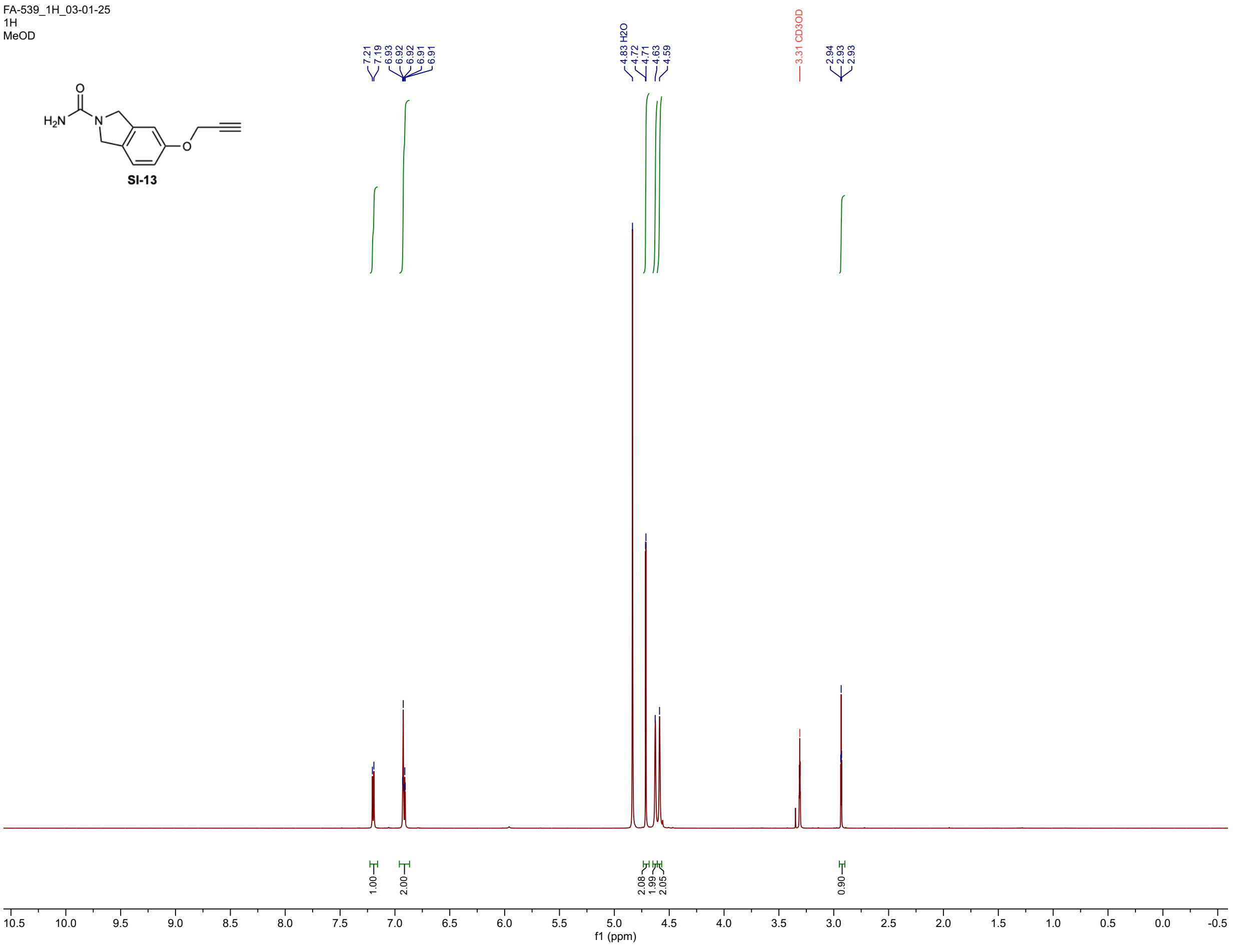
FA-539_1H_03-01-25
1H
MeOD
SI-13
7.21
7.19
6.93
6.92
6.92
6.91
6.91
4.83 H2O
4.72
4.71
4.63
4.59
3.31 CD3OD
2.94
2.93
2.93
1.00
2.00
2.08
1.99
2.05
0.90
f1 (ppm)

1338

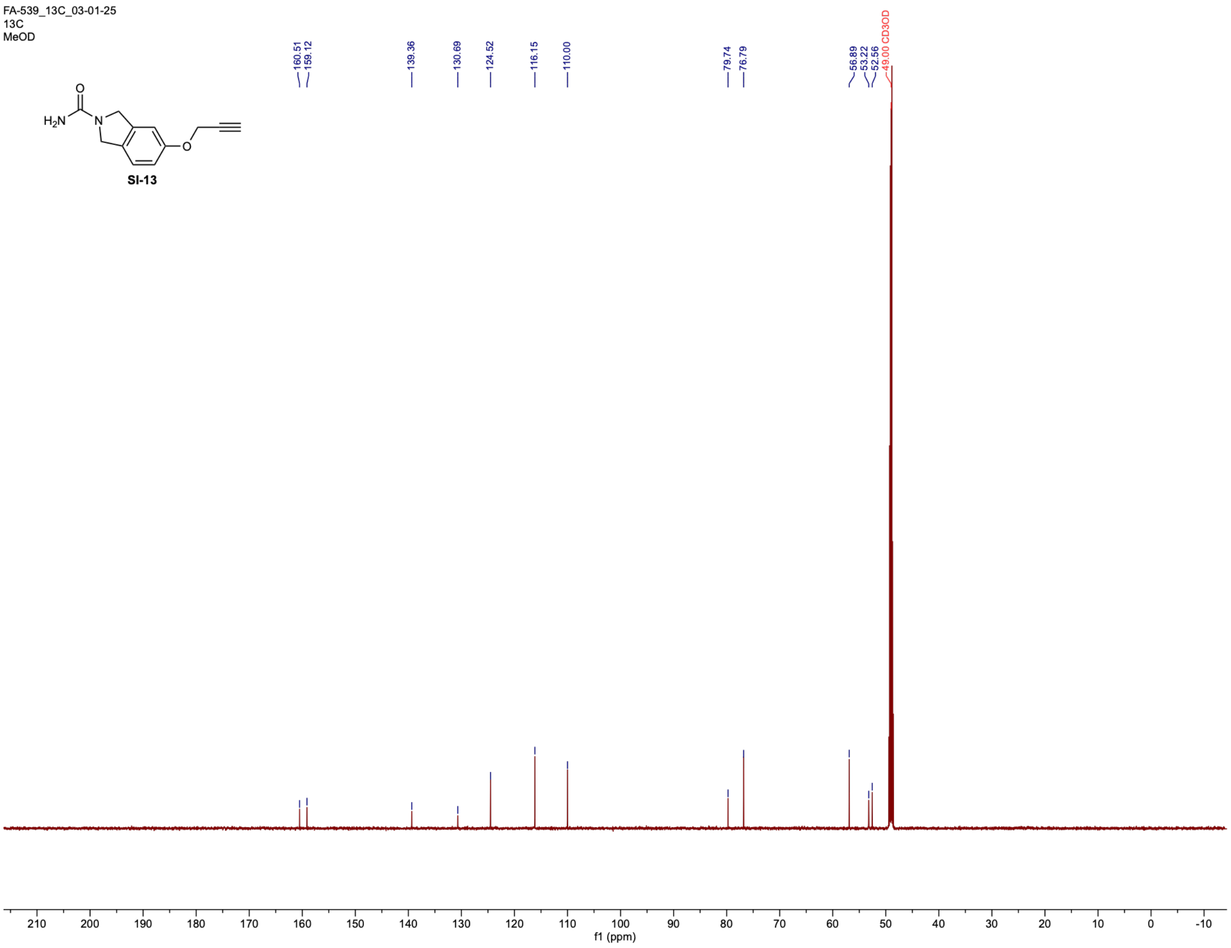

FA-539_13C_03-01-25
13C
MeOD
160.51
159.12
139.36
130.69
124.52
116.15
110.00
79.74
76.79
56.89
53.22
52.56
49.00 CD3OD
SI-13
210
200
190
180
170
160
150
140
130
120
110
100
90
80
70
60
50
40
30
20
10
0
-10
f1 (ppm)

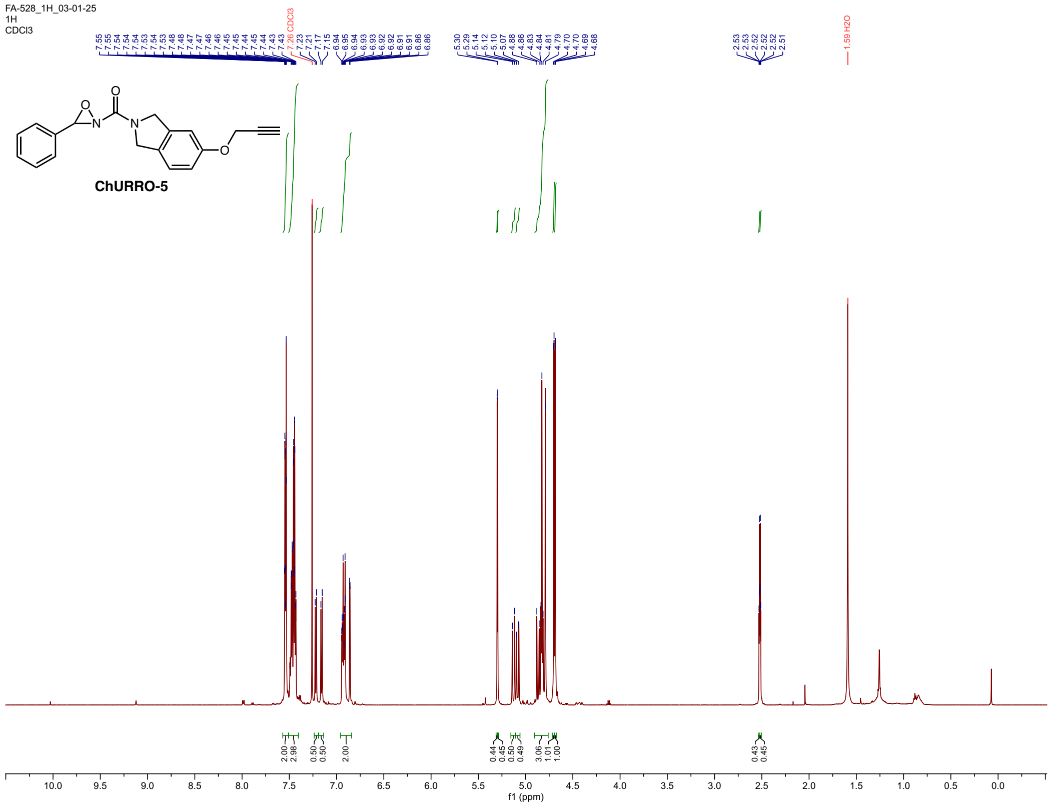
FA-528_1H_03-01-25
1H
CDCl3
7.55 7.55 7.54 7.54 7.54 7.53 7.53 7.54 7.48 7.48 7.47 7.47 7.46 7.46 7.45 7.45 7.44 7.45 7.44 7.43 7.43
7.26 CDCl3
7.23 7.21 7.17 7.15 6.94 6.95 6.94 6.93 6.93 6.92 6.92 6.91 6.91 6.86 6.86
5.30 5.29 5.14 5.12 5.10 5.07 4.88 4.86 4.83 4.84 4.81 4.79 4.70 4.70 4.69 4.68
2.53 2.53 2.52 2.52 2.52 2.51
1.59 H2O
ChURRO-5
2.00 2.98 0.50 0.50 2.00
0.44 0.45 0.50 0.49 3.06 1.01 1.00
0.43 0.45
10.0 9.5 9.0 8.5 8.0 7.5 7.0 6.5 6.0 5.5 5.0 4.5 4.0 3.5 3.0 2.5 2.0 1.5 1.0 0.5 0.0
f1 (ppm)

1339

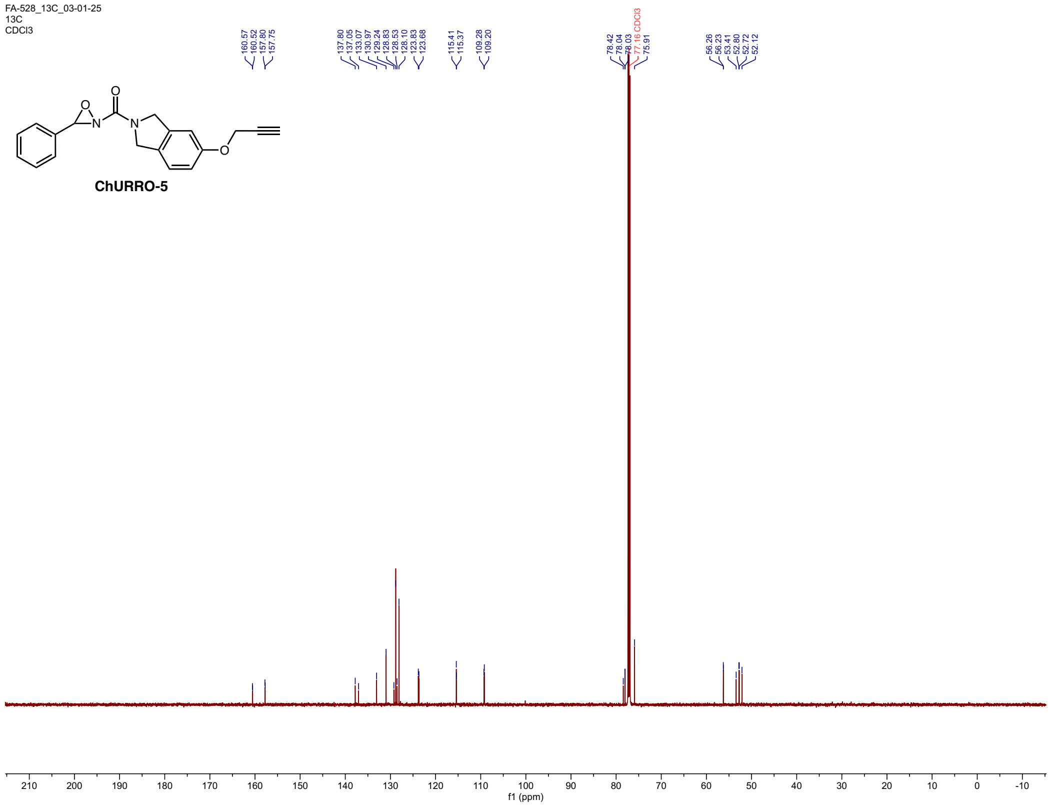


1340

1341

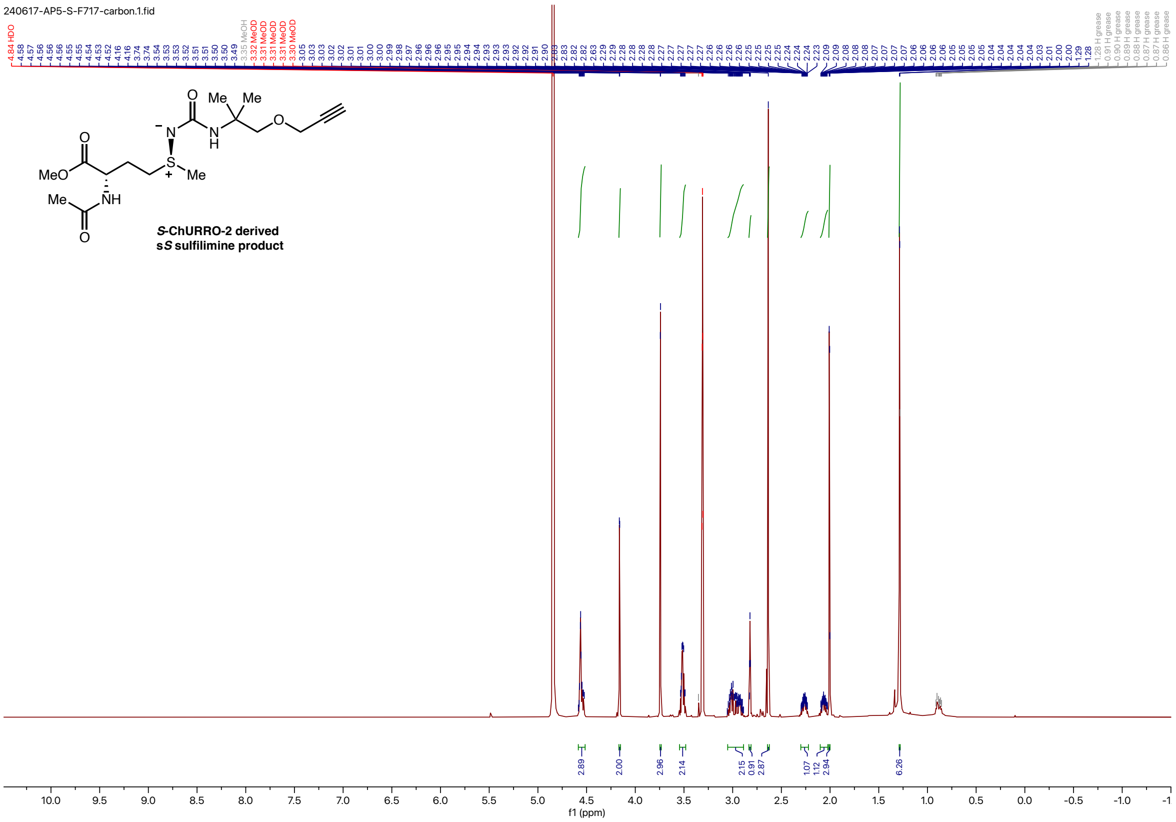


1342
1343

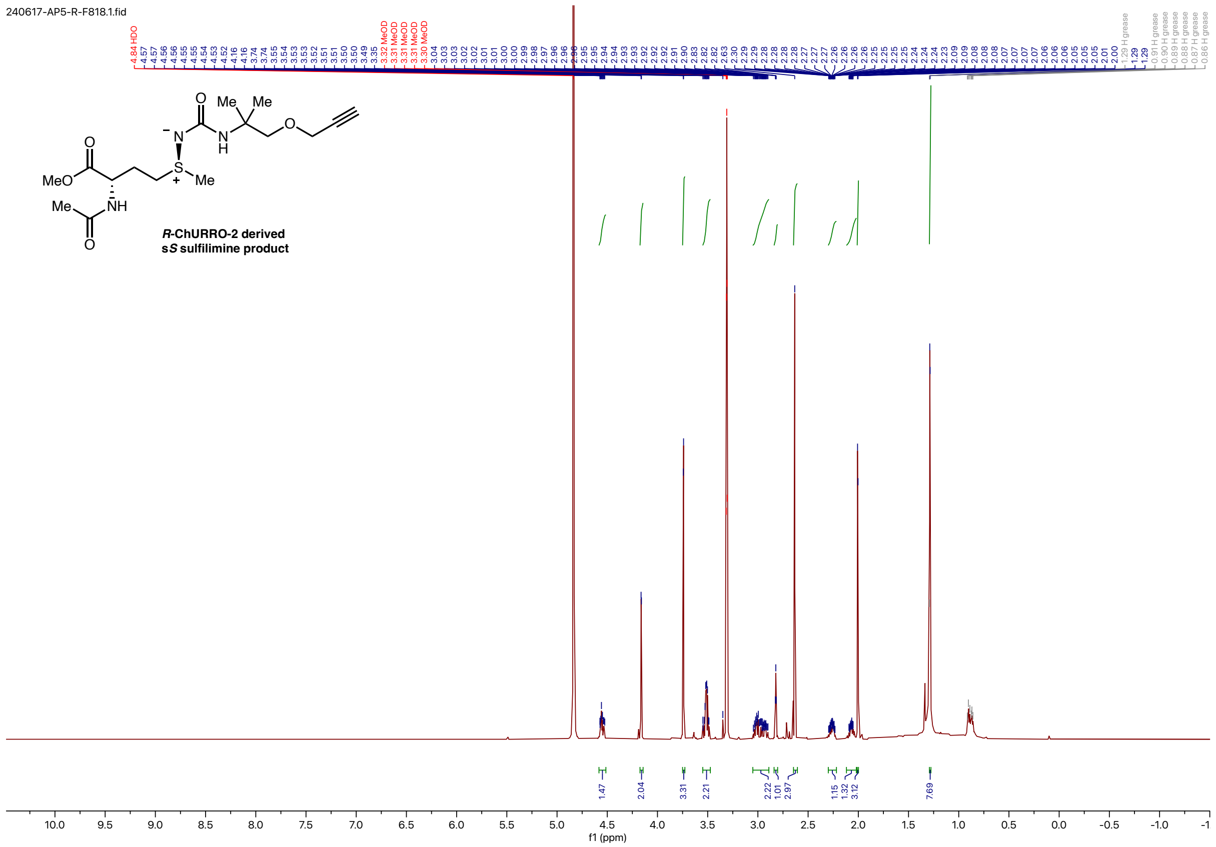
240617-AP5-R-F818.1.fid
*R*-ChURRO-2 derived
s*S* sulfilimine product
f1 (ppm)

1344
